# Lung Adenocarcinoma Just Desserts: An Expanding Pie of Activating Oncogenes or a Layer Cake of Integrated Alterations

**DOI:** 10.1101/2025.09.19.677365

**Authors:** Minjeong Kim, Wisut Lamlertthon, Heejoon Jo, Yan Cui, Hyo Young Choi, Katherine A. Hoadley, Matthew P. Smeltzer, Michele C. Hayward, Matthew D. Wilkerson, Liza Makowski, D. Neil Hayes

## Abstract

The molecular landscape of lung adenocarcinoma (LUAD) is often summarized as a “pie chart” of driver oncogenes, suggesting identification and targeting of oncogenic drivers is the best clinical approach. However, this model oversimplifies LUAD biology. Patients with identical RAS or RAF mutations exhibit heterogeneous signaling influenced by co-mutations, transcriptional programs, and lineage context. We propose a hierarchical framework integrating oncogenes within cellular context by examining canonical *EGFR* mutations. We defined an *EGFR* mutation signature (mSig) by identifying differentially expressed genes in *EGFR*-mutant LUADs. Semi-supervised clustering and machine learning models were used to test performance and reproducibility across independent datasets. We analyzed molecular subtypes, lineage markers, co-occurring mutations, and candidate drug targets in EGFR-mSig positive (+) versus negative (-) tumors. The EGFR mSig showed robust classification performance across datasets (AUROC = 0.83–0.95; mean NPV = 96.3%). Validated unsupervised gene expression subtypes and lung lineage markers were closely aligned with EGFR mSig status. mSig(+) tumors were identified, even in tumors without *EGFR* mutations. A subset of canonical RAS mutations mirrored the EGFR mutation pattern. *EGFR*-mutant/mSig(–) tumors were enriched for non-Bronchioid subtypes and had co-mutations in *TP53* or RAS. Coordinated mutations were identified including RAS, *KEAP1*, *STK11*, *TP53*, and *CDKN2A*, supportive of prior reports. In sum, novel EGFR mSig that captures the transcriptional footprint of EGFR activation, revealed a subset of EGFR wildtype LUADs with “mutant-like” features. Lineage-informed classification highlights subtype-dependent oncogene activity and supports new therapeutic strategies. A context-specific association between RAS mutation and expression of the dual-specificity phosphatase gene DUSP4 may have therapeutic potential.

## Introduction

More than 20 years ago, multiple investigators first reported frequent mutations in the epidermal growth factor receptor gene, *EGFR*, in association with therapeutic responses to selective kinase inhibitors (1). Progress in targeting EGFR inspired exhaustive efforts to identify additional oncogene targets with similar druggable potential, with numerous successes including activating mutations of *BRAF*, *MET*, *FGFR*, *RET*, fusions of the genes *ROS1* and *ALK*, as well as others (2). The precise fraction of lung adenocarcinoma (LUAD) with therapeutic targets remains an area of active research, as does the set of variants that are responsive to pharmacologic agents (3, 4). However, the pace of discovery has slowed in recent years despite many patients remain in the unsatisfying category of “druggable oncogene negative.” Likewise, the therapeutic potential of some variants remains to be fully realized. For example, pharmacologic targeting of the most common driver, KRAS, has recently achieved notable success, representing a major step forward while continued efforts aim to expand efficacy.(5, 6). Failure to target oncogenic drivers by direct inhibition has led investigators to consider indirect strategies including blockade of pathway-related signaling components. Downstream pathway blockade has been successful on occasion, such as MEK inhibition in some *BRAF*-mutant (mt) tumors (7). However, in most cases, attempts to inhibit downstream signaling elements of activated oncogenes have yielded limited clinical benefit.

Partial explanations for the failure of downstream blockade have emerged in selected cases. For example, *BRAF*-mt colon cancers are refractory to BRAF inhibitors due to compensatory upregulation of alternative pathway components (8). In contrast, for *KRAS*-mt LUAD, the mechanisms explaining failure of downstream inhibition appear to include heterogeneity in baseline signaling even in the absence of pharmacologic inhibition (9). Identical *KRAS* sequence variants have alternative downstream signaling targets as a function of co-mutated genetic elements as well as the molecular context of expressed lineage transcription factors (10, 11). At least three well-established signaling subtypes of *KRAS* have been suggested as a function of the lineage marker *NKX2-1* (encoding the protein TTF1), mutations in *TP53*, *STK11*, *KEAP1*, and other factors (12–14). While mostly untested, the hypothesis suggests that different groups might have different therapeutic options including downstream blockade in cases where the correct KRAS class and downstream targets were characterized.

Recognizing the slowing pace of therapeutic target identification in LUAD, as well as the data suggesting that multiple RAS-pathway elements in LUAD appear to have heterogenous signaling alternative pathways (*BRAF*, *KRAS*), we hypothesized that the gene expression profile of patients with mutated EGFR might provide insights into the signal status of other known and unknown downstream elements of RAS signaling. Based on the *BRAF* and *KRAS* experience, we also considered that the molecular context, such as previously validated gene expression subtypes (15–18), co-mutated oncogenes (such as *TP53*), and transcription factors, might influence signaling networks, therapeutic targets, and even unmask aspects of tumorigenesis.

## Materials and Methods

### Cohort Data Acquisition

For the purposes of generating expression signatures of EGFR activation, primary gene expression data was obtained from multiple independent sources which included both RNA characterization, DNA mutation data (EGFR mutation at a minimum), and clinical characteristics (16, 19). Cohort eligibility included the requirement that patients were primarily selected as part of an initial diagnosis and were previously untreated. Only validated kinase-activating mutations in exons 18-21 of *EGFR* were considered as mutated for the purpose of the study. To support the generalizability of expression signatures, we obtained RNA derived from three complimentary technology platforms: Illumina brand one-color gene expression arrays (Memorial-Sloan Kettering Cancer Center (MSKCC, n=192, GSE68465) and Tumor Sequencing Project (TSP, n=41, GSE12667), Agilent brand two-color gene expression arrays (University of North Carolina at Chapel Hill (UNC, n=73, GSE36471), and a cohort based on next generation mRNA-seq protocol (The Cancer Genome Atlas (TCGA, n=486, Genomic Data Commons (GDC) Portal) (20). The MSKCC cohort was selected as the training dataset and all other cohorts were withheld as independent validation cohorts. For additional integrated analysis, we considered copy number and additional mutation data from the MSKCC, TSP, and TCGA datasets. For integrative analysis, mutation calls and copy number of selected driver oncogenes including *EGFR*, *TP53*, *CDKN2A*, *KEAP1*, *STK11*, *KRAS*, *NRAS*, *HRAS*, *BRAF*, *HER2*, *ALK*, *MET*, and *ROS1* were obtained from the prior published datasets (21). This study was reviewed by the University of Tennessee Health Science Center (UTHSC) Institutional Review Board (IRB) 25-10419-NHSR.

### RNA Preprocessing

Pipelines for RNA processing and normalization have been previously reported (16, 17). Briefly, Agilent 2-color gene expression array assays were background corrected and normalized according to standard protocols of the platforms. Specifically, for Agilent 2-color microarray MSKCC and UNC data, Lowess normalization was applied to obtain expression values. For Affymetrix 1-colored microarray TSP data, the Robust Multi-array Average (RMA) algorithm was applied for background correction and normalization. Each dataset was then subjected to median centering. For TCGA LUAD mRNA-seq data, RNA-seq by Expectation-Maximization(RSEM)-normalized expression data (Fragments Per Kilobase of transcript per Million mapped reads) was obtained (22). All datasets were z-score normalized before visualization in heatmaps in R version 4.3.1 (23). For all other analyses, quantile normalization was applied via preprocessCore R package 1.64.0 (24).

### Establishment of EGFR Mutation Signature (mSig)

To investigate the role of gene expression in predicting *EGFR* mutation, we calculated the delta score for each gene in the supervised testing *EGFR*-mt tumors versus *EGFR* wild-type (WT) tumors using the training cohort (MSKCC) by the Significance Analysis of Microarray R package (SamR, version 3.0) (25). In this manner genes could be ranked for positive and negative correlation with mutation status. In parallel, we generated a family of centroid-based predictors for mt versus WT using the Classification to the Nearest Centroids (ClaNC) method (26, 27).

To visualize expression profiles of each molecular subtype, semi-supervised clustering was used and visualized via customized heatmap function based on gplots R package (version 3.1.3.1)(28) and complexHeatmap R package (version 2.18.0) (29, 30). We applied 6 machine learning (ML) algorithms for supervised *EGFR* mutant prediction - Support Vector Machines (SVM), with different kernel functions, including Linear, Polynomial, and Radial Basis Function (RBF) and Ensemble Learning algorithms - Random Forest (RF), AdaBoost, and LogitBoost. The top 1,000 genes were selected using *t*-test and were used as input features for ML. We used the receiver operating characteristic (ROC) curve based on 10-fold cross-validation to evaluate the performance of ML models. The area under the ROC curve (AUC) values were calculated and used as the primary performance metric. Other performance metrics including sensitivity, specificity, accuracy, positive predictive value (PPV) and negative predictive value (NPV) were also calculated. For external validation (or independent testing), the ML performances were further tested using different combinations of training and testing sets.

For the purpose of attributing biologic significance to lists of genes developed in the study, we used “Database for Annotation, Visualization and Integrated Discovery” (DAVID) and gene set enrichment analysis (GSEA) (version 4.3.2) (31–33). Gene lists derived from DAVID were documented in terms of enrichment for functional gene groups by biological pathways and statistical significance (p-value < 0.05 or highly enriched Fold Enrichment >2) was assessed for the purpose of visualization using the dplyr R package (version 1.1.3) (34) and ggplot2 R package (version 3.5.1) (35).

### Prediction Performance Test

In prediction performance tests, *EGFR*-mt/mSig(+) patients were classified as true positives (TP), *EGFR* WT/mSig(-) patients as true negatives (TN), *EGFR* WT/mSig(+) patients as false positives (FP), and *EGFR*-mt/mSig(-) patients as false negatives (FN). The performance test was calculated in R (version 4.3.1.) (23) and visualized via caret R package (version 6.0.94) (36).

### Statistical Analysis

In patient demographic data, molecular subtype, stage at diagnosis, and smoking history were tested by Chi-squared test. Pack-years were tested by ANOVA. Overall survival data from the MSKCC cohort was summarized with Kaplan-Meier survival curves. Overall survival was defined as the time from diagnosis of NSCLC to death from any causes or time of last follow-up within 36 months. Right censoring was applied to the last follow-up time point where no event (death) had occurred. Visualization of the curves and calculation of the log rank p-value was conducted via survival R package (version 3.7.0) (37, 38). To assess mutual exclusivity between two events, odds ratio (OR) was calculated and the *p*-values for mutual exclusivity were calculated by Fisher’s Exact test. OR and *p*-values were visualized in heatmaps via ComplexHeatmap (R package version 2.18.0) (29, 30). All data handling and statistical analyses were performed in the R version 4.3.1. environment (23).

### Data Availability

The data analyzed in this study were obtained from Gene Expression Omnibus (GEO) at GSE68465, GSE12667, GSE36471, and GDC Portal) (20).

## Results

### Patient Characteristics

A total of 792 patients with gene expression, mutation, and clinical data were available for analysis divided into training (n=192, MSKCC) and 2 validation cohorts (validation #1: n=114, combined UNC and TSP, and validation #2, n=486, TCGA; **Table 1**). Patients were assigned to gene expression subtypes using methods which have been previously reported (15, 16). Overall, the cohorts were similar in their composition with minor exceptions.

**Table 1.**
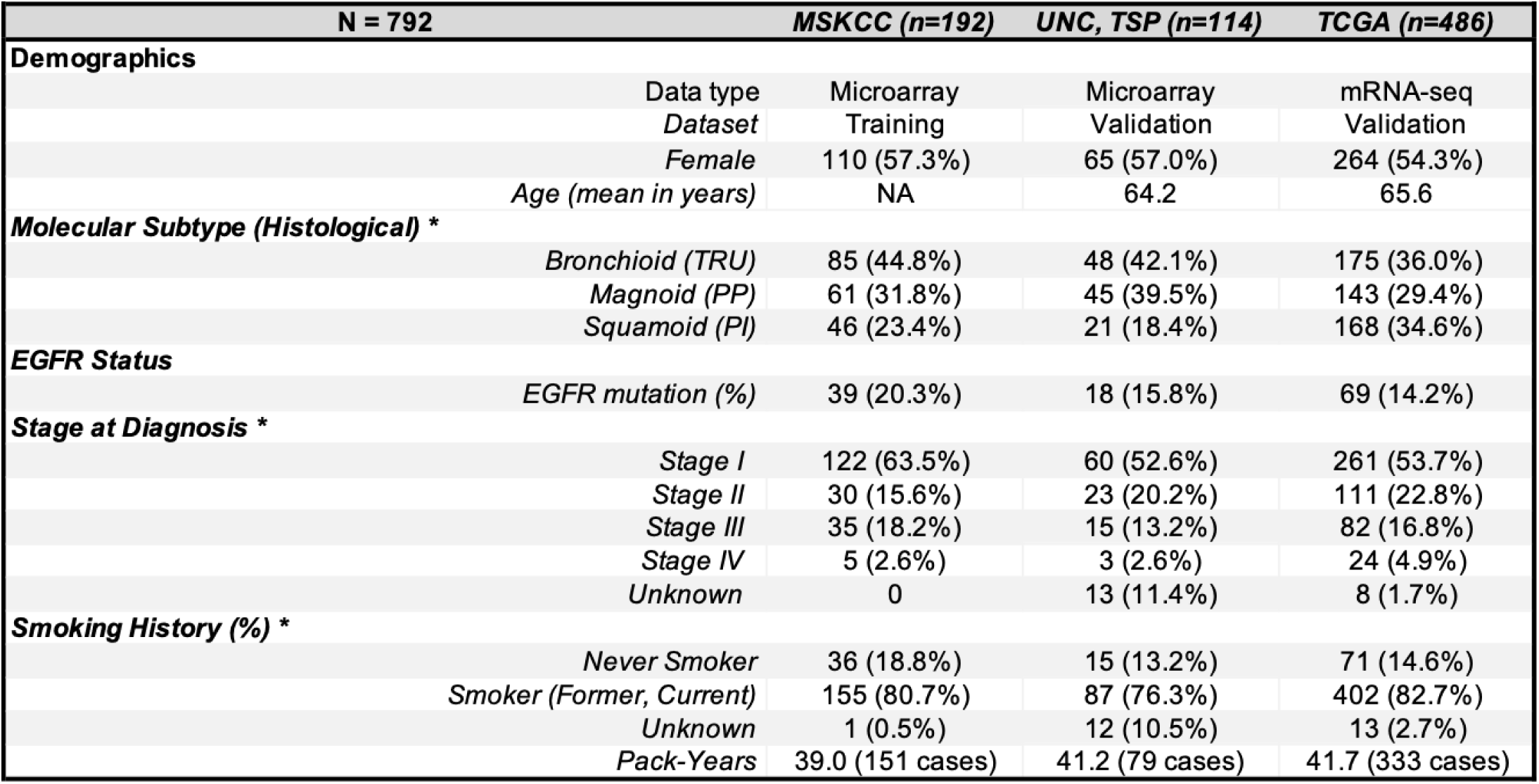
Clinical cohorts included in the study. TRU, Terminal Respiratory Unit, PP, Proximal Proliferative, PI, Proximal Inflammatory (* p-value < 0.001).

### EGFR Mutation Signature (mSig) Predicts EGFR-mutant-like LUAD Tumors

For the purpose of defining expression patterns associated with EGFR mutation activation, we constructed a series of supervised models predicting EGFR mutation status under a wide variety of conditions (**Figure 1**, Supplementary Figure S1-2). All models had very similar performance. From the training data set (MSKCC), a model with 1,020 DEGs was selected using the ClaNC technique (Supplementary Table S1; Supplementary Figure S2). A total of 690 genes were over-expressed and 330 genes were under-expressed in EGFR-mt tumors compared to EGFR WT. Varying the number of genes included in the model from a few dozen to several thousand had little impact on the performance of the ClaNC training performance. The choice of 1,020 genes was empiric and intended to be of sufficient size to allow other analyses to be performed such as gene set enrichment type studies.

**Figure 1.**
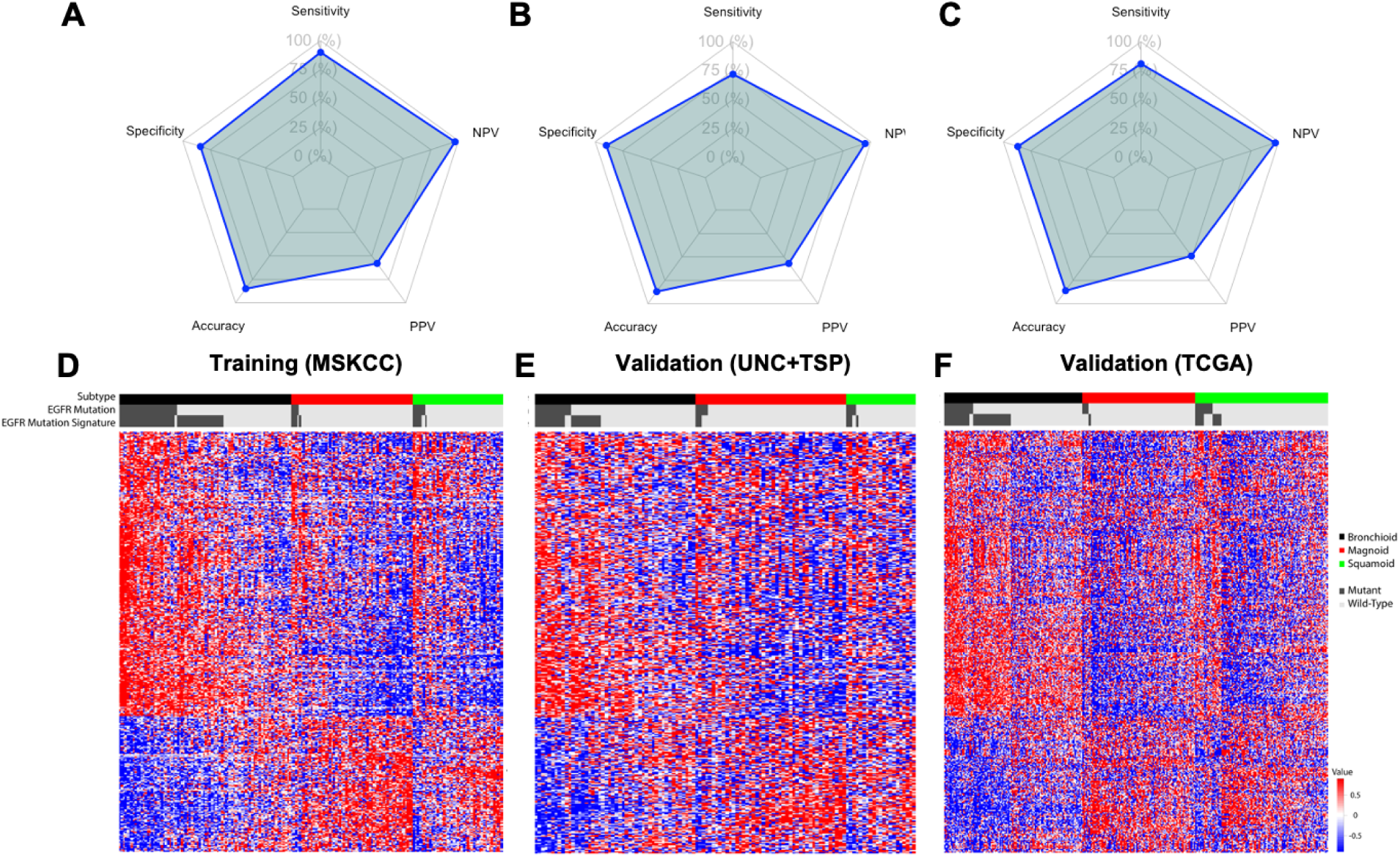
Performance and reproducibility of the EGFR mutation signature in LUAD. **A**–**C**. Radar plots showing performance metrics of the EGFR mutation signature across datasets. Metrics include positive predictive value (PPV) and negative predictive value (NPV). **D**–**F**. Heatmaps of gene expression profiles of LUAD patients stratified by EGFR mutation status. Rows represent 1,020 genes included in the EGFR mSig. Columns represent individual patients, clustered by LUAD molecular subtypes (Bronchioid-black; Magnoid-red; or Squamoid- green), with EGFR mutation and mSig status using a semi-supervised approach. Annotations indicate EGFR mutation status and EGFR mutation signature status with mutant (dark gray bars) or Wild-Type (light grey bars). **A** and **D**: training cohort (MSKCC); **B** and **E**: validation cohort 1 (UNC + TSP); **C** and **F**: validation cohort 2 (TCGA).

In training, sensitivity (90%), specificity (84%), and NPV (97%) were high, while PPV was only modest at 58%. Validation experiments showed little decline in performance (**Figure 1**, Supplementary Table S2). Alternative supervised models showed similar training and testing properties and thus, alternative models will not be presented in further detail (Supplementary Figure S1). To understand the contributions of each gene towards the supervised analysis, we also generated a ranked order of genes using the SamR algorithm (Supplementary Table S1). As expected, *EGFR* gene expression ranked among the most differential genes associated with mutation status. Pathway analysis of the over- and under-expressed genes documented signatures that have been previously reported in association with *EGFR* mutation such as upregulation of kinase activity and downregulation of apoptotic processes (Supplementary Figure S3). Thus, we concluded that using gene expression and across a wide variety model types and conditions, it was possible to obtain high specificity and NPV for EGFR. However, sensitivity was modest, averaging 81% and PPV was generally suboptimal at average 55% (**Figure 1A-C**, Supplementary Table S2). In other words, it was possible to predict samples that lacked the *EGFR* mutation signature (mSig), although many cases with an identical gene expression pattern lacked the activating mutation. We interpreted this to mean that some samples lacking the *EGFR* mutation nonetheless demonstrated gene expression signaling overlapping with samples that contained the *EGFR* mutation. All cases predicted to have the signature of *EGFR* mutation were labeled as mSig(+), whether the mutation was observed or not.

We then considered the properties of mSig(+) samples lacking the *EGFR* mutation. Noting that prior studies have validated a statistically significant association between activating *EGFR* mutations and the Bronchioid subtype (16), we considered the association between false positive calls for *EGFR* mutation and the molecular subtypes of LUAD (**Figure 1D-F**). In each of the training and validation datasets and considering the supervised predictor genes for *EGFR* mutation status, several reproducible patterns were observed. First, the EGFR mSig developed for the supervised analysis of *EGFR*-mt versus *EGFR* WT also clearly distinguished the Bronchioid subtype from the Magnoid and Squamoid subtypes, with most mSig(+) cases co-labeled as Bronchioid. Likewise, most of the *EGFR*-mt cases and nearly all of the false positive mSig(+) patients (*EGFR* WT/mSig(+)) were associated with the Bronchioid gene expression subtype. Conversely, most of the *EGFR*-mt cases falsely predicted as mSig(-) (false negative) were non-Bronchioid samples. In other words, the supervised mSig(+) prediction appeared to largely represent a subset of the unsupervised Bronchioid molecular subtype.

We next considered whether *EGFR* WT/mSig(+) cases share other properties with *EGFR*-mt cases, including clinical outcomes. First, we confirmed that *EGFR*-mt patients have more favorable outcomes than *EGFR* WT patients in three-year overall survival (log-rank p-value = 0.13, **Figure 2A**)(39). We then stratified the outcomes of those patients with EGFR mSig(+) as a function of *EGFR* mutation status showing a similar favorable outcome for *EGFR*-mt patients similar to *EGFR* WT/mSig(+) patients (log-rank p-value = 0.05, Cox likelihood ratio test p-value = 0.02) (**Figure 2B**). In many cohorts including UNC, MSKCC and TCGA, it has previously reported that the Bronchioid subtype also has a similar favorable outcome compared to Squamoid and Magnoid subtypes (15). In short, Bronchioid subtype, *EGFR*-mt, and *EGFR* WT/mSig(+) belong to a group with shared gene expression and differential patient outcome.

**Figure 2.**
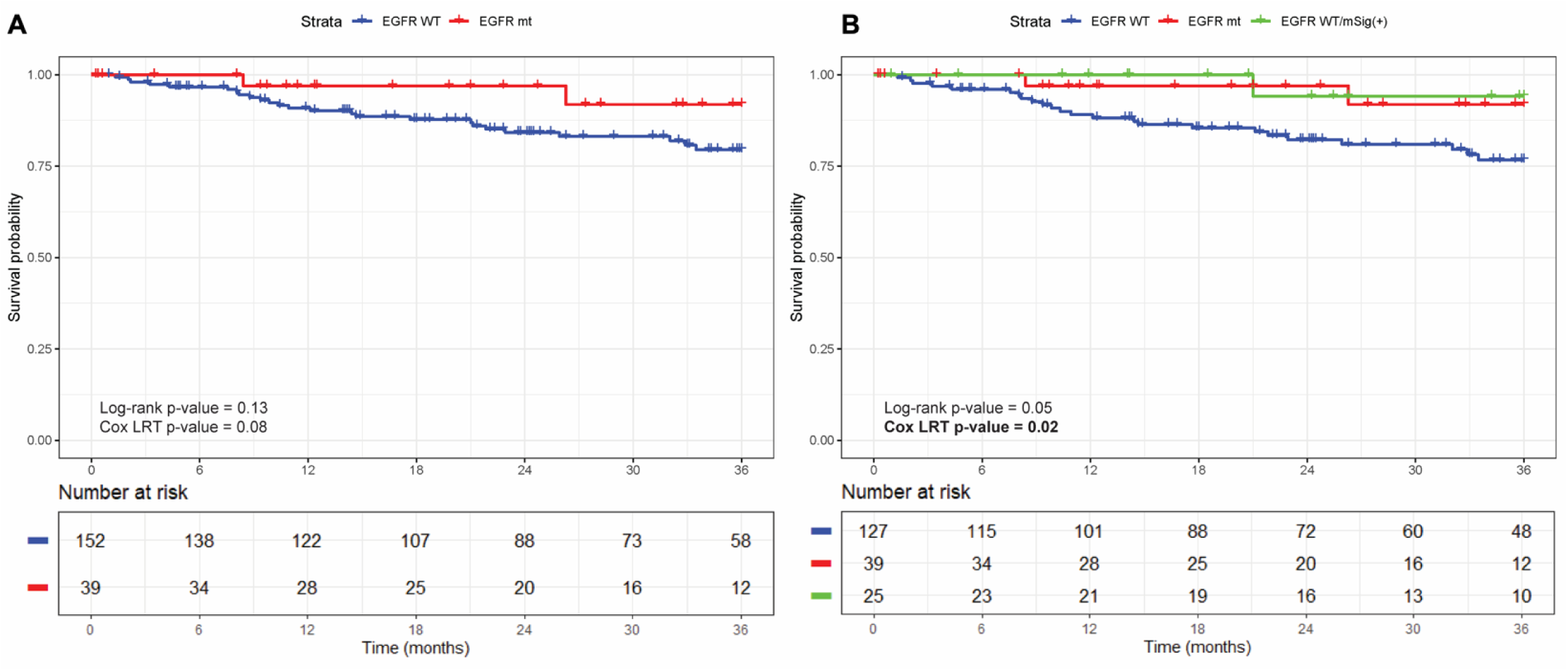
Three-year overall survival analysis based on EGFR-related signature groups in lung adenocarcinoma cohorts. P-values were calculated using the log-rank test and the Cox proportional hazards model (likelihood ratio test, LRT). Three-year Kaplan–Meier survival curves comparing in two groups (EGFR-mt, red, and EGFR WT, blue) (**A**) and three groups (EGFR mt, WT, EGFR WT/mSig(+), green) (**B**).

### EGFR Mutation Signature Correlates with LUAD Lineage Markers, Unsupervised Expression Subtypes, and Driver Genomic Alteration Events in LUAD

We next considered whether the shared EGFR pathway, expression signature and clinical outcomes might correlate with other molecular markers such as lineage-directing transcription factors or other driver gene mutations. To further define true and false positive mSig(+) patients, we examined the set of commonly altered driver LUAD genes and selected pulmonary lineage markers, the most highly associated with the canonical markers *NKX2-1* and *TP63* (40, 41). In our study as well, lung lineage genes were statistically associated with expression subtypes with Bronchioid (**Figure 3**). The Squamoid group was highly associated with *TP63* with overall lower *NKX2-1*, while the Magnoid group was relatively low for both markers. Such a pattern demonstrates that unsupervised molecular subtypes, supervised mutation expression signatures, and *EGFR* mutation itself correlate with expression of critical lung lineage markers.

**Figure 3.**
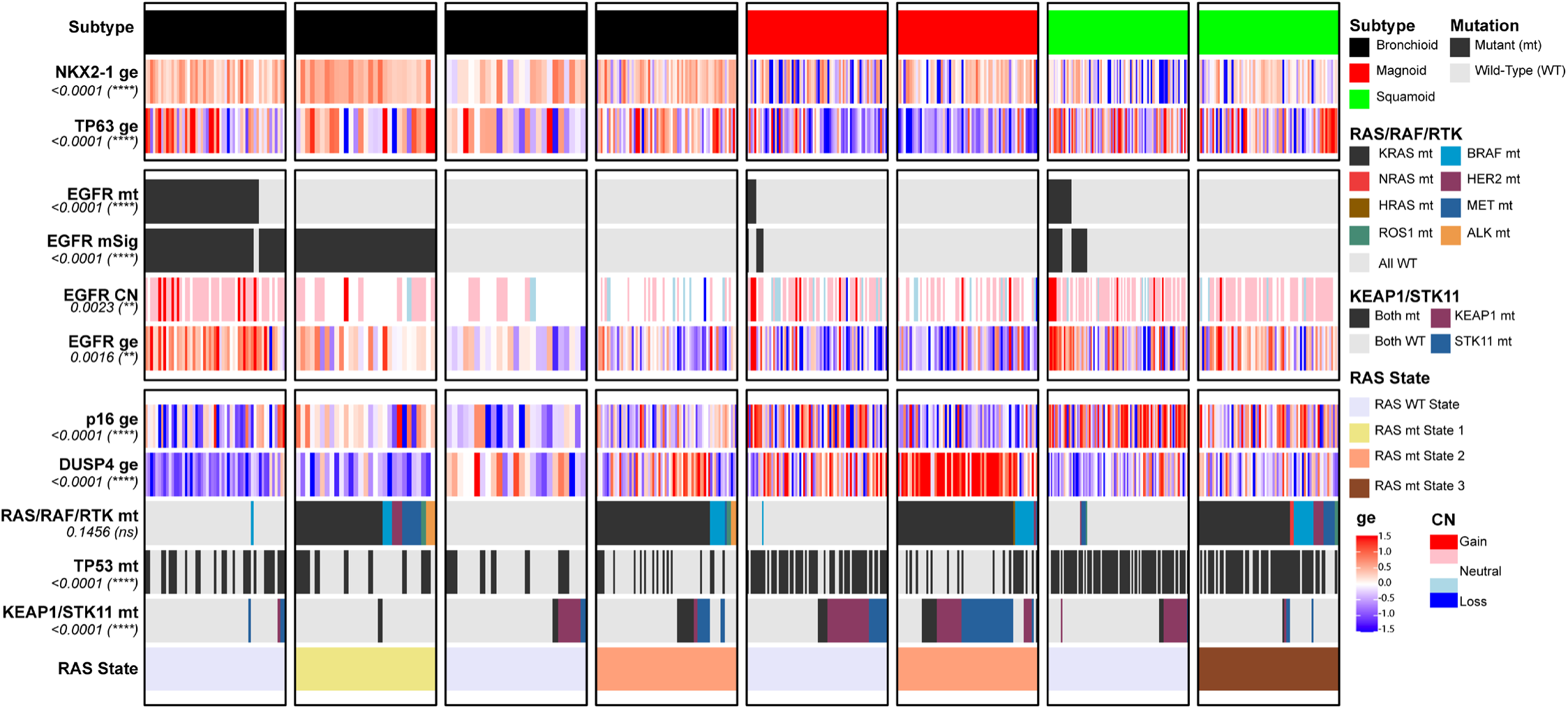
Integrative analysis of genomic alterations and gene expression across molecular subtypes of LUAD. Samples (n = 486, TCGA LUAD) are represented in columns and grouped by molecular subtype. Gene expression differences were assessed using one-way ANOVA, and categorical genomic alterations were evaluated using the chi-square test. Statistical significance reflects subtype-specific differences in the genomic features shown on the right. (* p < 0.05; ** p < 0.01; *** p < 0.001; **** p < 0.0001; ns, not significant). ge, gene expression; WT, wild-type; mt, mutant; CN, copy number.

We further investigated the correlation of driver gene alterations and mSig(+) cases as a function of unsupervised expression subtypes. Seventy five percent of mSig(+) patients (n=80/106) were of Bronchioid subtype (Bronchioid vs. EGFR mSig: OR = 9.2, *p*-value < 0.001), with the remaining 25% mostly Squamoid and very few Magnoid subtype (Supplementary Figure S4A, Supplementary Table S3). Inspection of *EGFR* gene expression follows a similar pattern with most of the highly expressed cases in the Bronchioid subgroup found in association with *EGFR*-mt, *EGFR* amplification, or both in association with mSig(+) status (Supplementary Figure S4A, B, Supplementary Tale S4). A significantly lower or absent *EGFR* expression and absent amplification was observed across the Magnoid group in association with *EGFR* WT and mSig(-) status (Supplementary Figure S4C). The few *EGFR*-mt Magnoid cases that were observed did not generally present with an associated mSig(+) status, a pattern also observed to a lesser degree in Squamoid samples (Supplementary Figure S4D). Interestingly, approximately half of the mSig(+) Bronchioid patients expressed *EGFR* at much more modest level, all of which were *EGFR* WT and nearly all of which had a documented alternative canonical RAS-activating mutation. By contrast, although most Squamoid samples expressed *EGFR* at higher levels, the mSig(+) signature was generally absent and although RAS mutations were common, almost none demonstrated the mSig(+) pattern.

The Bronchioid subtype was remarkable in that 75% of cases had either a canonical *EGFR* mutation or RAS mutation, with 50% of the tumors with RAS mutations demonstrating the mSig(+) signature and 50% showing an alternative signaling pathway. The observation that RAS mutations assume more than one signaling configuration has been previously reported by Skoulidis and others (10, 42), although to our knowledge the emphasis was on differential signaling in association with *TP53* mutation, *CDKN2A* loss, *STK11* loss, and low *NKX2-1* gene expression and not the RAS-mutant (RAS-mt) group described here which is most remarkable for demonstrating the *EGFR* mSig(+) signal in combination with high expression of *NKX2-1*, *TP53* WT and *STK11* WT. In support of the previously reported heterogenous KRAS signaling subtype characterization (10), we investigated the role of *CDKN2A* alteration including inactivating mutations / losses (data not shown), and gene expression (**Figure 3**). We confirmed patterns of differential *CDKN2A* gene expression, including relatively low expression in the *EGFR*-mt samples, a finding which could be interpreted as the loss of senescence pathways in tumors with an oncogene addiction to activated *EGFR*. Likewise, RAS- mt tumors of the Magnoid subgroup, but not other RAS-mt samples, demonstrated statistically significant decreased *CDKN2A* expression (RAS/RAF/RTK-mt vs. p16/CDKN2A ge: OR = 0.33, p-value <0.001) (Supplemental Figure S4C).

### DUSP4 as a Potential Therapeutic Target of EGFR Wild-Type LUAD

Since most *EGFR* mutations are in the Bronchioid subtype and most *EGFR* WTs are non-Bronchioid, comparison of *EGFR*-mt versus WT could be confounded by genes that define tumor molecular subtype. To the extent that the differences in molecular subtypes are driven by differences in driver genes, confounding may not be a problem. However, as Figure 3 demonstrates, other factors such as lineage transcription factors are a strong component of the tumor subtype expression signatures. For example, since Bronchioid and Magnoid differ dramatically by *NKX2-1* expression, *EGFR* expression, copy number, and mutation, genes associated with lineage and EGFR signaling will be highly confounded. Lineage confounding (and other potential unknown confounders) of *EGFR* mutation status might be partially overcome by stratifying differential gene expression between driver gene variants within a single molecular subtype. Although a number of supervised analysis strata could be entertained, we considered *EGFR* mutation status for mt versus WT within the Bronchioid subtype and across all samples and all subtypes (Supplementary Figure S5A). Although the prediction direction and effects were highly correlated for many genes within the Bronchioid subtype versus across all patients, candidates related to RAS signaling not previously appreciated rose in their discriminating power. Differentially expressed genes in Bronchioid tumors recapitulated the distinct molecular profiles observed across all subtypes in multiple cohorts (Supplementary Figure S5B-C). Most notably, the gene dual specificity phosphatase 4 (*DUSP4)* was identified significantly upregulated in the *EGFR*-mt/mSig(+) versus EGFR mSig(-) within the Bronchioid subtype via GSEA (normalized enrichment score = 1.3, nominal p-value <0.01, data not shown), but not across all subtypes. DUSP4 is a protein phosphatase that inactivates kinases by dephosphorylating both the phosphoserine/threonine and phosphotyrosine residues. It negatively regulates members of the mitogen-activated protein (MAP) kinase superfamily (MAPK/ERK, SAPK/JNK, p38), which are associated with cellular proliferation and differentiation (43, 44). As such, it is reasonable that differential *DUSP4* gene expression might correlate with differential activation of the MAPK pathway. Projecting *DUSP4* expression onto the integrated driver data revealed intriguing patterns both for the *EGFR* WT Magnoids as well as other subgroups of LUAD patients. For all *EGFR*-mt and mSig(+)/RAS-mt tumors, *DUSP4* expression was uniformly low. In contrast, *DUSP4* demonstrated significantly elevated expression, especially in EGFR WT/*KRAS*-mt Bronchioid and Magnoid tumors, but not *KRAS*-mt Squamoid tumors.

### Coordination of Lineage, Oncogene Signaling, Proliferation, Metabolic / Oxidative Stress in LUAD

We next sought to augment the previously reported RAS classification proposed by Skoulidis et al. with unsupervised gene expression subtypes first reported more than 20 years ago (10, 15). In this scheme, three unsupervised tumor classes (Bronchioid, Squamoid, Magnoid) correlate highly with the lineage markers *NKX2-1* and *TP63*, suggesting an underlying cellular context of differentiation or cell-of-origin among tumors characterized as LUAD (**Figure 4**). Within each cellular context, tumors demonstrated differential patterns of alterations in common driver genes associated with oncogene activation, altered proliferation, and metabolic / oxidative stress. Oncogene activation was defined by either an mSig(+) signature associated with *EGFR*-mt or RAS-mt (referred to as RAS Mutation 1 or Type I), or an mSig(-) / DUSP4-high RAS mutation 2 (Type II). The remaining RAS-mt samples, entirely limited to Squamoid subtype demonstrated neither mSig(+) signature nor high DUSP4 expression, suggesting a RAS mutation 3 (Type III) as described by Skoulidis et al. for its high proportion of *TP53* mutations. Greater than 90% of all Squamoid tumors, including RAS Type III patients, had TP53 mutations. In contrast, EGFR-mt/mSig(+), mSig(+)/RAS Type I and mSig(-)/RAS Type II were infrequently TP53-mt. Interestingly, Squamoid samples, although largely lacking the mSig and having approximately 50% RAS mutation, demonstrated frequent amplifications and higher *EGFR* gene expression. We empirically describe this group as demonstrating atypical *EGFR* expression based on lack of mSig and the coordinated expression of *EGFR* in the presence of RAS mutation.

**Figure 4.**
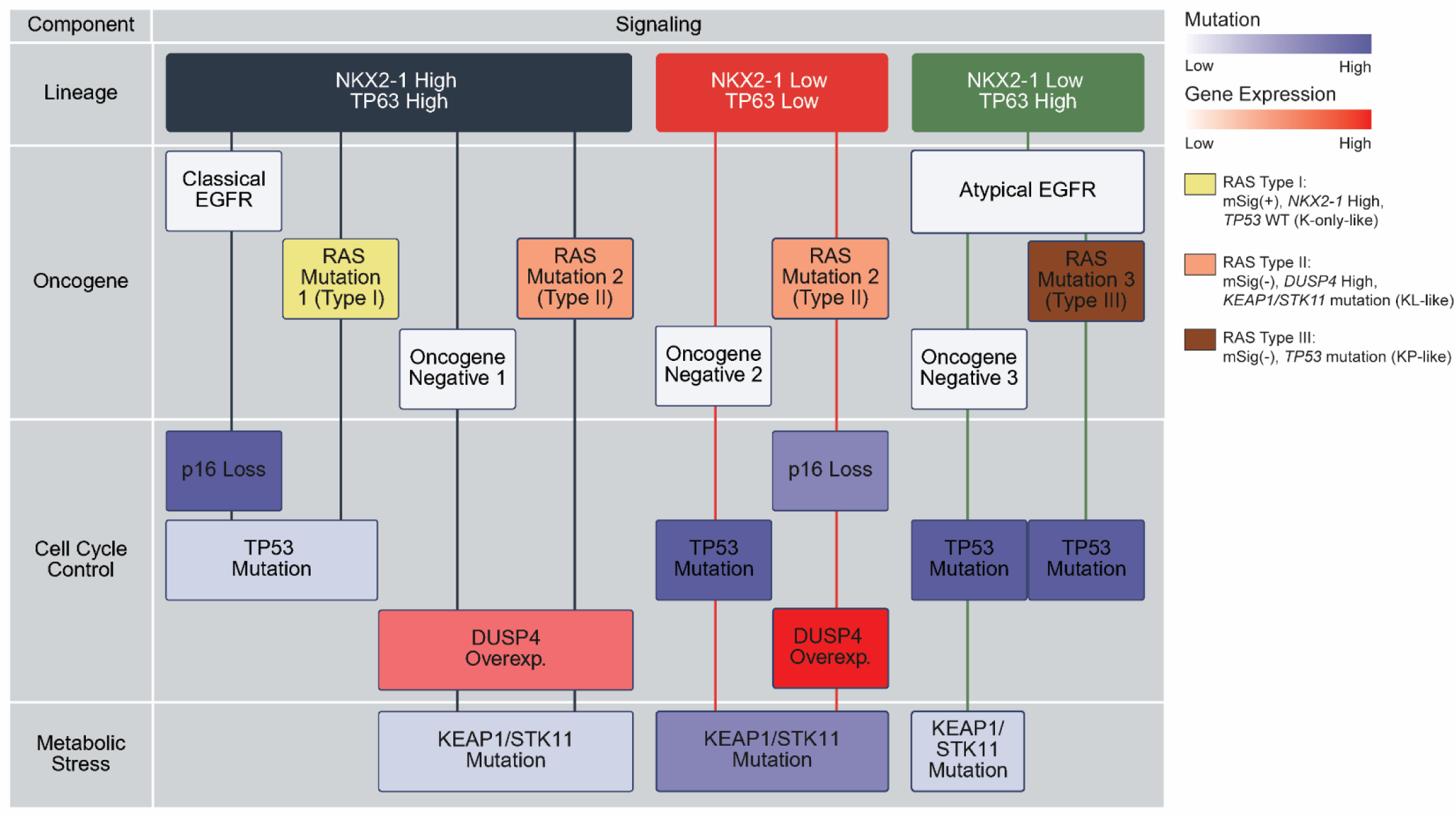
A proposed framework linking four biological components across lineage-defined molecular subtypes of LUAD. This schematic model illustrates the proposed subtypes of LUAD stratified by lineage features (*NKX2-1* and *TP63* expression), and integrates alterations across key oncogenic, cell cycle control, and metabolic stress-related signaling. Each component highlights distinct molecular features that collectively define the biological heterogeneity and potential therapeutic vulnerabilities of each subtype. RAS mutations (Type I, II, and III) correspond to previously characterized KRAS subgroups described (42): Type I tumors are mSig-positive, *NKX2-1* high, and *TP53* wild-type (K-only–like); Type II tumors are mSig-negative, characterized by *DUSP4* overexpression and frequent KEAP1/STK11 co-mutations (KL-like); and Type III tumors are mSig-negative with *TP53* mutation (KP-like).

Skoulidis et al. (10, 42) described not only the association of one RAS mutation type with *TP53* mutation such as we see in Squamoid expression class, but also of an alternative means of impacting proliferation through altered *CDKN2A*. We also observe this to a limited degree with RAS Type II cases but record more prominent losses of *CDKN2A* for EGFR-mt/mSig(+) patients. Considering DUSP4, a repressor of RAS signaling, as the 3^rd^ most prominent proliferation signal demonstrates a high degree of coordination across the subtypes proposed in the model. Finally, as suggested by both Skoulidis and others (17, 21), altered oxidative / metabolic stress mutations co-occur in reproducible ways, especially in the Magnoid group where nearly every RAS-mt case has a concomitant oxidative / metabolic stress mutations as opposed to almost none of the Type I or Type III RAS-mt cases.

## Discussion

EGFR kinase mutations generate a reproducible gene expression signature, with consistent prediction accuracy across predictor type, model parameters, and expression assay. The prominence of a single *EGFR* mutation signature is unexpected since other LUAD oncogenes including *BRAF* and *KRAS* often display multiple, context-dependent signatures. For BRAF, it signals differently depending on tissue and anatomic location (e.g., colonic epithelium vs. melanocyte) (45). KRAS signaling is heterogeneous within a single tissue, shaped by co-mutations (*TP53*, *KEAP1*, *STK11*, etc.) and pulmonary lineage marker *NKX2-1*(12–14). Context is also crucial for *EGFR*. Although EGFR signaling is relevant in many tumor types, kinase mutations occur almost exclusively in LUAD. Herein, nearly all EGFR-mt cases were mSig(+), a signature enriched in *NKX2- 1*/*TP63* co-expressing Bronchioid cells. Importantly, mSig(+) is not unique to *EGFR* mutations; we report that canonical RAS mutations can also yield this profile (**Figure 4**). Thus, while mSig(+) is critical for certain Bronchioid tumors, it is not synonymous with the Bronchioid subtype. Roughly half of Bronchioid tumors are mSig(–), many driven by canonical RAS mutations signaling with MAPK repressor DUSP4. Another quarter lack both mSig(+) and canonical RAS mutations but appear to follow DUSP4-driven signaling.

Prior work links Bronchioid tumors to a more peripheral “terminal respiratory unit” (TRU) phenotype (15). TRU refers to the distal lung—respiratory bronchioles, alveolar ducts, and alveoli—where gas exchange occurs. The Bronchioid subtype was historically associated with “bronchioloalveolar carcinoma (BAC),” often reflecting lepidic or differentiated LUAD histology, sharing associations such as *EGFR* mutation, nonsmoker status, higher *NKX2-1* expression, and favorable outcomes (16, 46). This report strengthens the view that these diverse phenotypes, genotypes, and morphologies may reflect a unified molecular classification with clinical relevance. At least 75% of Bronchioid tumors were EGFR- or RAS-driven through mSig(+) *EGFR* mutation or Type 1/2 RAS mutations. The remaining 25% lacked clear oncogenes but signaling cascades were identified that are similar to Type 2 RAS cases. By contrast, Squamoid tumors were uniformly *TP53*-mt, showing moderate to high *EGFR* expression, which is atypical, as these tumors are overwhelmingly *EGFR* mSig(–). The Squamoid transcription factor profile reflects reduced NKX2-1 dependence, consistent with a subset of LUAD but rarely contextualized alongside mutation patterns, as in this report. About half of Squamoid tumors display evidence of combined *TP53* mutations with canonical RAS mutations, a known pattern that we show is linked to this expression subtype. The other half are oncogene negative but display signaling distinct from Bronchioids, as mSig(–) Squamoids are *DUSP4*-low, unlike the *DUSP4*-high mSig (-) Bronchioids. The Magnoid subtype, historically associated with large cell undifferentiated carcinoma histology, exhibited very low expression of lung lineage markers (*TP63*, *NKX2*-1) and absent *EGFR* expression. The Magnoids are a solid/large-cell histologic subtype of LUAD, fall into two genotypes. Roughly half of the Magnoid samples carry RAS mutations with *STK11* and/or *KEAP1* mutations, combined with high *DUSP4* and *TP53* WT status. The remainder of tumors from the Magnoid subtype were *TP53*-mt and either oncogene-negative or co-mutant for *KEAP1*/*STK11*, with moderate *DUSP4* expression. It is plausible to expect that the oncogene-negative Magnoids would demonstrate signaling unique to that subtype (and different from Bronchioid and Squamoid) for the purposes of modeling and therapy.

The discovery of EGFR mutations as therapeutic targets in LUAD paralleled landmark findings such as BCR-ABL fusions in chronic myelogenous leukemia (CML) and HER2 amplifications in breast cancer, fueling modern cancer genomics(47–49)￼. The focus on druggable somatic variants as the basis for LUAD classification has been especially productive in LUAD patient care, but this focus may overshadow the role of lineage and differentiation state for understanding tumor genesis and model systems. Leukemia illustrates the alternative approach of emphasizing the role of lineage and differentiation state as opposed to simply focusing on druggable targets, as has been the case in LUAD. For example, the BCR-ABL fusion defines CML as a disease, as well as defines a variety of therapeutic pathways.(50–53). Similarly, the transcription factor fusion, PML-RARA, defines acute promyelocytic leukemia, where differentiation arrest occurs at the promyelocyte stage (54). Sarcomas also demonstrate lineage-defining lesions, such as SS18-SSX in synovial sarcoma and EWSR1-ATF1 in clear cell sarcoma (55–57). In epithelial tumors, oncogenic variants often align with the cell-of-origin and differentiation. Salivary gland cancers exemplify this: secretory carcinoma harbors *ETV6*-*NTRK3* fusions in >95% of cases (58); the ductal variant shows *AR* overexpression in >70% and *HER2* (59); mucoepidermoid carcinoma has *CRTC1*-*MAML2* (60). While salivary carcinomas showcase striking correlations between morphologic subtype and molecular lesion, many solid tumors with less distinct morphology can also be subclassified through molecular profiling, which reveals underlying cell-of-origin. Copy number alterations also show tissue specificity, embedding canonical targets within tumor-specific genomic patterns. Thus, oncogenesis is not random but rather highly tissue-context dependent: drivers active in one tissue may be inert in another, even when pathways overlap.

Just as transcriptional and genomic features distinguish tumor subtypes, oncogenic signaling behavior is context dependent (61). *BRAF* mutations, for example, are targetable in melanoma but resistant in colorectal cancer (62). Similarly, EGFR kinase mutations are restricted to LUAD, despite EGFR amplification in other tumors. If universally oncogenic, EGFR kinase mutations would appear across tissues; instead, their selection depends on lineage, reinforced by co-expression of NKX2-1 (46). *EGFR* mutations also align with expression subtypes, mirroring lineage-dependent subgroups in breast cancer (63). Histopathology echoes this, with differentiated morphologies in Bronchioids and solid/undifferentiated morphologies in Magnoids. Together, these findings emphasize that cell identity—defined by origin, differentiation, and lineage factors—shapes malignant phenotypes. Even without clear histology, molecular lesions can reveal lineage, as in triple-negative breast cancer.

The pace of new oncogene discovery has slowed, and many known drivers remain difficult to target, such as transcription factors (TP63, MYC), hormone receptors, and “hard-to-drug” genes like RAS and *PIK3CA* (15, 64–69). Some candidates (e.g., FGFR inhibitors) face challenges due to rarity, low efficacy, or toxicity.

Thus, LUAD drug development is often illustrated as a pie chart in which small slices represent sparse subsets of “oncogene positive” cases, while a large portion remains “other genes” or “oncogene negative.” This pie chart model is appealing but it is oversimplified by lumping oncogene-negative tumors and ignoring co-occurrence patterns (e.g., RAS with *KEAP1*/*STK11*). The pie chart approach fails to highlight expression subtype contexts, making therapeutic translational research even more difficult (Supplementary Figure S6) (70). Therefore, we proposed a layered cake concept integrating oncogenes, mutations, and signaling cascades across multiple layers that together lead to oncogenesis and tumor subtype phenotypes.

This study proposes a lineage-directed framework integrating driver genes and expression subtypes. By examining EGFR signaling, LUAD gene expression subtypes, and canonical drivers, we develop an integrated model (**Figure 4**) that offers an improved structure compared to the pie chart. Oncogene-negative cases are stratified by subtype context, and difficult targets like *KRAS* are logically grouped. Although the multiple strata suggested by this model would challenge clinical trialists, this framework reflects the heterogeneity of the disease and appears feasible and comparable in complexity to other heterogeneous diseases such as leukemia with its many lineage-driven subsets.

In summary, *EGFR* mutations are tightly linked to the mSig(+) pattern, especially in Bronchioid LUAD, though RAS mutations can also generate this profile. Magnoid tumors show little or no mSig(+) signal, even with *EGFR* mutations, while Squamoid tumors express *EGFR* abundantly yet remain mSig(–). These findings underscore that LUAD taxonomy is best understood through lineage and co-mutation context. Figure 4 illustrates how clinically relevant groups can be defined without overwhelming sparsity, potentially reframing “oncogene-negative” tumors within a broader taxonomy. Even absent druggable targets, such stratification may enable more personalized treatment approaches.

## Acknowledgements

We thank the UTHSC Center for Cancer Research for input and support. This work is supported by the National Cancer Institute (NCI) of the National Institutes of Health (NIH) under award number U01CA272541 (LM), which is part of the Metabolic Dysregulation and Cancer Risk Program (MeDOC) Consortium; NCI R01CA262112 (LM), NCI R01CA262296 (YC, DNH), NCI U24CA264021 (DNH), NCI UG1CA233333 (DNH), NCI R01CA211939 (DNH). The views expressed in this manuscript are solely of the authors and do not reflect the official policy of the Departments of Army/Navy/Air Force, Department of Defense, USUHS, or the United States government. This research content is solely the responsibility of the authors and does not necessarily represent the official views of the NIH.

## Contributions

Funding Acquisition: DNH, LM, YC; Supervision: DNH, LM; Conceptualization: WL, MK, DNH; Data Collection: HJ, HYC, WL, MW, MH; Data Generation and Curation: MK, HJ, WL; Writing: MK, LM, DNH; Reviewing: All authors

## Conflict of Interest Statement

DNH and MDW has a patent on lung cancer and molecular markers. The other authors declare no potential conflicts of interest.

## Supplementary Data

**Supplementary Figure S1.**
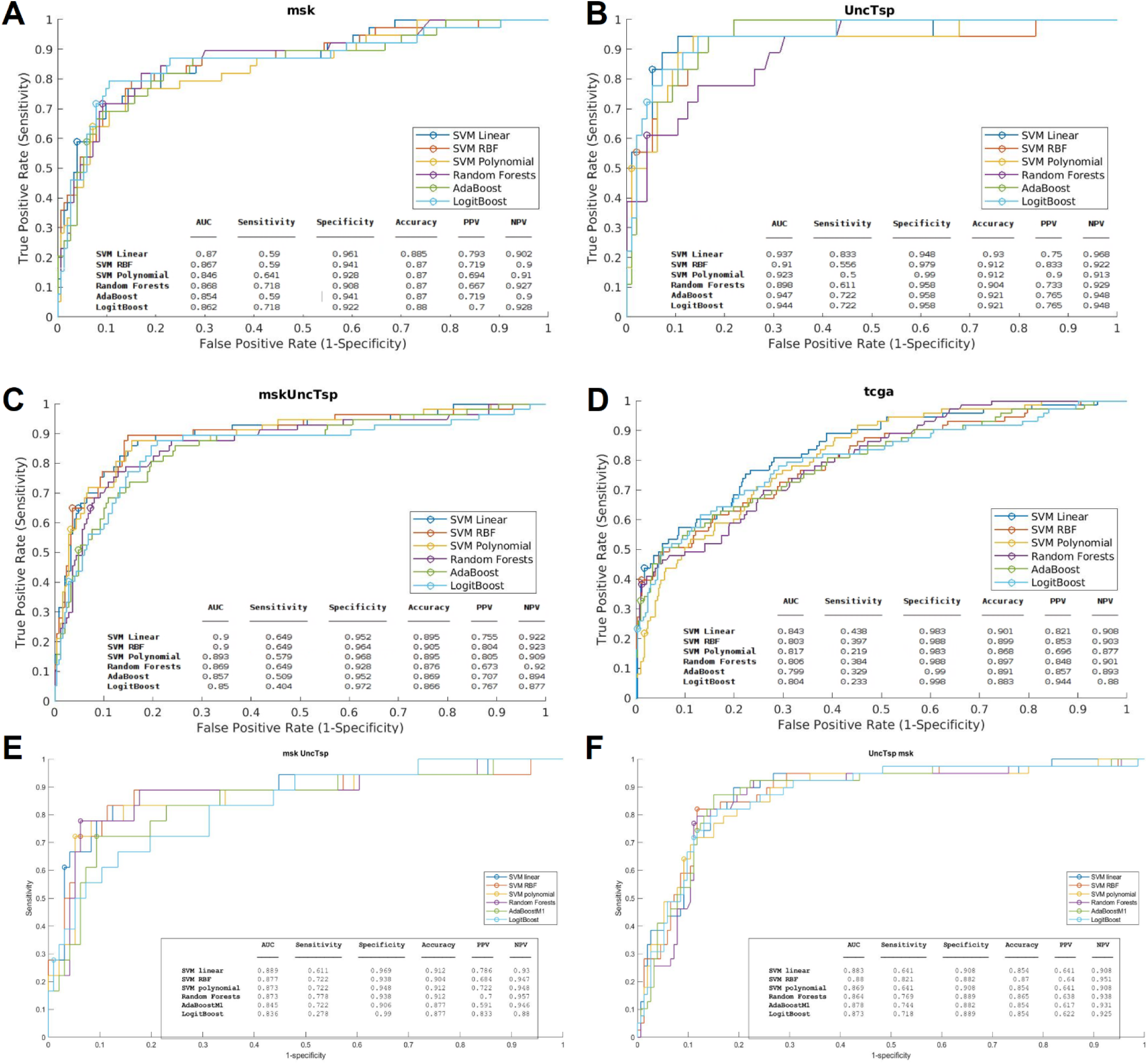
ML-based performance evaluation and validation of the EGFR mSig. The top 1,000 genes selected by *t*-test were used as input features for 10-fold cross-validation. ROC curves of six different machine learning models are shown in distinct colors. **A**. MSKCC, **B**. UNC + TSP, **C**. MSKCC + UNC + TSP, **D**. TCGA, **E**. Training: MSKCC, Validation: UNC+TSP, **F**. Training: UNC+TSP, Validation: MSKCC.

**Supplementary Figure S2.**
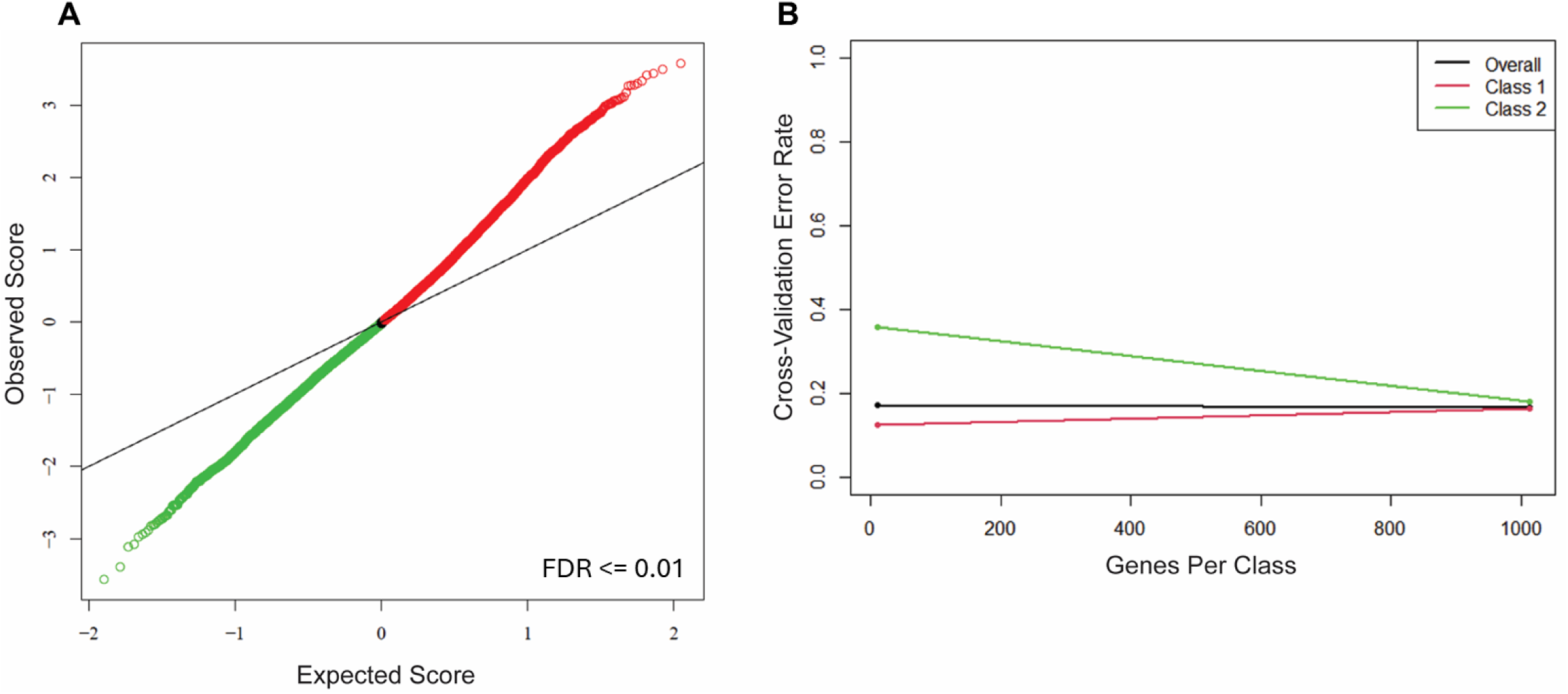
Selection and validation of EGFR mutation signature genes. **A**. DEGs between EGFR-mt and EGFR WT LUAD samples were identified using SamR, and only statistically significant DEGs with a false discovery rate (FDR) < 1% were selected as candidate signature genes. **B**. Cross-validation of EGFR-related signature genes was performed using ClaNC. A total of 1,020 genes were selected as the final EGFR mSig based on the lowest classification error rate (Error rate < 0.2). Red and green lines indicate different patient groups used in the classification model.

**Supplementary Figure S3.**
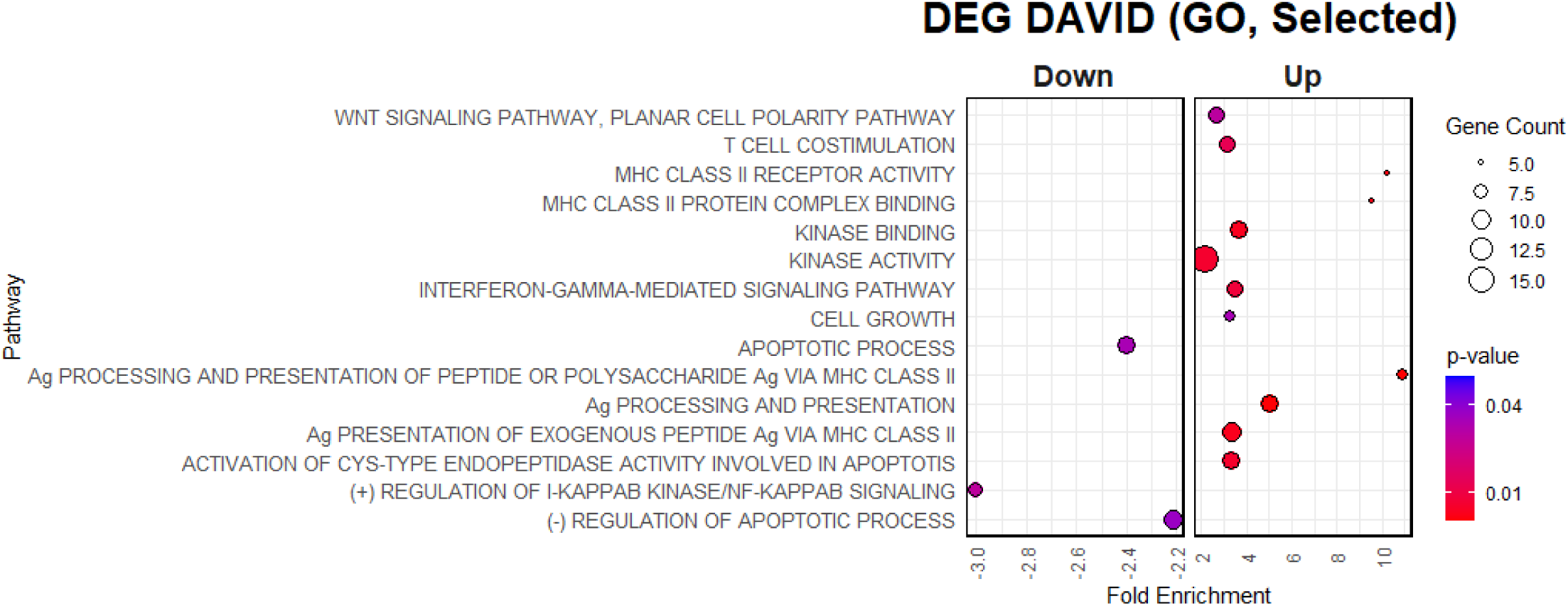
Pathway enrichment analysis of EGFR mSig genes. Biological pathways associated with 1,020 EGFR-related signature genes (690 upregulated, 330 downregulated) were analyzed using the web-based DAVID functional annotation tool. Gene Ontology (GO) term enrichment results are visualized in a dot plot. The x-axis indicates fold enrichment for each pathway, dot size represents the number of genes involved, and dot color reflects statistical significance based on p-values.

**Supplementary Figure S4.**
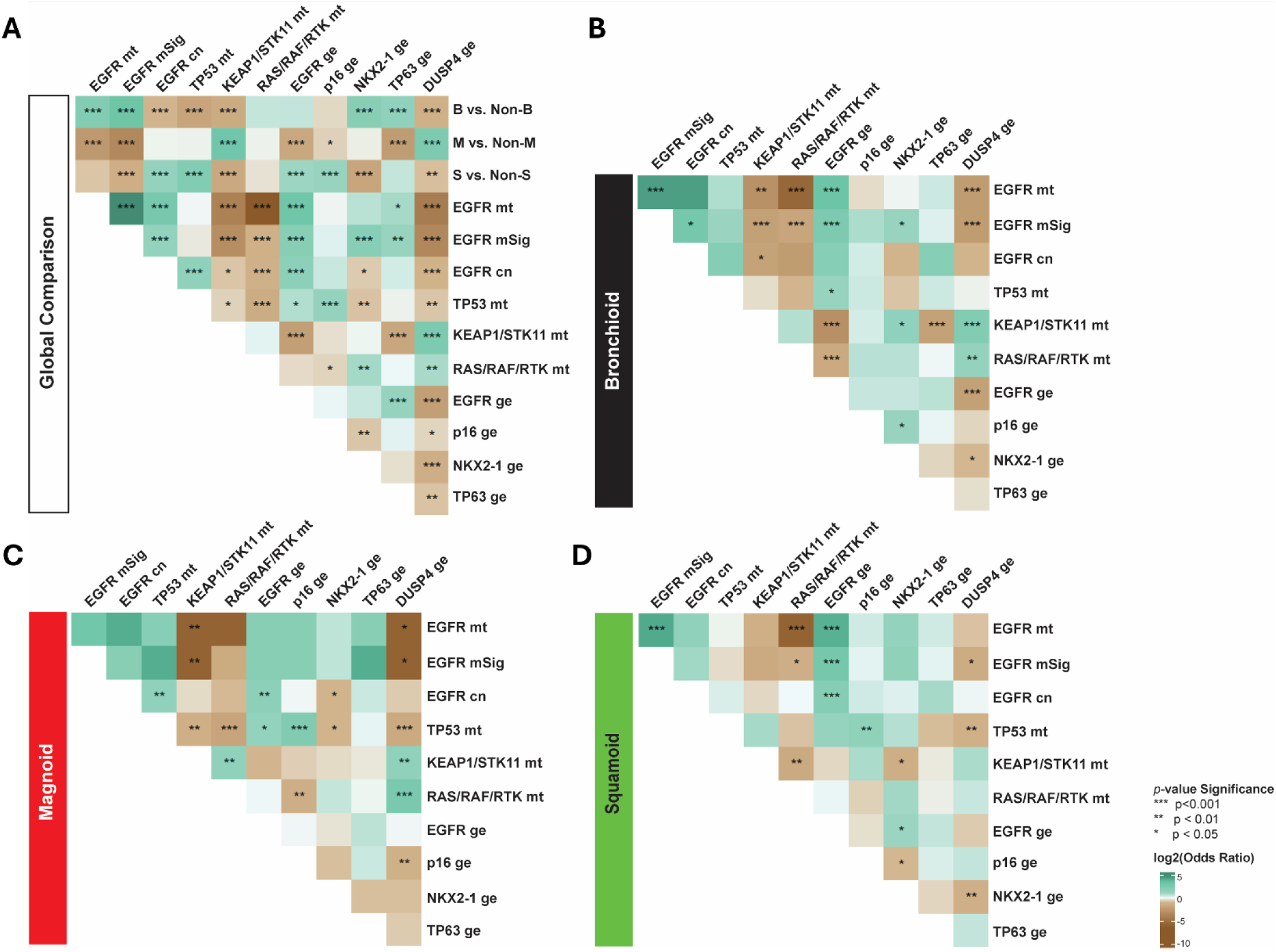
Co-occurrence and mutual exclusivity of selected genomic alterations. Heatmaps display odds ratios for the co-occurrence of genomic events between pairs of genes, with color indicating the magnitude of the odds ratio. Co-occurring events exhibit odds ratios greater than 1, while mutually exclusive events have odds ratios less than 1. The strength of mutual exclusivity (or co-occurrence) increases as the odds ratio deviates further from 1. P-values were calculated using Fisher’s exact test. ge, gene expression; mt, mutant; CN, copy number. **A.** Global comparison across all LUAD samples. **B–D.** Subtype-specific analyses of mutual exclusivity within the Bronchioid (**B**), Magnoid (**C**), and Squamoid (**D**) subtypes.

**Supplementary Figure S5.**
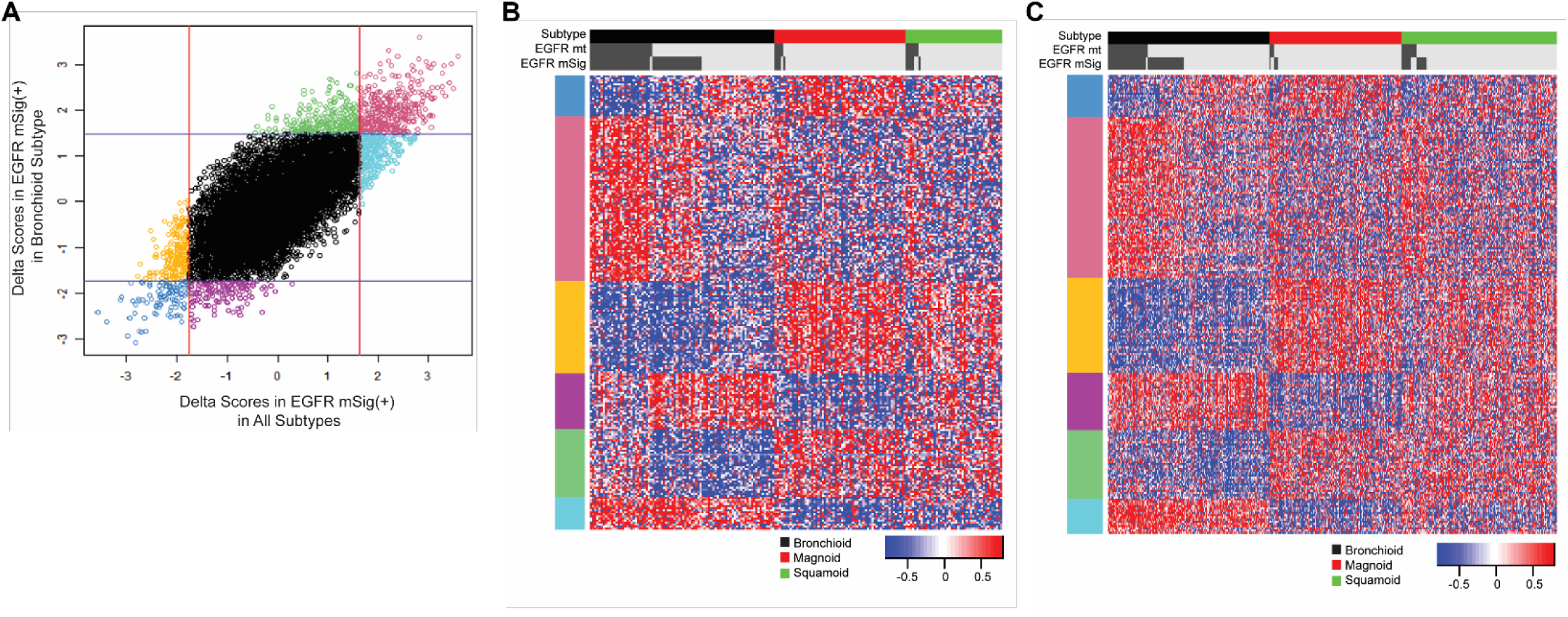
Differentially expressed genes in the Bronchioid subtype and distinctive molecular profiles defined by the EGFR mSig. **A**. Dot plot showing the distribution of delta scores calculated by SamR for 11,807 genes. Each dot represents a gene, with the x-axis showing the delta score for genes in the EGFR-predicted group across all LUAD subtypes, and the y-axis showing the delta score for genes in the EGFR-predicted group specifically within the Bronchioid subtype. Genes are colored based on empirically chosen cutoff values indicated by red vertical lines (delta score > 1.6 or < -1.7) and blue horizontal lines (delta score > 1.5 or < -1.7), capturing statistically significant differential expression. The 221 genes highlighted in color were selected for further analysis shown in **B** and **C**. **B**, **C**. Heatmaps display gene expression profiles of the selected 221 genes across different LUAD subtypes, with top annotations for EGFR mutation status and EGFR prediction in the MSKCC (**B**) and TCGA (**C**) cohorts. Each row represents one gene, with the left color bar indicating subgroup classification from **A**. Dot colors correspond to gene expression patterns as indicated by the heatmap color bars.

**Supplementary Figure S6.**
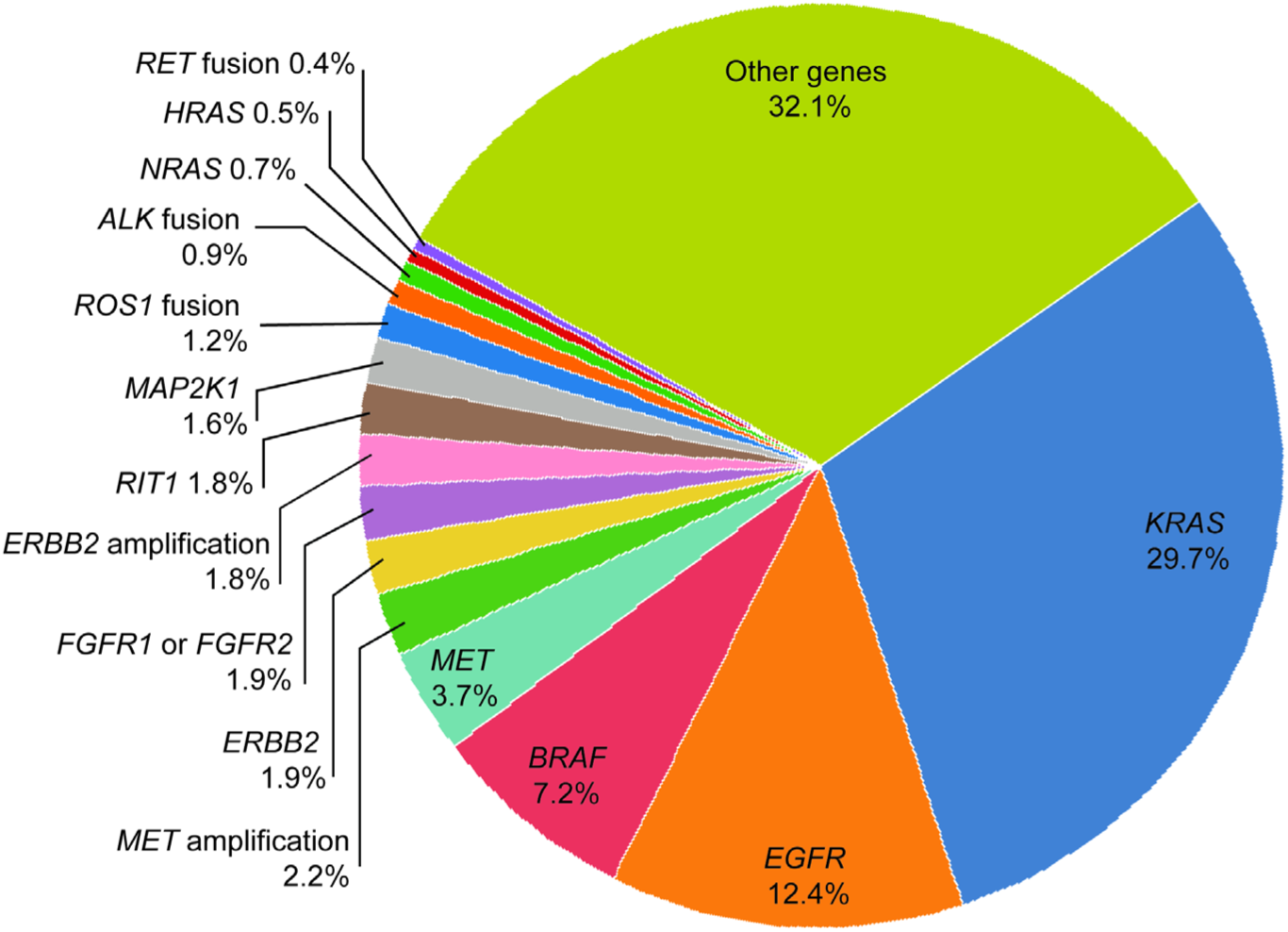
Conventional pie chart of oncogenic driver gene mutation frequencies in LUAD. This figure is presented to exemplify the traditional use of pie charts for mutation frequency representation (TCGA PanCancer Atlas LUAD from cBioPortal).

**Supplementary Table 1.** EGFR mSig gene list (1,020 genes) ST1 is supplied separately.

**Supplementary Table 2.**
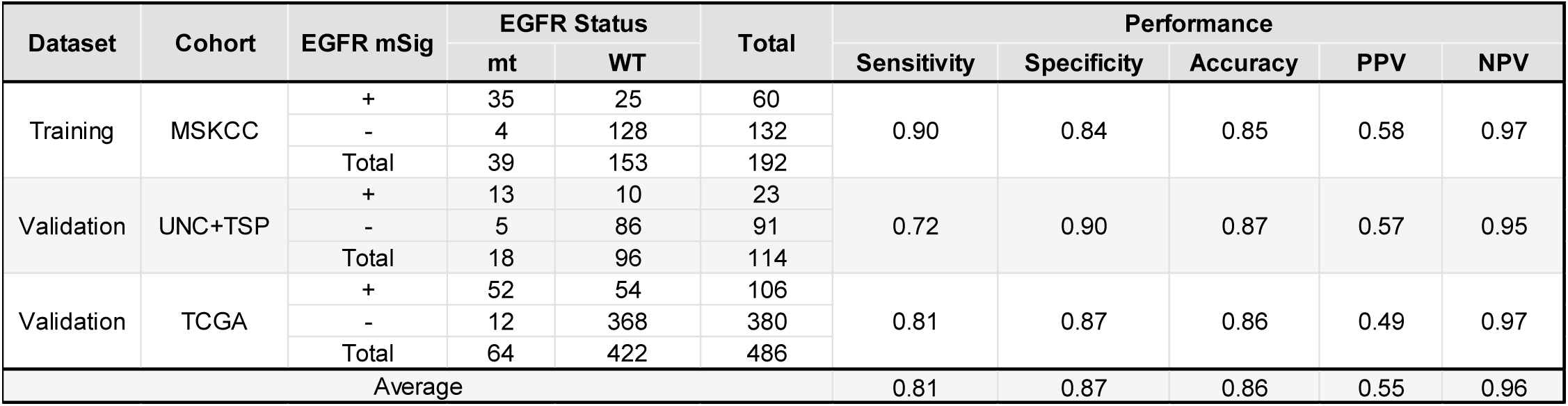
Performance test results between EGFR-like mutant group and WT group based on EGFR-related gene signature. mt, mutant, WT, wild-type, PPV, positive predictive value, NPV, negative predictive value.

**Supplementary Table S3.**
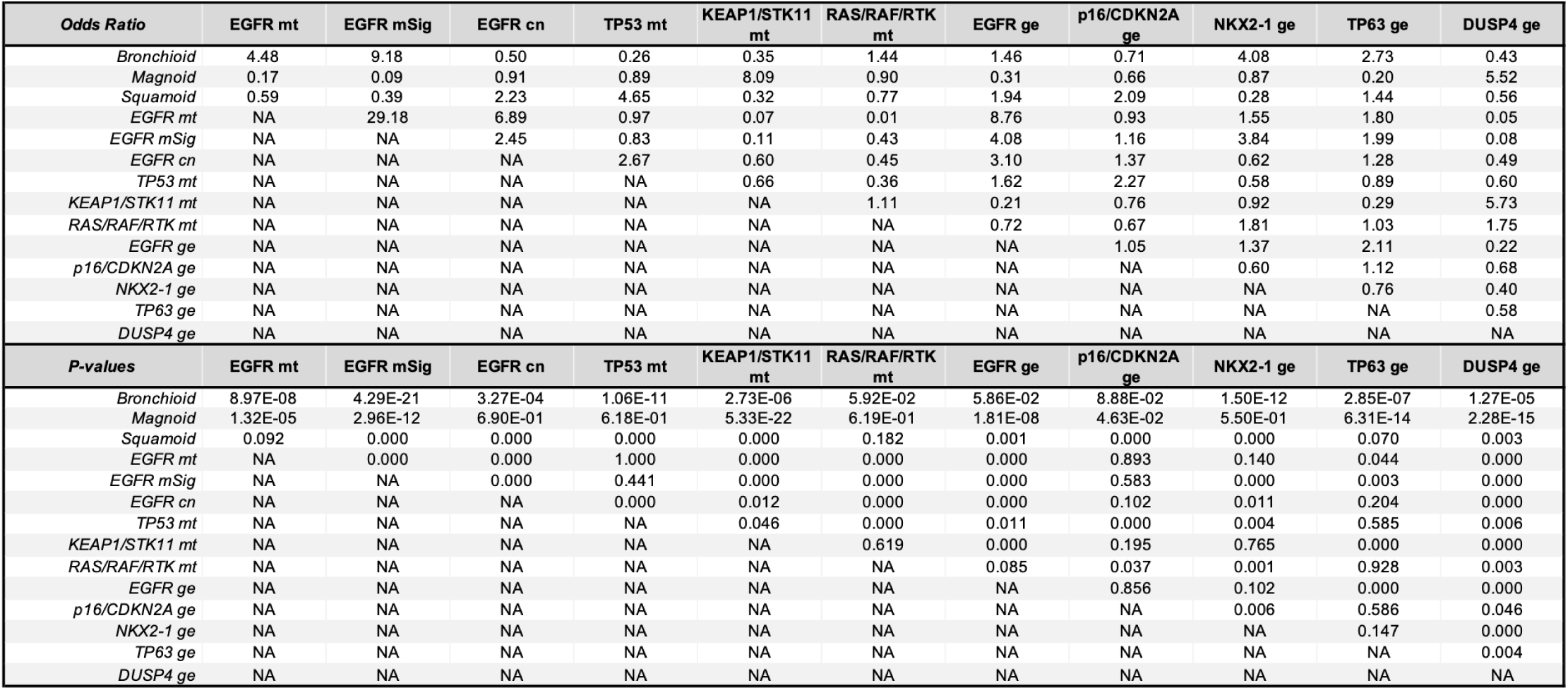
Odds Ratio and p-value; Subtype vs. Non-Subtypes.

| Supplementary Table S3. Odds Ratio and p-value; Subtype vs. Non-Subtypes. |  |  |  |  |  |  |  |  |  |  |  |
| --- | --- | --- | --- | --- | --- | --- | --- | --- | --- | --- | --- |
| Odds Ratio | EGFR mt | EGFR mSig | EGFR cn | TP53 mt | KEAP1/STK11 mt | RAS/RAF/RTK mt | EGFR ge | p16/CDKN2A ge | NKX2-1 ge | TP63 ge | DUSP4 ge |
| Bronchioid | 4.48 | 9.18 | 0.50 | 0.26 | 0.35 | 1.44 | 1.46 | 0.71 | 4.08 | 2.73 | 0.43 |
| Magnoid | 0.17 | 0.09 | 0.91 | 0.89 | 8.09 | 0.90 | 0.31 | 0.66 | 0.87 | 0.20 | 5.52 |
| Squamoid | 0.59 | 0.39 | 2.23 | 4.65 | 0.32 | 0.77 | 1.94 | 2.09 | 0.28 | 1.44 | 0.56 |
| EGFR mt | NA | 29.18 | 6.89 | 0.97 | 0.07 | 0.01 | 8.76 | 0.93 | 1.55 | 1.80 | 0.05 |
| EGFR mSig | NA | NA | 2.45 | 0.83 | 0.11 | 0.43 | 4.08 | 1.16 | 3.84 | 1.99 | 0.08 |
| EGFR cn | NA | NA | NA | 2.67 | 0.60 | 0.45 | 3.10 | 1.37 | 0.62 | 1.28 | 0.49 |
| TP53 mt | NA | NA | NA | NA | 0.66 | 0.36 | 1.62 | 2.27 | 0.58 | 0.89 | 0.60 |
| KEAP1/STK11 mt | NA | NA | NA | NA | NA | 1.11 | 0.21 | 0.76 | 0.92 | 0.29 | 5.73 |
| RAS/RAF/RTK mt | NA | NA | NA | NA | NA | NA | 0.72 | 0.67 | 1.81 | 1.03 | 1.75 |
| EGFR ge | NA | NA | NA | NA | NA | NA | NA | 1.05 | 1.37 | 2.11 | 0.22 |
| p16/CDKN2A ge | NA | NA | NA | NA | NA | NA | NA | NA | 0.60 | 1.12 | 0.68 |
| NKX2-1 ge | NA | NA | NA | NA | NA | NA | NA | NA | NA | 0.76 | 0.40 |
| TP63 ge | NA | NA | NA | NA | NA | NA | NA | NA | NA | NA | 0.58 |
| DUSP4 ge | NA | NA | NA | NA | NA | NA | NA | NA | NA | NA | NA |
| P-values | EGFR mt | EGFR mSig | EGFR cn | TP53 mt | KEAP1/STK11 mt | RAS/RAF/RTK mt | EGFR ge | p16/CDKN2A ge | NKX2-1 ge | TP63 ge | DUSP4 ge |
| Bronchioid | 8.97E-08 | 4.29E-21 | 3.27E-04 | 1.06E-11 | 2.73E-06 | 5.92E-02 | 5.86E-02 | 8.88E-02 | 1.50E-12 | 2.85E-07 | 1.27E-05 |
| Magnoid | 1.32E-05 | 2.96E-12 | 6.90E-01 | 6.18E-01 | 5.33E-22 | 6.19E-01 | 1.81E-08 | 4.63E-02 | 5.50E-01 | 6.31E-14 | 2.28E-15 |
| Squamoid | 0.092 | 0.000 | 0.000 | 0.000 | 0.000 | 0.182 | 0.001 | 0.000 | 0.000 | 0.070 | 0.003 |
| EGFR mt | NA | 0.000 | 0.000 | 1.000 | 0.000 | 0.000 | 0.000 | 0.893 | 0.140 | 0.044 | 0.000 |
| EGFR mSig | NA | NA | 0.000 | 0.441 | 0.000 | 0.000 | 0.000 | 0.583 | 0.000 | 0.003 | 0.000 |
| EGFR cn | NA | NA | NA | 0.000 | 0.012 | 0.000 | 0.000 | 0.102 | 0.011 | 0.204 | 0.000 |
| TP53 mt | NA | NA | NA | NA | 0.046 | 0.000 | 0.011 | 0.000 | 0.004 | 0.585 | 0.006 |
| KEAP1/STK11 mt | NA | NA | NA | NA | NA | 0.619 | 0.000 | 0.195 | 0.765 | 0.000 | 0.000 |
| RAS/RAF/RTK mt | NA | NA | NA | NA | NA | NA | 0.085 | 0.037 | 0.001 | 0.928 | 0.003 |
| EGFR ge | NA | NA | NA | NA | NA | NA | NA | 0.856 | 0.102 | 0.000 | 0.000 |
| p16/CDKN2A ge | NA | NA | NA | NA | NA | NA | NA | NA | 0.006 | 0.586 | 0.046 |
| NKX2-1 ge | NA | NA | NA | NA | NA | NA | NA | NA | NA | 0.147 | 0.000 |
| TP63 ge | NA | NA | NA | NA | NA | NA | NA | NA | NA | NA | 0.004 |
| DUSP4 ge | NA | NA | NA | NA | NA | NA | NA | NA | NA | NA | NA |

**Supplementary Table S4.**
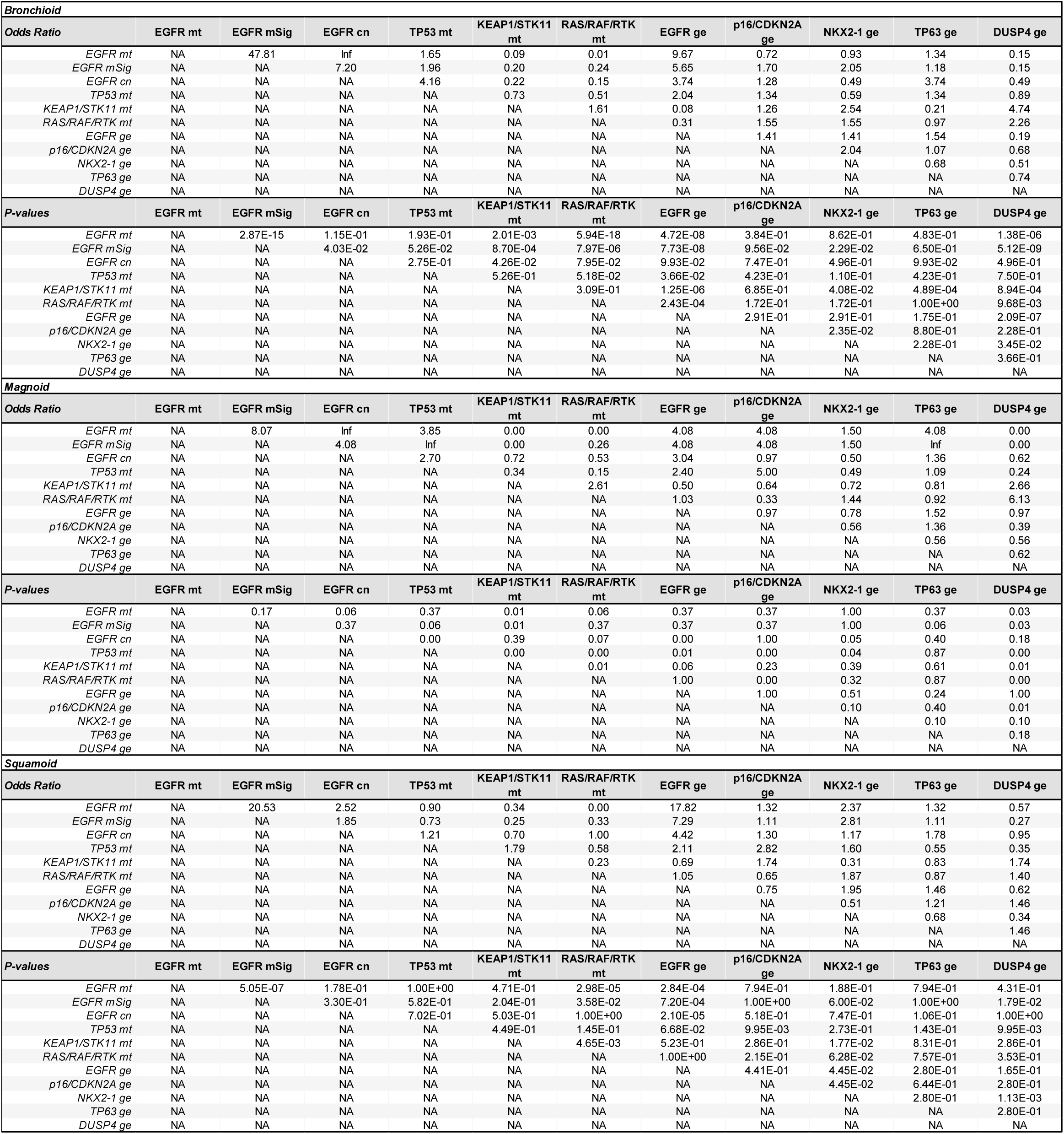
Odds Ratio and p-value; Within Subtype. mt, mutant, cn, copy number, ge, gene expression. Inf, infinite value due to zero count.

