## Supplementary Figure 1 for "Lung Adenocarcinoma Just Desserts: An Expanding Pie of Activating Oncogenes or a Layer Cake of Integrated Alterations"

**Supplementary Figure S1**. **ML-based performance evaluation and validation of the EGFR mSig.** The top 1,000 genes selected by t-test were used as input features for 10-fold cross-validation. ROC curves of six different machine learning models are shown in distinct colors. **A**. MSKCC, **B**. UNC + TSP, **C**. MSKCC + UNC + TSP, **D**. TCGA, **E**. Training: MSKCC, Validation: UNC+TSP, **F**. Training: UNC+TSP, Validation: MSKCC

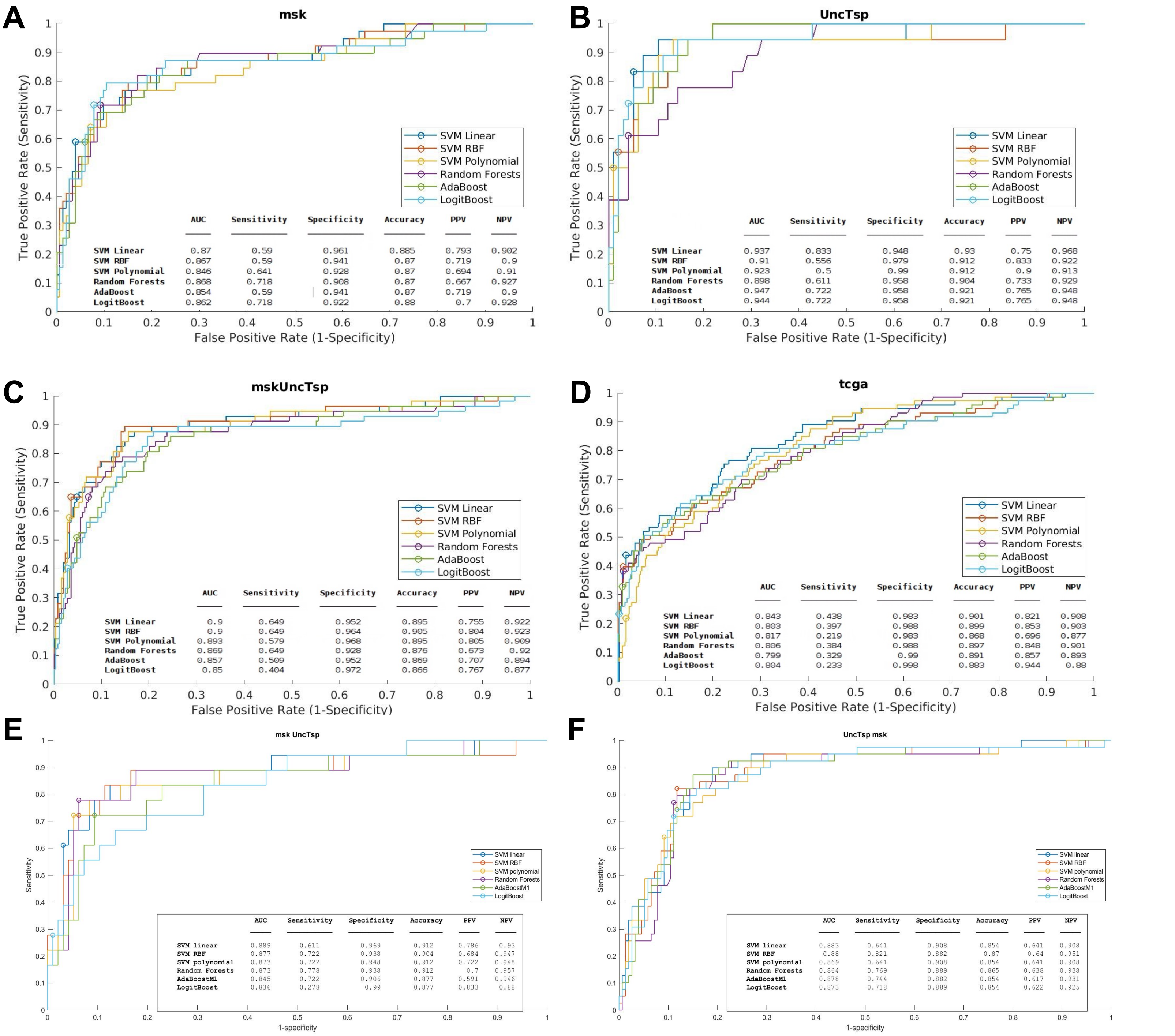
