## Supplementary Figure 2 for "Lung Adenocarcinoma Just Desserts: An Expanding Pie of Activating Oncogenes or a Layer Cake of Integrated Alterations"

**Supplementary Figure S2. Selection and validation of EGFR mutation signature genes.**

**A**. DEGs between EGFR-mt and EGFR WT LUAD samples were identified using SamR, and only statistically significant DEGs with a false discovery rate (FDR) < 1% were selected as candidate signature genes. **B**. Cross-validation of EGFR-related signature genes was performed using ClaNC. A total of 1,020 genes were selected as the final EGFR mSig based on the lowest classification error rate (Error rate < 0.2). Red and green lines indicate different patient groups used in the classification model.

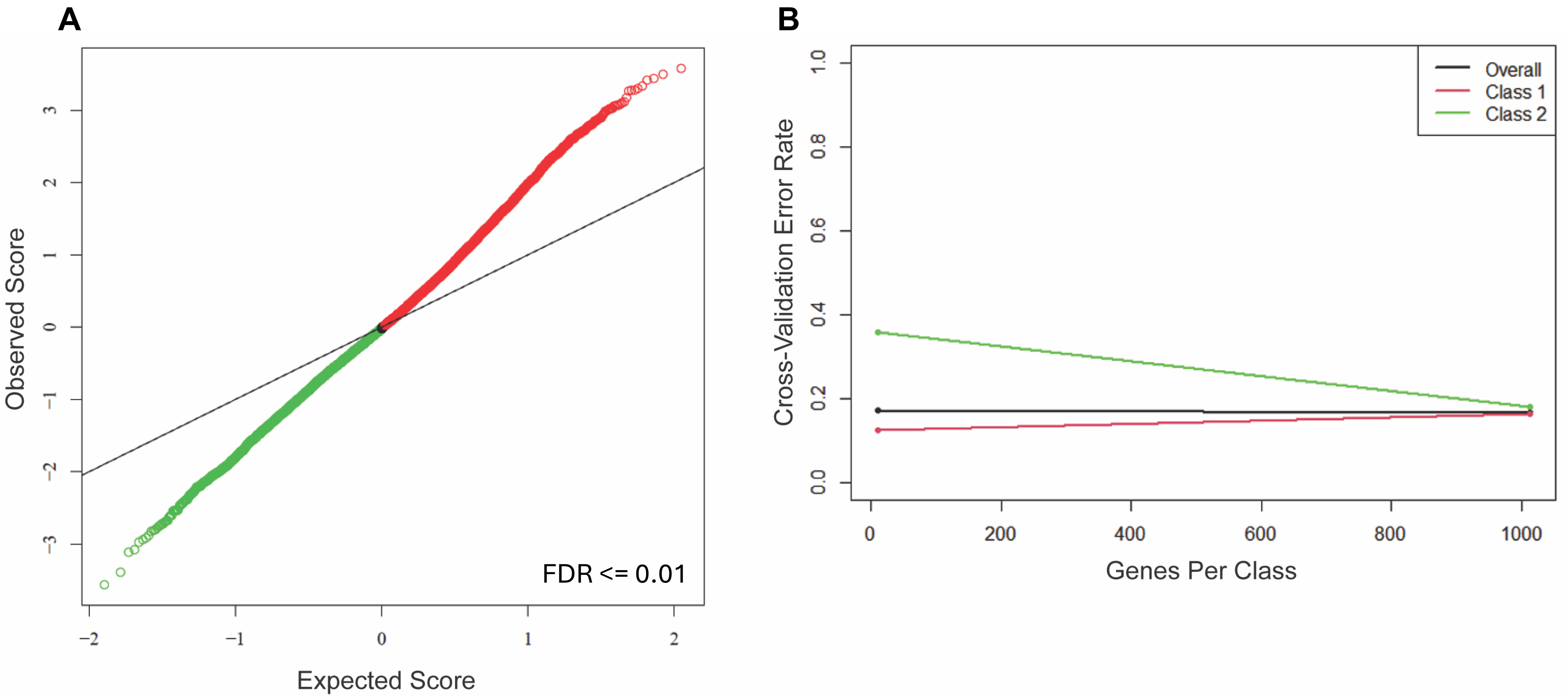
