## Supplementary Figure 3 for "Lung Adenocarcinoma Just Desserts: An Expanding Pie of Activating Oncogenes or a Layer Cake of Integrated Alterations"

**Supplementary Figure S3. Pathway enrichment analysis of EGFR mSig genes.**
Biological pathways associated with 1,020 EGFR-related signature genes (690 upregulated, 330 downregulated) were analyzed using the web-based DAVID functional annotation tool. Gene Ontology (GO) term enrichment results are visualized in a dot plot. The x-axis indicates fold enrichment for each pathway, dot size represents the number of genes involved, and dot color reflects statistical significance based on p-values.

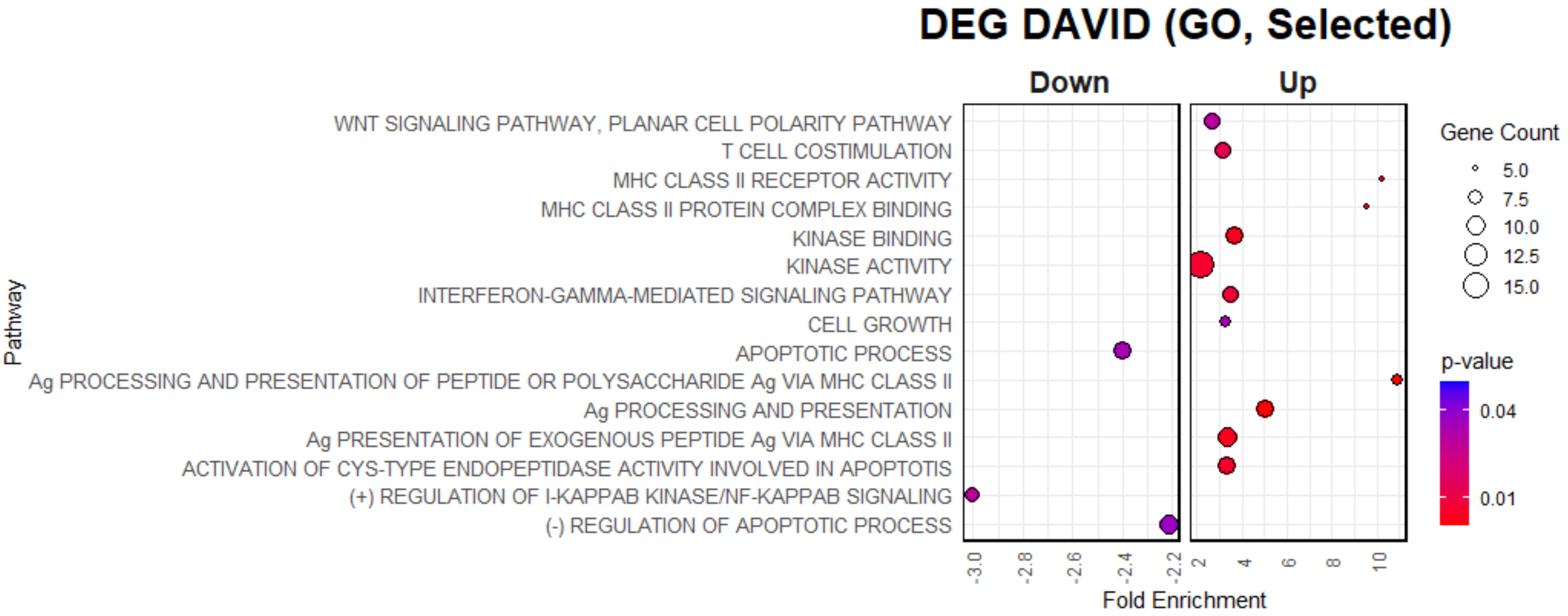
