## Supplementary Figure 4 for "Lung Adenocarcinoma Just Desserts: An Expanding Pie of Activating Oncogenes or a Layer Cake of Integrated Alterations"

**Supplementary Figure S4. Co-occurrence and mutual exclusivity of selected genomic alterations.** Heatmaps display odds ratios for the co-occurrence of genomic events between pairs of genes, with color indicating the magnitude of the odds ratio. Co-occurring events exhibit odds ratios greater than 1, while mutually exclusive events have odds ratios less than 1. The strength of mutual exclusivity (or co-occurrence) increases as the odds ratio deviates further from 1. P-values were calculated using Fisher's exact test. ge, gene expression; mt, mutant; CN, copy number. **A.** Global comparison across all LUAD samples. **B–D.** Subtype-specific analyses of mutual exclusivity within the Bronchioid (**B**), Magnoid (**C**), and Squamoid (**D**) subtypes.

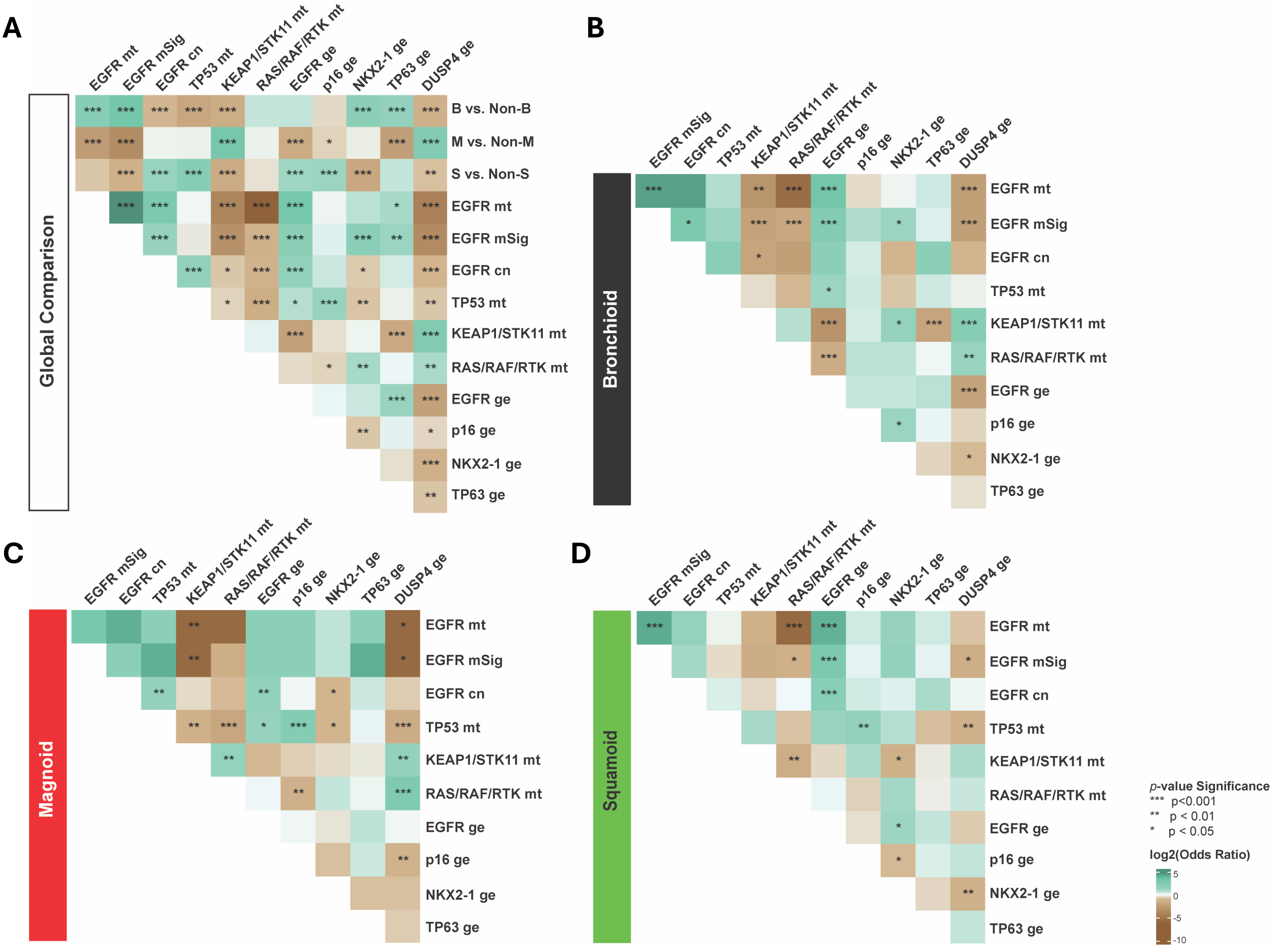
