## Supplementary Figure 5 for "Lung Adenocarcinoma Just Desserts: An Expanding Pie of Activating Oncogenes or a Layer Cake of Integrated Alterations"

**Supplementary Figure S5. Differentially expressed genes in the Bronchioid subtype and distinctive molecular profiles defined by the EGFR mSig. A**. Dot plot showing the distribution of delta scores calculated by SamR for 11,807 genes. Each dot represents a gene, with the x-axis showing the delta score for genes in the EGFR-predicted group across all LUAD subtypes, and the y-axis showing the delta score for genes in the EGFR-predicted group specifically within the Bronchioid subtype. Genes are colored based on empirically chosen cutoff values indicated by red vertical lines (delta score > 1.6 or < -1.7) and blue horizontal lines (delta score > 1.5 or < -1.7), capturing statistically significant differential expression. The 221 genes highlighted in color were selected for further analysis shown in **B** and **C**. **B**, **C**. Heatmaps display gene expression profiles of the selected 221 genes across different LUAD subtypes, with top annotations for EGFR mutation status and EGFR prediction in the MSKCC (**B**) and TCGA (**C**) cohorts. Each row represents one gene, with the left color bar indicating subgroup classification from **A**. Dot colors correspond to gene expression patterns as indicated by the heatmap color bars.

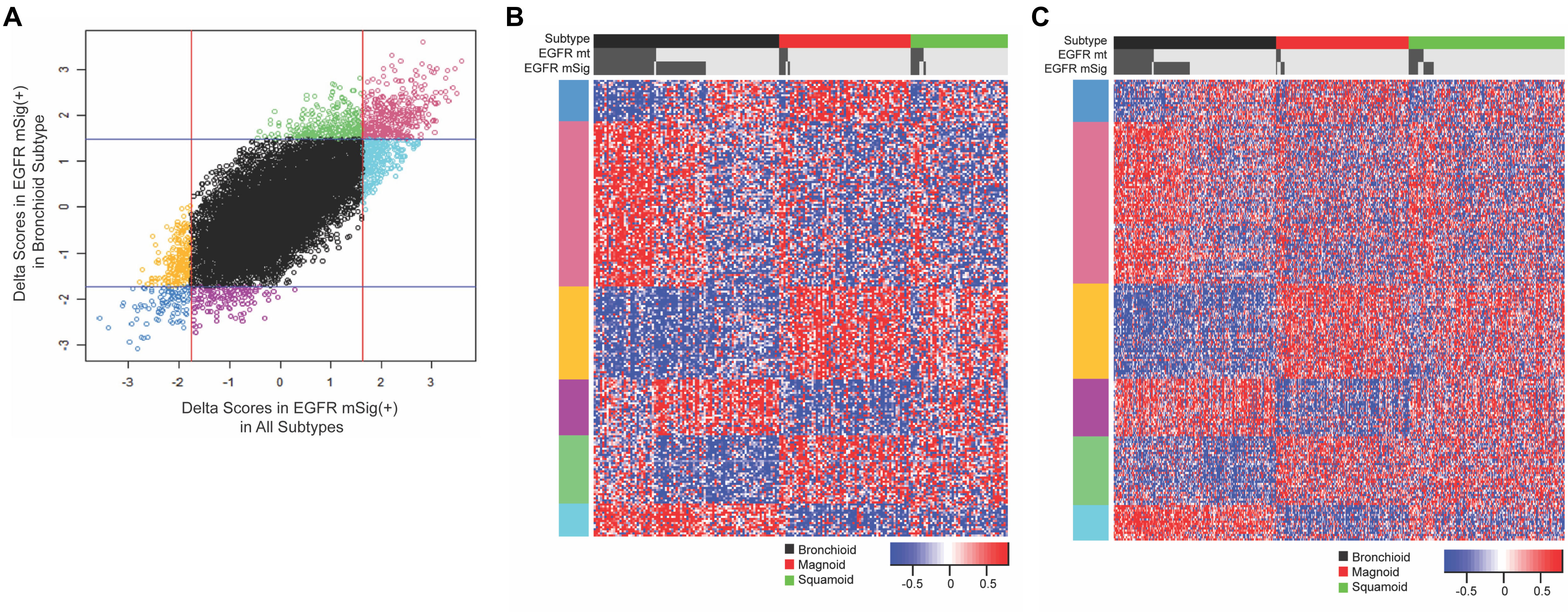
