## Supplementary Figure 6 for "Lung Adenocarcinoma Just Desserts: An Expanding Pie of Activating Oncogenes or a Layer Cake of Integrated Alterations"

**Supplementary Figure S6. Conventional pie chart of oncogenic driver gene mutation frequencies in LUAD.** This figure is presented to exemplify the traditional use of pie charts for mutation frequency representation (TCGA PanCancer Atlas LUAD, dataset from cBioPortal).

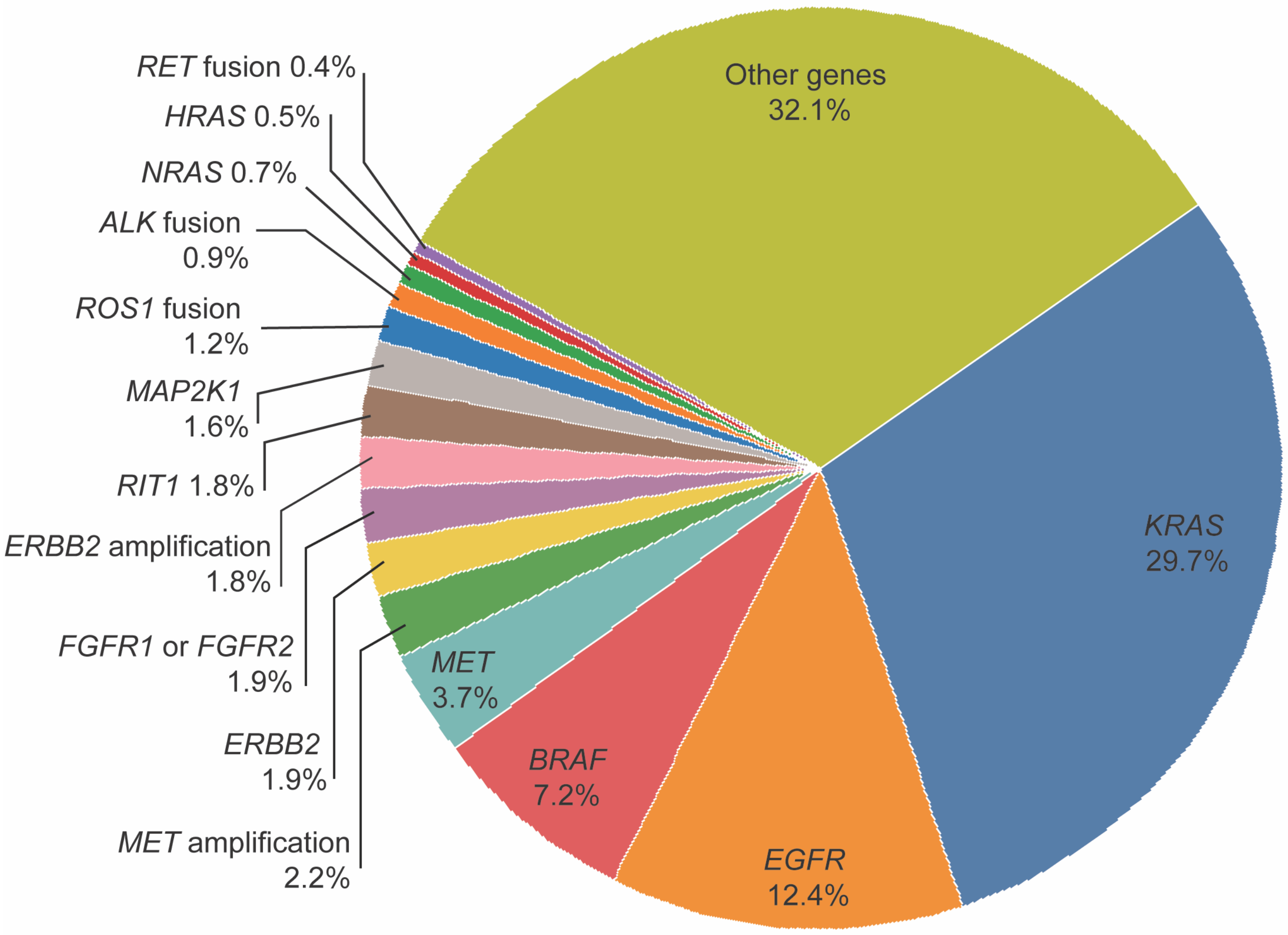
