## Supplementary Table 1 for "Lung Adenocarcinoma Just Desserts: An Expanding Pie of Activating Oncogenes or a Layer Cake of Integrated Alterations"

**Supplementary Table 1**. EGFR mutation signature gene list in ranked order.

| Rank | *Gene* | Delta Score | Fold Change | q-value |
|---|---|---|---|---|
| 1 | *HIP1* | 3.58 | 2.21 | 0.00 |
| 2 | PIGQ | 3.49 | 2.17 | 0.00 |
| 3 | *C16orf58* | 3.44 | 2.15 | 0.00 |
| 4 | UPK3A | 3.41 | 2.14 | 0.00 |
| 5 | *HMOX2* | 3.33 | 2.10 | 0.00 |
| 6 | RNF40 | 3.30 | 2.09 | 0.00 |
| 7 | *GALNT10* | 3.28 | 2.08 | 0.00 |
| 8 | KIAA0319L | 3.27 | 2.08 | 0.00 |
| 9 | *KIAA0494* | 3.27 | 2.07 | 0.00 |
| 10 | GPR177 | 3.17 | 2.03 | 0.00 |
| 11 | *C1orf149* | 3.11 | 2.01 | 0.00 |
| 12 | EGFR | 3.10 | 2.01 | 0.00 |
| 13 | *FLJ14154* | 3.08 | 2.00 | 0.00 |
| 14 | FTSJ2 | 3.07 | 1.99 | 0.00 |
| 15 | *MKL2* | 3.06 | 1.99 | 0.00 |
| 16 | LRRC31 | 3.06 | 1.99 | 0.00 |
| 17 | *KIAA0495* | 3.05 | 1.99 | 0.00 |
| 18 | RERE | 3.02 | 1.98 | 0.00 |
| 19 | *B4GALT7* | 3.02 | 1.98 | 0.00 |
| 20 | ADCY9 | 3.02 | 1.97 | 0.00 |
| 21 | *LRRC47* | 3.01 | 1.97 | 0.00 |
| 22 | C1orf174 | 2.99 | 1.96 | 0.00 |
| 23 | *HHLA3* | 2.99 | 1.96 | 0.00 |
| 24 | GPR172B | 2.98 | 1.96 | 0.00 |
| 25 | *PEX10* | 2.96 | 1.95 | 0.00 |
| 26 | CTF1 | 2.94 | 1.94 | 0.00 |
| 27 | *GUSB* | 2.93 | 1.94 | 0.00 |
| 28 | LDLRAP1 | 2.89 | 1.92 | 0.00 |
| 29 | *ROGDI* | 2.89 | 1.92 | 0.00 |
| 30 | FLJ10986 | 2.89 | 1.92 | 0.00 |
| 31 | *LYRM1* | 2.88 | 1.92 | 0.00 |
| 32 | ZNF688 | 2.87 | 1.91 | 0.00 |
| 33 | *HIRIP3* | 2.86 | 1.91 | 0.00 |
| 34 | PHLDB1 | 2.86 | 1.91 | 0.00 |
| 35 | *PPFIBP2* | 2.85 | 1.91 | 0.00 |
| 36 | PPIE | 2.85 | 1.90 | 0.00 |
| 37 | *TRAPPC3* | 2.84 | 1.90 | 0.00 |
| 38 | MMP15 | 2.84 | 1.90 | 0.00 |
| 39 | *COL21A1* | 2.81 | 1.89 | 0.00 |
| 40 | NSUN5 | 2.81 | 1.89 | 0.00 |
| 41 | *VWA1* | 2.80 | 1.89 | 0.00 |
| 42 | RHCE | 2.79 | 1.88 | 0.00 |
| 43 | *RAD51L1* | 2.79 | 1.88 | 0.00 |

| | | | | |
|---|---|---|---|---|
| 44 | PRKCSH | 2.79 | 1.88 | 0.00 |
| 45 | *GTF2I* | 2.79 | 1.88 | 0.00 |
| 46 | KCNK5 | 2.78 | 1.88 | 0.00 |
| 47 | *THUMPD1* | 2.78 | 1.88 | 0.00 |
| 48 | CCDC101 | 2.77 | 1.87 | 0.00 |
| 49 | *PDXDC1* | 2.77 | 1.87 | 0.00 |
| 50 | PIGV | 2.75 | 1.87 | 0.00 |
| 51 | *MGRN1* | 2.75 | 1.87 | 0.00 |
| 52 | ZNF263 | 2.74 | 1.86 | 0.00 |
| 53 | *RPA2* | 2.72 | 1.85 | 0.00 |
| 54 | SLC29A1 | 2.72 | 1.85 | 0.00 |
| 55 | *PARK7* | 2.72 | 1.85 | 0.00 |
| 56 | TBL3 | 2.71 | 1.85 | 0.00 |
| 57 | *ZFP36L1* | 2.71 | 1.85 | 0.00 |
| 58 | HMGCL | 2.71 | 1.85 | 0.00 |
| 59 | *PEF1* | 2.71 | 1.85 | 0.00 |
| 60 | GGA2 | 2.70 | 1.85 | 0.00 |
| 61 | *ANKMY2* | 2.70 | 1.84 | 0.00 |
| 62 | FAAH | 2.69 | 1.84 | 0.00 |
| 63 | *LRRC41* | 2.69 | 1.84 | 0.00 |
| 64 | IFT140 | 2.68 | 1.84 | 0.00 |
| 65 | *TELO2* | 2.68 | 1.84 | 0.00 |
| 66 | ATP9A | 2.67 | 1.84 | 0.00 |
| 67 | *HSPB1* | 2.67 | 1.83 | 0.00 |
| 68 | MANBA | 2.67 | 1.83 | 0.00 |
| 69 | *TRMT5* | 2.67 | 1.83 | 0.00 |
| 70 | CTNNBIP1 | 2.67 | 1.83 | 0.00 |
| 71 | *SALL2* | 2.66 | 1.83 | 0.00 |
| 72 | PTK7 | 2.66 | 1.83 | 0.00 |
| 73 | *PRNPIP* | 2.66 | 1.83 | 0.00 |
| 74 | RPS6KA1 | 2.65 | 1.83 | 0.00 |
| 75 | *FUCA1* | 2.64 | 1.82 | 0.00 |
| 76 | WDTC1 | 2.64 | 1.82 | 0.00 |
| 77 | *MR1* | 2.63 | 1.82 | 0.00 |
| 78 | C7orf23 | 2.63 | 1.82 | 0.00 |
| 79 | *LOC643641* | 2.62 | 1.81 | 0.00 |
| 80 | IGSF3 | 2.61 | 1.81 | 0.00 |
| 81 | *VPS13D* | 2.61 | 1.81 | 0.00 |
| 82 | LDOC1 | 2.61 | 1.81 | 0.00 |
| 83 | *OGDH* | 2.61 | 1.81 | 0.00 |
| 84 | BCL7C | 2.60 | 1.81 | 0.00 |
| 85 | *HYI* | 2.60 | 1.81 | 0.00 |
| 86 | SEZ6L2 | 2.60 | 1.81 | 0.00 |
| 87 | *STX4* | 2.60 | 1.81 | 0.00 |
| 88 | H2AFV | 2.60 | 1.81 | 0.00 |

| | | | | |
|---|---|---|---|---|
| 89 | *CAMTA1* | 2.60 | 1.81 | 0.00 |
| 90 | CABIN1 | 2.59 | 1.80 | 0.00 |
| 91 | *LEFTY2* | 2.59 | 1.80 | 0.00 |
| 92 | DDAH1 | 2.59 | 1.80 | 0.00 |
| 93 | *CTA-216E10.6* | 2.58 | 1.80 | 0.00 |
| 94 | NPTXR | 2.58 | 1.80 | 0.00 |
| 95 | *ZBTB48* | 2.57 | 1.80 | 0.00 |
| 96 | ZDHHC11 | 2.55 | 1.79 | 0.00 |
| 97 | *NFYC* | 2.55 | 1.79 | 0.00 |
| 98 | HLA-DMA | 2.54 | 1.79 | 0.00 |
| 99 | *C1orf160* | 2.54 | 1.78 | 0.00 |
| 100 | NUBP2 | 2.54 | 1.78 | 0.00 |
| 101 | *MGC16824* | 2.54 | 1.78 | 0.00 |
| 102 | C14orf94 | 2.53 | 1.78 | 0.00 |
| 103 | *NPC2* | 2.53 | 1.78 | 0.00 |
| 104 | TSPAN13 | 2.53 | 1.78 | 0.00 |
| 105 | *BLVRA* | 2.53 | 1.78 | 0.00 |
| 106 | HSD17B8 | 2.53 | 1.78 | 0.00 |
| 107 | *LYPLA2* | 2.52 | 1.78 | 0.00 |
| 108 | GGTLA4 | 2.52 | 1.78 | 0.00 |
| 109 | *LTBP2* | 2.51 | 1.77 | 0.00 |
| 110 | LTBP3 | 2.51 | 1.77 | 0.00 |
| 111 | *SLC22A18* | 2.50 | 1.77 | 0.00 |
| 112 | DIRAS3 | 2.50 | 1.77 | 0.00 |
| 113 | *TMEM159* | 2.50 | 1.77 | 0.00 |
| 114 | HAGH | 2.50 | 1.77 | 0.00 |
| 115 | *NUDCD3* | 2.50 | 1.77 | 0.00 |
| 116 | AUTS2 | 2.49 | 1.77 | 0.00 |
| 117 | *BCKDK* | 2.49 | 1.77 | 0.00 |
| 118 | CD2BP2 | 2.49 | 1.76 | 0.00 |
| 119 | *SDC1* | 2.48 | 1.76 | 0.00 |
| 120 | PLEKHM1 | 2.48 | 1.76 | 0.00 |
| 121 | *HCFC1R1* | 2.47 | 1.76 | 0.00 |
| 122 | SSBP3 | 2.47 | 1.76 | 0.00 |
| 123 | *DDEFL1* | 2.46 | 1.75 | 0.00 |
| 124 | ZDHHC4 | 2.45 | 1.75 | 0.00 |
| 125 | *TSPAN6* | 2.45 | 1.75 | 0.00 |
| 126 | CA10 | 2.44 | 1.75 | 0.00 |
| 127 | *ZMIZ2* | 2.44 | 1.75 | 0.00 |
| 128 | STOML1 | 2.44 | 1.75 | 0.00 |
| 129 | *NCALD* | 2.44 | 1.75 | 0.00 |
| 130 | MMP24 | 2.44 | 1.75 | 0.00 |
| 131 | *NME3* | 2.44 | 1.75 | 0.00 |
| 132 | ELN | 2.43 | 1.74 | 0.00 |
| 133 | *FOXJ1* | 2.43 | 1.74 | 0.00 |

| | | | | |
|---|---|---|---|---|
| 134 | UROD | 2.43 | 1.74 | 0.00 |
| 135 | *ATP6V0A1* | 2.43 | 1.74 | 0.00 |
| 136 | CPT2 | 2.42 | 1.74 | 0.00 |
| 137 | *EFHC2* | 2.42 | 1.74 | 0.00 |
| 138 | C1orf123 | 2.41 | 1.73 | 0.00 |
| 139 | *CDC42EP1* | 2.40 | 1.73 | 0.00 |
| 140 | MAGED1 | 2.40 | 1.73 | 0.00 |
| 141 | *TMEM39B* | 2.40 | 1.73 | 0.00 |
| 142 | STYXL1 | 2.40 | 1.73 | 0.00 |
| 143 | *APLP2* | 2.40 | 1.73 | 0.00 |
| 144 | ZNF629 | 2.40 | 1.73 | 0.00 |
| 145 | *GBAS* | 2.39 | 1.73 | 0.00 |
| 146 | RHBDF1 | 2.39 | 1.73 | 0.00 |
| 147 | *RNASE1* | 2.39 | 1.73 | 0.00 |
| 148 | GLS2 | 2.39 | 1.73 | 0.00 |
| 149 | *ITPR3* | 2.39 | 1.73 | 0.00 |
| 150 | FMO4 | 2.39 | 1.73 | 0.00 |
| 151 | *MEGF6* | 2.38 | 1.73 | 0.00 |
| 152 | ADAMTSL2 | 2.38 | 1.72 | 0.00 |
| 153 | *FHOD1* | 2.38 | 1.72 | 0.00 |
| 154 | C7orf42 | 2.38 | 1.72 | 0.00 |
| 155 | *SPSB3* | 2.37 | 1.72 | 0.00 |
| 156 | MIR16 | 2.37 | 1.72 | 0.00 |
| 157 | *PUM1* | 2.37 | 1.72 | 0.00 |
| 158 | CES3 | 2.37 | 1.72 | 0.00 |
| 159 | *SLC2A4RG* | 2.37 | 1.72 | 0.00 |
| 160 | GPR116 | 2.37 | 1.72 | 0.00 |
| 161 | *SERPIND1* | 2.37 | 1.72 | 0.00 |
| 162 | PPCS | 2.36 | 1.72 | 0.00 |
| 163 | *RABEP2* | 2.36 | 1.72 | 0.00 |
| 164 | GPBP1L1 | 2.36 | 1.72 | 0.00 |
| 165 | *MRPL20* | 2.36 | 1.72 | 0.00 |
| 166 | APOH | 2.35 | 1.71 | 0.00 |
| 167 | *RAB11FIP3* | 2.35 | 1.71 | 0.00 |
| 168 | SPATA6 | 2.35 | 1.71 | 0.00 |
| 169 | *PHKB* | 2.35 | 1.71 | 0.00 |
| 170 | ORC3L | 2.35 | 1.71 | 0.00 |
| 171 | *ZNF219* | 2.34 | 1.71 | 0.00 |
| 172 | C16orf42 | 2.34 | 1.71 | 0.00 |
| 173 | *SLC15A2* | 2.34 | 1.71 | 0.00 |
| 174 | KIAA0841 | 2.34 | 1.71 | 0.00 |
| 175 | *DECR2* | 2.34 | 1.71 | 0.00 |
| 176 | CLCN7 | 2.33 | 1.71 | 0.00 |
| 177 | *TSC2* | 2.33 | 1.70 | 0.00 |
| 178 | HPCAL4 | 2.33 | 1.70 | 0.00 |

| | | | | |
|---|---|---|---|---|
| 179 | *MYST1* | 2.32 | 1.70 | 0.00 |
| 180 | CREBBP | 2.32 | 1.70 | 0.00 |
| 181 | *MVP* | 2.32 | 1.70 | 0.00 |
| 182 | GPC4 | 2.32 | 1.70 | 0.00 |
| 183 | *CADPS2* | 2.32 | 1.70 | 0.00 |
| 184 | THRA | 2.32 | 1.70 | 0.00 |
| 185 | *FLJ10781* | 2.31 | 1.70 | 0.00 |
| 186 | NUBP1 | 2.31 | 1.70 | 0.00 |
| 187 | *NCDN* | 2.31 | 1.70 | 0.00 |
| 188 | SLC9A1 | 2.31 | 1.70 | 0.00 |
| 189 | *AKR7A2* | 2.30 | 1.70 | 0.00 |
| 190 | PKD1 | 2.30 | 1.70 | 0.00 |
| 191 | *MALL* | 2.30 | 1.70 | 0.00 |
| 192 | CDIPT | 2.30 | 1.70 | 0.00 |
| 193 | *LCT* | 2.30 | 1.69 | 0.00 |
| 194 | PRELP | 2.30 | 1.69 | 0.00 |
| 195 | *ROR1* | 2.30 | 1.69 | 0.00 |
| 196 | CST5 | 2.29 | 1.69 | 0.00 |
| 197 | *SF3A3* | 2.29 | 1.69 | 0.00 |
| 198 | UBN1 | 2.28 | 1.69 | 0.00 |
| 199 | *CCT6B* | 2.28 | 1.69 | 0.00 |
| 200 | ETV5 | 2.28 | 1.69 | 0.00 |
| 201 | *FZD1* | 2.27 | 1.68 | 0.00 |
| 202 | ALPL | 2.27 | 1.68 | 0.00 |
| 203 | *CTSH* | 2.27 | 1.68 | 0.00 |
| 204 | DPY19L1 | 2.27 | 1.68 | 0.00 |
| 205 | *TMEM112* | 2.27 | 1.68 | 0.00 |
| 206 | ETNK2 | 2.26 | 1.68 | 0.00 |
| 207 | *COMP* | 2.26 | 1.68 | 0.00 |
| 208 | PCSK1N | 2.26 | 1.68 | 0.00 |
| 209 | *GATAD1* | 2.26 | 1.68 | 0.00 |
| 210 | UBE2D4 | 2.26 | 1.68 | 0.00 |
| 211 | *ARMCX6* | 2.25 | 1.68 | 0.00 |
| 212 | ABCA4 | 2.25 | 1.68 | 0.00 |
| 213 | *CEBPA* | 2.25 | 1.68 | 0.00 |
| 214 | NARFL | 2.25 | 1.67 | 0.00 |
| 215 | *DPP4* | 2.25 | 1.67 | 0.00 |
| 216 | CPSF3L | 2.24 | 1.67 | 0.00 |
| 217 | *PRSS16* | 2.24 | 1.67 | 0.00 |
| 218 | TRAPPC2 | 2.24 | 1.67 | 0.00 |
| 219 | *RNPS1* | 2.24 | 1.67 | 0.00 |
| 220 | ZNF500 | 2.23 | 1.67 | 0.00 |
| 221 | *SDF4* | 2.23 | 1.67 | 0.00 |
| 222 | CLDN4 | 2.23 | 1.67 | 0.00 |
| 223 | *EXOD1* | 2.23 | 1.67 | 0.00 |

| | | | | |
|---|---|---|---|---|
| 224 | PER3 | 2.23 | 1.67 | 0.00 |
| 225 | *LGALS3BP* | 2.22 | 1.67 | 0.00 |
| 226 | C16orf35 | 2.22 | 1.66 | 0.00 |
| 227 | *HMGN2* | 2.22 | 1.66 | 0.00 |
| 228 | DDR1 | 2.21 | 1.66 | 0.00 |
| 229 | *URG4* | 2.21 | 1.66 | 0.00 |
| 230 | RCP9 | 2.21 | 1.66 | 0.00 |
| 231 | *BAZ1B* | 2.21 | 1.66 | 0.00 |
| 232 | MYO1D | 2.21 | 1.66 | 0.00 |
| 233 | *METRN* | 2.20 | 1.66 | 0.00 |
| 234 | BSDC1 | 2.20 | 1.66 | 0.00 |
| 235 | *FASTK* | 2.20 | 1.66 | 0.00 |
| 236 | GDPD5 | 2.20 | 1.66 | 0.00 |
| 237 | *ATP6V0E2* | 2.20 | 1.66 | 0.00 |
| 238 | LIMK1 | 2.19 | 1.66 | 0.00 |
| 239 | *TAOK2* | 2.19 | 1.66 | 0.00 |
| 240 | SFRP4 | 2.19 | 1.66 | 0.00 |
| 241 | *PDPK1* | 2.19 | 1.65 | 0.00 |
| 242 | KIAA1305 | 2.19 | 1.65 | 0.00 |
| 243 | *CLEC16A* | 2.19 | 1.65 | 0.00 |
| 244 | CD207 | 2.19 | 1.65 | 0.00 |
| 245 | *PHACTR4* | 2.18 | 1.65 | 0.00 |
| 246 | AKR1A1 | 2.17 | 1.65 | 0.00 |
| 247 | *ZNF107* | 2.17 | 1.65 | 0.00 |
| 248 | ARSD | 2.17 | 1.65 | 0.00 |
| 249 | *DCLK1* | 2.17 | 1.65 | 0.00 |
| 250 | APOD | 2.16 | 1.64 | 0.00 |
| 251 | *FOLR1* | 2.16 | 1.64 | 0.00 |
| 252 | YIPF2 | 2.16 | 1.64 | 0.00 |
| 253 | *C16orf5* | 2.15 | 1.64 | 0.00 |
| 254 | RGL1 | 2.15 | 1.64 | 0.00 |
| 255 | *POLR2C* | 2.15 | 1.64 | 0.00 |
| 256 | C16orf53 | 2.15 | 1.64 | 0.00 |
| 257 | *RRAD* | 2.15 | 1.64 | 0.00 |
| 258 | MAP3K13 | 2.15 | 1.64 | 0.00 |
| 259 | *ORAI3* | 2.14 | 1.64 | 0.00 |
| 260 | HOXD1 | 2.14 | 1.63 | 0.00 |
| 261 | *CST2* | 2.14 | 1.63 | 0.00 |
| 262 | ARID1A | 2.13 | 1.63 | 0.00 |
| 263 | *SCP2* | 2.13 | 1.63 | 0.00 |
| 264 | BBS9 | 2.13 | 1.63 | 0.00 |
| 265 | *ARHGEF10L* | 2.13 | 1.63 | 0.00 |
| 266 | TOMM7 | 2.12 | 1.63 | 0.00 |
| 267 | *IPO13* | 2.12 | 1.63 | 0.00 |
| 268 | KPNA6 | 2.12 | 1.63 | 0.00 |

| | | | | |
|---|---|---|---|---|
| 269 | *ADCK2* | 2.12 | 1.63 | 0.00 |
| 270 | TRIOBP | 2.12 | 1.63 | 0.00 |
| 271 | *CLDN3* | 2.11 | 1.63 | 0.00 |
| 272 | VKORC1 | 2.11 | 1.63 | 0.00 |
| 273 | *OSGEP* | 2.11 | 1.63 | 0.00 |
| 274 | STUB1 | 2.11 | 1.63 | 0.00 |
| 275 | *NTHL1* | 2.10 | 1.62 | 0.00 |
| 276 | FKBPL | 2.10 | 1.62 | 0.00 |
| 277 | *TYRP1* | 2.10 | 1.62 | 0.00 |
| 278 | TNNI3 | 2.10 | 1.62 | 0.00 |
| 279 | *FLJ10357* | 2.10 | 1.62 | 0.00 |
| 280 | MPZL2 | 2.10 | 1.62 | 0.00 |
| 281 | *BCAM* | 2.09 | 1.62 | 0.00 |
| 282 | RHOT2 | 2.09 | 1.62 | 0.00 |
| 283 | *FAM3A* | 2.09 | 1.62 | 0.00 |
| 284 | ELOVL1 | 2.09 | 1.62 | 0.00 |
| 285 | *APITD1* | 2.09 | 1.62 | 0.00 |
| 286 | UNC84A | 2.09 | 1.62 | 0.00 |
| 287 | *DAAM1* | 2.08 | 1.62 | 0.00 |
| 288 | NPAL3 | 2.08 | 1.62 | 0.00 |
| 289 | *PHKG2* | 2.08 | 1.61 | 0.00 |
| 290 | ZNF34 | 2.07 | 1.61 | 0.00 |
| 291 | *C1orf116* | 2.07 | 1.61 | 0.00 |
| 292 | HN1L | 2.07 | 1.61 | 0.00 |
| 293 | *ABCC6* | 2.07 | 1.61 | 0.00 |
| 294 | SCUBE2 | 2.06 | 1.61 | 0.00 |
| 295 | *AGRN* | 2.06 | 1.61 | 0.00 |
| 296 | FGF18 | 2.06 | 1.61 | 0.00 |
| 297 | *PHF1* | 2.06 | 1.61 | 0.00 |
| 298 | RAC1 | 2.06 | 1.61 | 0.00 |
| 299 | *CLUAP1* | 2.06 | 1.61 | 0.00 |
| 300 | WDR91 | 2.06 | 1.61 | 0.00 |
| 301 | *PCTK3* | 2.06 | 1.61 | 0.00 |
| 302 | ZNF212 | 2.05 | 1.61 | 0.00 |
| 303 | *COL10A1* | 2.05 | 1.61 | 0.00 |
| 304 | TMEM50A | 2.05 | 1.60 | 0.00 |
| 305 | *ICMT* | 2.05 | 1.60 | 0.00 |
| 306 | STEAP3 | 2.05 | 1.60 | 0.00 |
| 307 | *TMED9* | 2.05 | 1.60 | 0.00 |
| 308 | STK19 | 2.05 | 1.60 | 0.00 |
| 309 | *ABHD11* | 2.05 | 1.60 | 0.00 |
| 310 | RER1 | 2.05 | 1.60 | 0.00 |
| 311 | *EPB41L1* | 2.04 | 1.60 | 0.00 |
| 312 | ZNF764 | 2.04 | 1.60 | 0.00 |
| 313 | *NINJ2* | 2.04 | 1.60 | 0.00 |

| | | | | |
|---|---|---|---|---|
| 314 | CD1E | 2.04 | 1.60 | 0.00 |
| 315 | *SOX13* | 2.04 | 1.60 | 0.00 |
| 316 | C5orf3 | 2.04 | 1.60 | 0.00 |
| 317 | *GCDH* | 2.04 | 1.60 | 0.00 |
| 318 | TCEB3 | 2.04 | 1.60 | 0.00 |
| 319 | *CD74* | 2.04 | 1.60 | 0.00 |
| 320 | BAIAP3 | 2.03 | 1.60 | 0.00 |
| 321 | *USP7* | 2.03 | 1.60 | 0.00 |
| 322 | SCMH1 | 2.03 | 1.60 | 0.00 |
| 323 | *MOSPD3* | 2.03 | 1.60 | 0.00 |
| 324 | MAPK8IP3 | 2.03 | 1.60 | 0.00 |
| 325 | *MPG* | 2.03 | 1.60 | 0.00 |
| 326 | TLR2 | 2.03 | 1.60 | 0.00 |
| 327 | *ZMYM3* | 2.03 | 1.60 | 0.00 |
| 328 | GJB1 | 2.03 | 1.60 | 0.00 |
| 329 | *CLSTN1* | 2.03 | 1.60 | 0.00 |
| 330 | ATP7A | 2.03 | 1.60 | 0.00 |
| 331 | *TMEM63A* | 2.03 | 1.60 | 0.00 |
| 332 | TMEM112B | 2.02 | 1.59 | 0.00 |
| 333 | *KIAA0240* | 2.02 | 1.59 | 0.00 |
| 334 | MTCH1 | 2.02 | 1.59 | 0.00 |
| 335 | *ARHGEF9* | 2.02 | 1.59 | 0.00 |
| 336 | PNPLA4 | 2.02 | 1.59 | 0.00 |
| 337 | *SFRS4* | 2.02 | 1.59 | 0.00 |
| 338 | GRB14 | 2.02 | 1.59 | 0.00 |
| 339 | *SARM1* | 2.01 | 1.59 | 0.00 |
| 340 | PLA2G1B | 2.01 | 1.59 | 0.00 |
| 341 | *NUDC* | 2.01 | 1.59 | 0.00 |
| 342 | PDK2 | 2.00 | 1.59 | 0.00 |
| 343 | *CLDN9* | 2.00 | 1.59 | 0.00 |
| 344 | ACAD8 | 2.00 | 1.59 | 0.00 |
| 345 | *GFER* | 2.00 | 1.59 | 0.00 |
| 346 | CXorf56 | 2.00 | 1.59 | 0.00 |
| 347 | *CEACAM4* | 2.00 | 1.59 | 0.00 |
| 348 | RAPGEF5 | 2.00 | 1.58 | 0.00 |
| 349 | *MRPS18B* | 1.99 | 1.58 | 0.00 |
| 350 | ZNF354A | 1.99 | 1.58 | 0.00 |
| 351 | *C19orf56* | 1.99 | 1.58 | 0.00 |
| 352 | C18orf1 | 1.99 | 1.58 | 0.00 |
| 353 | *POU2F3* | 1.99 | 1.58 | 0.00 |
| 354 | PRSS8 | 1.99 | 1.58 | 0.00 |
| 355 | *SGSH* | 1.99 | 1.58 | 0.00 |
| 356 | WWP2 | 1.99 | 1.58 | 0.00 |
| 357 | *WBSCR22* | 1.99 | 1.58 | 0.00 |
| 358 | TRIM26 | 1.99 | 1.58 | 0.00 |

| | | | | |
|---|---|---|---|---|
| 359 | *AEBP1* | 1.99 | 1.58 | 0.00 |
| 360 | CUEDC1 | 1.99 | 1.58 | 0.00 |
| 361 | *THOC6* | 1.98 | 1.58 | 0.00 |
| 362 | MGP | 1.98 | 1.58 | 0.00 |
| 363 | *TCFL5* | 1.98 | 1.58 | 0.13 |
| 364 | REV1 | 1.98 | 1.58 | 0.13 |
| 365 | *KIAA0467* | 1.98 | 1.58 | 0.13 |
| 366 | ITGA9 | 1.98 | 1.58 | 0.13 |
| 367 | *UBTD1* | 1.98 | 1.58 | 0.13 |
| 368 | AHR | 1.98 | 1.58 | 0.13 |
| 369 | *PPP1R3C* | 1.98 | 1.58 | 0.13 |
| 370 | GALNT11 | 1.97 | 1.58 | 0.13 |
| 371 | *PGRMC1* | 1.97 | 1.58 | 0.13 |
| 372 | WDR13 | 1.97 | 1.58 | 0.13 |
| 373 | *IPP* | 1.97 | 1.58 | 0.13 |
| 374 | DOK5 | 1.97 | 1.57 | 0.13 |
| 375 | *RRN3* | 1.96 | 1.57 | 0.13 |
| 376 | ALG12 | 1.96 | 1.57 | 0.13 |
| 377 | *HLA-DPB1* | 1.96 | 1.57 | 0.13 |
| 378 | HSPB7 | 1.96 | 1.57 | 0.13 |
| 379 | *CDH1* | 1.96 | 1.57 | 0.13 |
| 380 | MUTYH | 1.96 | 1.57 | 0.13 |
| 381 | *F11* | 1.96 | 1.57 | 0.13 |
| 382 | HLA-DPA1 | 1.95 | 1.57 | 0.13 |
| 383 | *UXS1* | 1.95 | 1.57 | 0.13 |
| 384 | ZFHX3 | 1.95 | 1.57 | 0.13 |
| 385 | *LOC81691* | 1.95 | 1.57 | 0.13 |
| 386 | AMOT | 1.95 | 1.57 | 0.13 |
| 387 | *TP53AP1* | 1.95 | 1.57 | 0.13 |
| 388 | CALCOCO2 | 1.95 | 1.57 | 0.13 |
| 389 | *AGPAT1* | 1.94 | 1.57 | 0.13 |
| 390 | C1orf50 | 1.94 | 1.57 | 0.13 |
| 391 | *FLJ22222* | 1.94 | 1.57 | 0.13 |
| 392 | GPR137 | 1.94 | 1.57 | 0.13 |
| 393 | *IL27RA* | 1.94 | 1.56 | 0.13 |
| 394 | CASD1 | 1.94 | 1.56 | 0.13 |
| 395 | *NOD1* | 1.93 | 1.56 | 0.13 |
| 396 | VPS41 | 1.93 | 1.56 | 0.13 |
| 397 | *FBXO42* | 1.93 | 1.56 | 0.13 |
| 398 | ST7 | 1.93 | 1.56 | 0.13 |
| 399 | *COL8A2* | 1.93 | 1.56 | 0.13 |
| 400 | GSPT1 | 1.92 | 1.56 | 0.13 |
| 401 | *HAND1* | 1.92 | 1.56 | 0.13 |
| 402 | ZNF12 | 1.92 | 1.56 | 0.13 |
| 403 | *GGCX* | 1.92 | 1.56 | 0.13 |

| | | | | |
|---|---|---|---|---|
| 404 | C7orf26 | 1.92 | 1.56 | 0.13 |
| 405 | *GBL* | 1.92 | 1.56 | 0.13 |
| 406 | SPINT2 | 1.92 | 1.56 | 0.13 |
| 407 | *CNNM3* | 1.92 | 1.56 | 0.13 |
| 408 | ZMYM6 | 1.92 | 1.56 | 0.13 |
| 409 | *ECHDC2* | 1.92 | 1.56 | 0.13 |
| 410 | HSD11B2 | 1.91 | 1.56 | 0.13 |
| 411 | *SKIV2L* | 1.91 | 1.56 | 0.13 |
| 412 | NUPL2 | 1.91 | 1.56 | 0.13 |
| 413 | *STK17A* | 1.91 | 1.55 | 0.13 |
| 414 | ENC1 | 1.91 | 1.55 | 0.13 |
| 415 | *AZGP1* | 1.91 | 1.55 | 0.13 |
| 416 | LRRC20 | 1.91 | 1.55 | 0.13 |
| 417 | *SLC13A3* | 1.91 | 1.55 | 0.13 |
| 418 | SH3BGR | 1.91 | 1.55 | 0.13 |
| 419 | *IFI6* | 1.91 | 1.55 | 0.13 |
| 420 | ARAF | 1.90 | 1.55 | 0.13 |
| 421 | *USP11* | 1.90 | 1.55 | 0.13 |
| 422 | GRAMD3 | 1.90 | 1.55 | 0.13 |
| 423 | *CLN3* | 1.89 | 1.55 | 0.13 |
| 424 | TMEM186 | 1.89 | 1.55 | 0.13 |
| 425 | *RABL4* | 1.89 | 1.55 | 0.13 |
| 426 | DIO1 | 1.89 | 1.55 | 0.13 |
| 427 | *SULT1C2* | 1.89 | 1.55 | 0.13 |
| 428 | WIPI2 | 1.89 | 1.55 | 0.13 |
| 429 | *UBE2I* | 1.89 | 1.55 | 0.30 |
| 430 | PARN | 1.88 | 1.55 | 0.30 |
| 431 | *CRYM* | 1.88 | 1.55 | 0.30 |
| 432 | TBC1D10B | 1.88 | 1.54 | 0.30 |
| 433 | *ACTB* | 1.88 | 1.54 | 0.30 |
| 434 | MFN2 | 1.88 | 1.54 | 0.30 |
| 435 | *GGT1* | 1.88 | 1.54 | 0.30 |
| 436 | MGC10334 | 1.88 | 1.54 | 0.30 |
| 437 | *TBL2* | 1.88 | 1.54 | 0.30 |
| 438 | TXLNA | 1.88 | 1.54 | 0.30 |
| 439 | *BSCL2* | 1.88 | 1.54 | 0.30 |
| 440 | RASGRF1 | 1.88 | 1.54 | 0.30 |
| 441 | *C7orf10* | 1.87 | 1.54 | 0.30 |
| 442 | SLC2A10 | 1.87 | 1.54 | 0.30 |
| 443 | *FCGBP* | 1.87 | 1.54 | 0.30 |
| 444 | TCEB2 | 1.87 | 1.54 | 0.30 |
| 445 | *C20orf103* | 1.87 | 1.54 | 0.30 |
| 446 | DAP | 1.87 | 1.54 | 0.30 |
| 447 | *EIF2C1* | 1.87 | 1.54 | 0.30 |
| 448 | RAP1GAP | 1.87 | 1.54 | 0.30 |

| | | | | |
|---|---|---|---|---|
| 449 | *PIB5PA* | 1.86 | 1.54 | 0.30 |
| 450 | WBSCR16 | 1.86 | 1.54 | 0.30 |
| 451 | *DOC2A* | 1.86 | 1.54 | 0.30 |
| 452 | RNF216 | 1.86 | 1.54 | 0.30 |
| 453 | *LTBP1* | 1.86 | 1.54 | 0.30 |
| 454 | RORB | 1.86 | 1.54 | 0.30 |
| 455 | *PPT1* | 1.86 | 1.54 | 0.30 |
| 456 | ZNF434 | 1.86 | 1.54 | 0.30 |
| 457 | *ARHGDIB* | 1.85 | 1.53 | 0.30 |
| 458 | FAF1 | 1.85 | 1.53 | 0.30 |
| 459 | *SUSD4* | 1.85 | 1.53 | 0.30 |
| 460 | AP4S1 | 1.85 | 1.53 | 0.30 |
| 461 | *PPL* | 1.85 | 1.53 | 0.30 |
| 462 | MYOZ1 | 1.85 | 1.53 | 0.30 |
| 463 | *DNAJA3* | 1.85 | 1.53 | 0.30 |
| 464 | RIMS3 | 1.84 | 1.53 | 0.30 |
| 465 | *NOTCH2* | 1.84 | 1.53 | 0.30 |
| 466 | CYB5R1 | 1.84 | 1.53 | 0.30 |
| 467 | *MRPS34* | 1.84 | 1.53 | 0.30 |
| 468 | RING1 | 1.84 | 1.53 | 0.30 |
| 469 | *TLE2* | 1.84 | 1.53 | 0.30 |
| 470 | CD1A | 1.84 | 1.53 | 0.30 |
| 471 | *DLK1* | 1.84 | 1.53 | 0.30 |
| 472 | THTPA | 1.83 | 1.53 | 0.30 |
| 473 | *B3GNT1* | 1.83 | 1.53 | 0.30 |
| 474 | SHROOM2 | 1.83 | 1.53 | 0.30 |
| 475 | *KIAA1026* | 1.83 | 1.53 | 0.30 |
| 476 | SIRT3 | 1.83 | 1.53 | 0.30 |
| 477 | *WFDC2* | 1.82 | 1.53 | 0.30 |
| 478 | PINK1 | 1.82 | 1.52 | 0.30 |
| 479 | *TRIM24* | 1.82 | 1.52 | 0.30 |
| 480 | CD82 | 1.82 | 1.52 | 0.30 |
| 481 | *BBS1* | 1.82 | 1.52 | 0.30 |
| 482 | HHLA2 | 1.82 | 1.52 | 0.30 |
| 483 | *PLXNA2* | 1.82 | 1.52 | 0.30 |
| 484 | BTG3 | 1.82 | 1.52 | 0.30 |
| 485 | *MED8* | 1.82 | 1.52 | 0.30 |
| 486 | WDR8 | 1.81 | 1.52 | 0.30 |
| 487 | *MAGED2* | 1.81 | 1.52 | 0.30 |
| 488 | NSUN5C | 1.81 | 1.52 | 0.30 |
| 489 | *ABCB8* | 1.81 | 1.52 | 0.30 |
| 490 | BNIP1 | 1.81 | 1.52 | 0.30 |
| 491 | *DHX58* | 1.81 | 1.52 | 0.30 |
| 492 | HNF1B | 1.81 | 1.52 | 0.30 |
| 493 | *PTPRF* | 1.81 | 1.52 | 0.30 |

| | | | | |
|---|---|---|---|---|
| 494 | KRIT1 | 1.81 | 1.52 | 0.30 |
| 495 | *S100A13* | 1.80 | 1.52 | 0.30 |
| 496 | CDR2 | 1.80 | 1.52 | 0.30 |
| 497 | *JUN* | 1.80 | 1.52 | 0.40 |
| 498 | FLJ20920 | 1.80 | 1.52 | 0.40 |
| 499 | *PPP1R9A* | 1.80 | 1.52 | 0.40 |
| 500 | C6orf47 | 1.80 | 1.52 | 0.40 |
| 501 | *PRKD1* | 1.80 | 1.52 | 0.40 |
| 502 | GPC3 | 1.80 | 1.52 | 0.40 |
| 503 | *WWC1* | 1.80 | 1.52 | 0.40 |
| 504 | ATAD4 | 1.80 | 1.52 | 0.40 |
| 505 | *SYNPO* | 1.79 | 1.52 | 0.40 |
| 506 | RAB38 | 1.79 | 1.51 | 0.40 |
| 507 | *IFNGR2* | 1.79 | 1.51 | 0.40 |
| 508 | MYH2 | 1.79 | 1.51 | 0.40 |
| 509 | *FBXW11* | 1.79 | 1.51 | 0.40 |
| 510 | LBA1 | 1.79 | 1.51 | 0.40 |
| 511 | *USP13* | 1.79 | 1.51 | 0.40 |
| 512 | CAPZB | 1.79 | 1.51 | 0.40 |
| 513 | *FLJ20323* | 1.79 | 1.51 | 0.40 |
| 514 | PEX14 | 1.78 | 1.51 | 0.40 |
| 515 | *TAX1BP1* | 1.78 | 1.51 | 0.40 |
| 516 | CXCL14 | 1.78 | 1.51 | 0.40 |
| 517 | *EMP2* | 1.78 | 1.51 | 0.40 |
| 518 | DNAJC4 | 1.78 | 1.51 | 0.40 |
| 519 | *C7orf24* | 1.78 | 1.51 | 0.40 |
| 520 | SLC12A9 | 1.78 | 1.51 | 0.40 |
| 521 | *NDUFB2* | 1.78 | 1.51 | 0.40 |
| 522 | THYN1 | 1.77 | 1.51 | 0.40 |
| 523 | *ZNF768* | 1.77 | 1.51 | 0.40 |
| 524 | PDGFD | 1.77 | 1.51 | 0.40 |
| 525 | *SYT17* | 1.77 | 1.51 | 0.40 |
| 526 | HLA-DRA | 1.77 | 1.51 | 0.40 |
| 527 | *NKX2-1* | 1.77 | 1.51 | 0.40 |
| 528 | NUMA1 | 1.77 | 1.51 | 0.40 |
| 529 | *UBB* | 1.76 | 1.51 | 0.40 |
| 530 | PRODH | 1.76 | 1.50 | 0.40 |
| 531 | *PXMP4* | 1.76 | 1.50 | 0.40 |
| 532 | EXPH5 | 1.76 | 1.50 | 0.40 |
| 533 | *AURKAIP1* | 1.76 | 1.50 | 0.40 |
| 534 | TCEAL4 | 1.76 | 1.50 | 0.40 |
| 535 | *E4F1* | 1.76 | 1.50 | 0.40 |
| 536 | MICALL2 | 1.76 | 1.50 | 0.40 |
| 537 | *KIAA0247* | 1.76 | 1.50 | 0.40 |
| 538 | BEX4 | 1.76 | 1.50 | 0.40 |

| | | | | |
|---|---|---|---|---|
| 539 | *SLC1A7* | 1.75 | 1.50 | 0.40 |
| 540 | MXRA8 | 1.75 | 1.50 | 0.40 |
| 541 | *ZNF174* | 1.75 | 1.50 | 0.40 |
| 542 | HLA-DMB | 1.75 | 1.50 | 0.40 |
| 543 | *ZNF673* | 1.75 | 1.50 | 0.40 |
| 544 | GFRA3 | 1.75 | 1.50 | 0.40 |
| 545 | *CSDC2* | 1.75 | 1.50 | 0.40 |
| 546 | EIF4H | 1.75 | 1.50 | 0.40 |
| 547 | *CNKSR1* | 1.75 | 1.50 | 0.40 |
| 548 | HSPG2 | 1.75 | 1.50 | 0.40 |
| 549 | *TRADD* | 1.75 | 1.50 | 0.40 |
| 550 | BGN | 1.74 | 1.50 | 0.40 |
| 551 | *PDE9A* | 1.74 | 1.50 | 0.40 |
| 552 | HLA-DOA | 1.74 | 1.50 | 0.40 |
| 553 | *TACSTD2* | 1.74 | 1.50 | 0.40 |
| 554 | XAB2 | 1.74 | 1.50 | 0.40 |
| 555 | *SRCAP* | 1.74 | 1.50 | 0.40 |
| 556 | PEX3 | 1.74 | 1.50 | 0.40 |
| 557 | *ZNF702* | 1.74 | 1.50 | 0.40 |
| 558 | FOXJ3 | 1.74 | 1.50 | 0.40 |
| 559 | *CROCC* | 1.73 | 1.49 | 0.40 |
| 560 | DHDDS | 1.73 | 1.49 | 0.40 |
| 561 | *LRRC36* | 1.73 | 1.49 | 0.40 |
| 562 | GDF11 | 1.73 | 1.49 | 0.40 |
| 563 | *F8A1* | 1.73 | 1.49 | 0.40 |
| 564 | CBX6 | 1.73 | 1.49 | 0.40 |
| 565 | *C1orf66* | 1.72 | 1.49 | 0.40 |
| 566 | NELL1 | 1.72 | 1.49 | 0.40 |
| 567 | *PSMC2* | 1.72 | 1.49 | 0.40 |
| 568 | LOC89944 | 1.72 | 1.49 | 0.40 |
| 569 | *SRRM1* | 1.72 | 1.49 | 0.40 |
| 570 | MBIP | 1.72 | 1.49 | 0.40 |
| 571 | *PKD2* | 1.72 | 1.49 | 0.40 |
| 572 | NASP | 1.72 | 1.49 | 0.40 |
| 573 | *VTCN1* | 1.72 | 1.49 | 0.66 |
| 574 | MMP28 | 1.71 | 1.49 | 0.66 |
| 575 | *C11orf61* | 1.71 | 1.49 | 0.66 |
| 576 | NRIP3 | 1.71 | 1.49 | 0.66 |
| 577 | *ADORA1* | 1.71 | 1.49 | 0.66 |
| 578 | ABCD3 | 1.71 | 1.49 | 0.66 |
| 579 | *TRPM4* | 1.71 | 1.49 | 0.66 |
| 580 | CETN2 | 1.71 | 1.49 | 0.66 |
| 581 | *NPTX2* | 1.71 | 1.49 | 0.66 |
| 582 | SNX13 | 1.71 | 1.49 | 0.66 |
| 583 | *AP2A2* | 1.71 | 1.49 | 0.66 |

| | | | | |
|---|---|---|---|---|
| 584 | C6orf60 | 1.71 | 1.49 | 0.66 |
| 585 | *PIK3R3* | 1.70 | 1.48 | 0.66 |
| 586 | ZNF302 | 1.70 | 1.48 | 0.66 |
| 587 | *WDR42A* | 1.70 | 1.48 | 0.66 |
| 588 | AP1M2 | 1.70 | 1.48 | 0.66 |
| 589 | *CCDC121* | 1.70 | 1.48 | 0.66 |
| 590 | TNFSF10 | 1.70 | 1.48 | 0.66 |
| 591 | *VPS52* | 1.70 | 1.48 | 0.66 |
| 592 | TRIM68 | 1.70 | 1.48 | 0.66 |
| 593 | *GNAS* | 1.69 | 1.48 | 0.66 |
| 594 | CNNM1 | 1.69 | 1.48 | 0.66 |
| 595 | *ECM1* | 1.69 | 1.48 | 0.66 |
| 596 | ROS1 | 1.69 | 1.48 | 0.66 |
| 597 | *DAAM2* | 1.69 | 1.48 | 0.66 |
| 598 | TNFRSF12A | 1.69 | 1.48 | 0.66 |
| 599 | *PFKL* | 1.69 | 1.48 | 0.66 |
| 600 | EN2 | 1.69 | 1.48 | 0.66 |
| 601 | *ANPEP* | 1.69 | 1.48 | 0.66 |
| 602 | CHST12 | 1.69 | 1.48 | 0.66 |
| 603 | *FGF13* | 1.69 | 1.48 | 0.66 |
| 604 | C20orf149 | 1.69 | 1.48 | 0.66 |
| 605 | *FLJ23861* | 1.68 | 1.48 | 0.66 |
| 606 | FAM90A1 | 1.68 | 1.48 | 0.66 |
| 607 | *SSPN* | 1.68 | 1.48 | 0.66 |
| 608 | ABR | 1.68 | 1.48 | 0.66 |
| 609 | *ROM1* | 1.68 | 1.48 | 0.66 |
| 610 | ATPBD1B | 1.68 | 1.48 | 0.66 |
| 611 | *CRYZ* | 1.68 | 1.48 | 0.66 |
| 612 | C1orf109 | 1.68 | 1.48 | 0.66 |
| 613 | *CD1C* | 1.68 | 1.48 | 0.66 |
| 614 | LAPTM4A | 1.68 | 1.48 | 0.66 |
| 615 | *CRABP2* | 1.68 | 1.48 | 0.66 |
| 616 | IFI44 | 1.68 | 1.48 | 0.66 |
| 617 | *DKK3* | 1.68 | 1.48 | 0.66 |
| 618 | RBM23 | 1.68 | 1.48 | 0.66 |
| 619 | *GCAT* | 1.68 | 1.48 | 0.66 |
| 620 | RANBP17 | 1.67 | 1.47 | 0.66 |
| 621 | *RBM9* | 1.67 | 1.47 | 0.66 |
| 622 | BCORL1 | 1.67 | 1.47 | 0.66 |
| 623 | *MDK* | 1.67 | 1.47 | 0.66 |
| 624 | SPRED2 | 1.67 | 1.47 | 0.66 |
| 625 | *DOK4* | 1.67 | 1.47 | 0.66 |
| 626 | LSP1 | 1.67 | 1.47 | 0.66 |
| 627 | *TUFM* | 1.67 | 1.47 | 0.66 |
| 628 | PGCP | 1.66 | 1.47 | 0.66 |

| | | | | |
|---|---|---|---|---|
| 629 | *FOXRED2* | 1.66 | 1.47 | 0.66 |
| 630 | C11orf68 | 1.66 | 1.47 | 0.66 |
| 631 | *IDS* | 1.66 | 1.47 | 0.66 |
| 632 | C9orf127 | 1.66 | 1.47 | 0.66 |
| 633 | *FAM134C* | 1.66 | 1.47 | 0.66 |
| 634 | TNFSF15 | 1.66 | 1.47 | 0.66 |
| 635 | *SLC17A3* | 1.66 | 1.47 | 0.66 |
| 636 | DEXI | 1.66 | 1.47 | 0.66 |
| 637 | *ST3GAL6* | 1.66 | 1.47 | 0.66 |
| 638 | HLA-A | 1.66 | 1.47 | 0.66 |
| 639 | *HOXC4* | 1.66 | 1.47 | 0.66 |
| 640 | MYH10 | 1.66 | 1.47 | 0.66 |
| 641 | *CAPN2* | 1.66 | 1.47 | 0.66 |
| 642 | PIK3IP1 | 1.66 | 1.47 | 0.66 |
| 643 | *TMEM8* | 1.66 | 1.47 | 0.66 |
| 644 | NPHP4 | 1.65 | 1.47 | 0.66 |
| 645 | *KIAA0644* | 1.65 | 1.47 | 0.66 |
| 646 | ICAM1 | 1.65 | 1.47 | 0.66 |
| 647 | *SPOP* | 1.65 | 1.47 | 0.66 |
| 648 | NFIX | 1.65 | 1.47 | 0.66 |
| 649 | *PHC2* | 1.65 | 1.47 | 0.66 |
| 650 | KIF22 | 1.65 | 1.47 | 0.66 |
| 651 | *C14orf133* | 1.65 | 1.47 | 0.66 |
| 652 | MACROD1 | 1.65 | 1.47 | 0.66 |
| 653 | *B3GAT1* | 1.65 | 1.47 | 0.66 |
| 654 | KIAA0082 | 1.65 | 1.47 | 0.66 |
| 655 | *SIDT1* | 1.65 | 1.47 | 0.66 |
| 656 | TMEM59 | 1.65 | 1.47 | 0.66 |
| 657 | *RPS6KA3* | 1.64 | 1.47 | 0.66 |
| 658 | ZNF43 | 1.64 | 1.46 | 0.66 |
| 659 | *ITGBL1* | 1.64 | 1.46 | 0.66 |
| 660 | IL10RB | 1.64 | 1.46 | 0.66 |
| 661 | *SLC4A5* | 1.64 | 1.46 | 0.66 |
| 662 | KIAA0562 | 1.64 | 1.46 | 0.66 |
| 663 | *TSPAN4* | 1.64 | 1.46 | 0.66 |
| 664 | SELENBP1 | 1.64 | 1.46 | 0.66 |
| 665 | *MATN3* | 1.64 | 1.46 | 0.66 |
| 666 | SLC35E1 | 1.64 | 1.46 | 0.66 |
| 667 | *FABP3* | 1.64 | 1.46 | 0.66 |
| 668 | POMGNT1 | 1.64 | 1.46 | 0.66 |
| 669 | *ZNF137* | 1.64 | 1.46 | 0.66 |
| 670 | TMPRSS2 | 1.64 | 1.46 | 0.66 |
| 671 | *AK1* | 1.64 | 1.46 | 0.66 |
| 672 | ADAT1 | 1.64 | 1.46 | 0.66 |
| 673 | *SPRY1* | 1.64 | 1.46 | 0.66 |

| | | | | |
|---|---|---|---|---|
| 674 | CHCHD2 | 1.64 | 1.46 | 0.66 |
| 675 | *AKT1* | 1.64 | 1.46 | 0.66 |
| 676 | NDUFA6 | 1.64 | 1.46 | 0.66 |
| 677 | *SYNJ2BP* | 1.64 | 1.46 | 0.66 |
| 678 | FAM129A | 1.63 | 1.46 | 0.66 |
| 679 | *ZNF76* | 1.63 | 1.46 | 0.66 |
| 680 | NEU1 | 1.63 | 1.46 | 0.66 |
| 681 | *TSPAN9* | 1.63 | 1.46 | 0.66 |
| 682 | CPM | 1.63 | 1.46 | 0.66 |
| 683 | *ETV4* | 1.63 | 1.46 | 0.66 |
| 684 | NLRX1 | 1.63 | 1.46 | 0.66 |
| 685 | *BZRAP1* | 1.63 | 1.46 | 0.66 |
| 686 | INHBB | 1.63 | 1.46 | 0.66 |
| 687 | *RAMP1* | 1.63 | 1.46 | 0.66 |
| 688 | KRCC1 | 1.62 | 1.46 | 0.66 |
| 689 | *DHRS3* | 1.62 | 1.46 | 0.66 |
| 690 | RXRB | 1.62 | 1.46 | 0.66 |
| 691 | *PRDM4* | -3.57 | 0.45 | 0.00 |
| 692 | GTF2E2 | -3.39 | 0.47 | 0.00 |
| 693 | *TXNRD1* | -3.11 | 0.50 | 0.00 |
| 694 | DUSP4 | -3.08 | 0.50 | 0.00 |
| 695 | *LEPROTL1* | -2.98 | 0.51 | 0.00 |
| 696 | SNF1LK | -2.94 | 0.52 | 0.00 |
| 697 | *DDX21* | -2.92 | 0.52 | 0.00 |
| 698 | KCTD9 | -2.88 | 0.52 | 0.00 |
| 699 | *TNFRSF10B* | -2.83 | 0.53 | 0.00 |
| 700 | CHMP7 | -2.81 | 0.53 | 0.00 |
| 701 | *PPP2R2A* | -2.81 | 0.53 | 0.00 |
| 702 | PAPD1 | -2.78 | 0.53 | 0.00 |
| 703 | *ZCCHC2* | -2.76 | 0.54 | 0.00 |
| 704 | RC3H2 | -2.74 | 0.54 | 0.00 |
| 705 | *DCTN6* | -2.73 | 0.54 | 0.00 |
| 706 | PPP1CC | -2.72 | 0.54 | 0.00 |
| 707 | *KIAA1033* | -2.70 | 0.54 | 0.00 |
| 708 | RPL7A | -2.69 | 0.54 | 0.00 |
| 709 | *RPLP0* | -2.68 | 0.54 | 0.00 |
| 710 | ID1 | -2.67 | 0.54 | 0.00 |
| 711 | *ENTPD4* | -2.63 | 0.55 | 0.00 |
| 712 | GCH1 | -2.61 | 0.55 | 0.00 |
| 713 | *RHOQ* | -2.61 | 0.55 | 0.00 |
| 714 | STK24 | -2.60 | 0.55 | 0.00 |
| 715 | *HRB* | -2.55 | 0.56 | 0.00 |
| 716 | DYNC1LI1 | -2.54 | 0.56 | 0.00 |
| 717 | *KIAA0020* | -2.54 | 0.56 | 0.00 |
| 718 | PTCD3 | -2.54 | 0.56 | 0.00 |

| | | | | |
|---|---|---|---|---|
| 719 | *MEMO1* | -2.54 | 0.56 | 0.00 |
| 720 | LSM1 | -2.54 | 0.56 | 0.00 |
| 721 | *WRN* | -2.54 | 0.56 | 0.00 |
| 722 | TDG | -2.53 | 0.56 | 0.00 |
| 723 | *ERLIN1* | -2.49 | 0.57 | 0.00 |
| 724 | UBE2E1 | -2.47 | 0.57 | 0.00 |
| 725 | *AGPAT5* | -2.46 | 0.57 | 0.00 |
| 726 | GNL3 | -2.46 | 0.57 | 0.00 |
| 727 | *ARPP-19* | -2.46 | 0.57 | 0.00 |
| 728 | STAM | -2.44 | 0.57 | 0.00 |
| 729 | *CYB5R4* | -2.43 | 0.57 | 0.00 |
| 730 | EIF3A | -2.43 | 0.57 | 0.00 |
| 731 | *ORC2L* | -2.41 | 0.58 | 0.00 |
| 732 | ISG20 | -2.41 | 0.58 | 0.00 |
| 733 | *FGG* | -2.40 | 0.58 | 0.00 |
| 734 | BAG1 | -2.40 | 0.58 | 0.00 |
| 735 | *CHUK* | -2.39 | 0.58 | 0.00 |
| 736 | PDSS1 | -2.39 | 0.58 | 0.00 |
| 737 | *INTS6* | -2.39 | 0.58 | 0.00 |
| 738 | RAB35 | -2.39 | 0.58 | 0.00 |
| 739 | *RPS6* | -2.36 | 0.58 | 0.00 |
| 740 | CNOT7 | -2.35 | 0.58 | 0.00 |
| 741 | *PTP4A1* | -2.34 | 0.59 | 0.00 |
| 742 | C8orf41 | -2.34 | 0.59 | 0.00 |
| 743 | *UBXD6* | -2.32 | 0.59 | 0.00 |
| 744 | URM1 | -2.32 | 0.59 | 0.00 |
| 745 | *MUC4* | -2.32 | 0.59 | 0.00 |
| 746 | IRS2 | -2.31 | 0.59 | 0.00 |
| 747 | *MRPS2* | -2.31 | 0.59 | 0.00 |
| 748 | RFK | -2.30 | 0.59 | 0.00 |
| 749 | *NOLA3* | -2.29 | 0.59 | 0.00 |
| 750 | TEX10 | -2.29 | 0.59 | 0.00 |
| 751 | *SLC39A14* | -2.28 | 0.59 | 0.00 |
| 752 | KIF5B | -2.28 | 0.59 | 0.00 |
| 753 | *EIF3J* | -2.27 | 0.59 | 0.00 |
| 754 | DDHD2 | -2.26 | 0.60 | 0.00 |
| 755 | *EIF4A1* | -2.26 | 0.60 | 0.00 |
| 756 | AVPI1 | -2.25 | 0.60 | 0.00 |
| 757 | *DNM1L* | -2.25 | 0.60 | 0.13 |
| 758 | NR4A2 | -2.24 | 0.60 | 0.13 |
| 759 | *ARHGEF10* | -2.22 | 0.60 | 0.13 |
| 760 | FBXL2 | -2.22 | 0.60 | 0.13 |
| 761 | *SEC61B* | -2.22 | 0.60 | 0.13 |
| 762 | SHOC2 | -2.22 | 0.60 | 0.13 |
| 763 | *SOCS6* | -2.21 | 0.60 | 0.13 |

| | | | | |
|---|---|---|---|---|
| 764 | UTP3 | -2.21 | 0.60 | 0.13 |
| 765 | *RBM13* | -2.21 | 0.60 | 0.13 |
| 766 | POLR1D | -2.21 | 0.60 | 0.13 |
| 767 | *KIAA0368* | -2.21 | 0.60 | 0.13 |
| 768 | RANBP5 | -2.21 | 0.60 | 0.13 |
| 769 | *CHMP1B* | -2.20 | 0.60 | 0.13 |
| 770 | SNRPD1 | -2.20 | 0.60 | 0.13 |
| 771 | *KLHL9* | -2.20 | 0.60 | 0.13 |
| 772 | C18orf8 | -2.20 | 0.60 | 0.13 |
| 773 | *ACTR3* | -2.20 | 0.60 | 0.13 |
| 774 | MYC | -2.18 | 0.61 | 0.13 |
| 775 | *KRAS* | -2.18 | 0.61 | 0.13 |
| 776 | ADAM10 | -2.17 | 0.61 | 0.13 |
| 777 | *MAPKAPK5* | -2.17 | 0.61 | 0.13 |
| 778 | PSMA4 | -2.17 | 0.61 | 0.13 |
| 779 | *SLC7A11* | -2.17 | 0.61 | 0.13 |
| 780 | GOLGA7 | -2.16 | 0.61 | 0.13 |
| 781 | *PPIF* | -2.16 | 0.61 | 0.13 |
| 782 | SETX | -2.16 | 0.61 | 0.13 |
| 783 | *KCMF1* | -2.16 | 0.61 | 0.13 |
| 784 | NARS | -2.15 | 0.61 | 0.13 |
| 785 | *RAP2A* | -2.15 | 0.61 | 0.13 |
| 786 | GPX2 | -2.15 | 0.61 | 0.13 |
| 787 | *SRP72* | -2.14 | 0.61 | 0.13 |
| 788 | CTSB | -2.14 | 0.61 | 0.13 |
| 789 | *TTLL4* | -2.14 | 0.61 | 0.13 |
| 790 | PIK3C3 | -2.14 | 0.61 | 0.13 |
| 791 | *RPL35* | -2.13 | 0.61 | 0.13 |
| 792 | KPNA3 | -2.13 | 0.61 | 0.13 |
| 793 | *BIN3* | -2.13 | 0.61 | 0.13 |
| 794 | AGTPBP1 | -2.13 | 0.61 | 0.30 |
| 795 | *PTPN11* | -2.13 | 0.61 | 0.30 |
| 796 | UBAC1 | -2.13 | 0.61 | 0.30 |
| 797 | *GSR* | -2.12 | 0.61 | 0.30 |
| 798 | EHBP1 | -2.12 | 0.61 | 0.30 |
| 799 | *RPL29* | -2.12 | 0.61 | 0.30 |
| 800 | MTHFD2 | -2.12 | 0.61 | 0.30 |
| 801 | *SOD2* | -2.12 | 0.61 | 0.30 |
| 802 | SET | -2.12 | 0.61 | 0.30 |
| 803 | *HMOX1* | -2.11 | 0.62 | 0.30 |
| 804 | VDAC3 | -2.10 | 0.62 | 0.30 |
| 805 | *VDAC2* | -2.10 | 0.62 | 0.30 |
| 806 | ZMYM2 | -2.10 | 0.62 | 0.30 |
| 807 | *GABPB2* | -2.09 | 0.62 | 0.30 |
| 808 | RPL6 | -2.09 | 0.62 | 0.30 |

| | | | | |
|---|---|---|---|---|
| 809 | *MEIS2* | -2.09 | 0.62 | 0.30 |
| 810 | PPP3CC | -2.09 | 0.62 | 0.30 |
| 811 | *SMAD2* | -2.08 | 0.62 | 0.30 |
| 812 | COPS2 | -2.08 | 0.62 | 0.30 |
| 813 | *TMF1* | -2.08 | 0.62 | 0.30 |
| 814 | TXNL4A | -2.07 | 0.62 | 0.30 |
| 815 | *XPO7* | -2.07 | 0.62 | 0.30 |
| 816 | PPP2R1B | -2.07 | 0.62 | 0.30 |
| 817 | *ODC1* | -2.07 | 0.62 | 0.30 |
| 818 | SCARB1 | -2.06 | 0.62 | 0.30 |
| 819 | *MRPS35* | -2.06 | 0.62 | 0.30 |
| 820 | POLR3D | -2.06 | 0.62 | 0.30 |
| 821 | *NDUFV2* | -2.06 | 0.62 | 0.30 |
| 822 | HYPK | -2.06 | 0.62 | 0.30 |
| 823 | *IKBKAP* | -2.06 | 0.62 | 0.30 |
| 824 | STX2 | -2.05 | 0.62 | 0.30 |
| 825 | *CHD7* | -2.05 | 0.62 | 0.30 |
| 826 | MAPK6 | -2.05 | 0.62 | 0.30 |
| 827 | *RAN* | -2.05 | 0.62 | 0.30 |
| 828 | MFHAS1 | -2.05 | 0.62 | 0.30 |
| 829 | *PDE4D* | -2.05 | 0.62 | 0.30 |
| 830 | RPS17 | -2.04 | 0.62 | 0.30 |
| 831 | *GTPBP4* | -2.04 | 0.62 | 0.30 |
| 832 | PWP1 | -2.04 | 0.62 | 0.30 |
| 833 | *RCBTB1* | -2.04 | 0.63 | 0.30 |
| 834 | ITGB1 | -2.03 | 0.63 | 0.30 |
| 835 | *KIAA0157* | -2.03 | 0.63 | 0.30 |
| 836 | SLC16A3 | -2.03 | 0.63 | 0.30 |
| 837 | *GSTO1* | -2.03 | 0.63 | 0.30 |
| 838 | PDHB | -2.03 | 0.63 | 0.30 |
| 839 | *KYNU* | -2.03 | 0.63 | 0.30 |
| 840 | CTDP1 | -2.02 | 0.63 | 0.30 |
| 841 | *VDAC1* | -2.02 | 0.63 | 0.30 |
| 842 | GALK2 | -2.02 | 0.63 | 0.30 |
| 843 | *PLEKHJ1* | -2.02 | 0.63 | 0.30 |
| 844 | C8orf4 | -2.02 | 0.63 | 0.30 |
| 845 | *C4orf16* | -2.02 | 0.63 | 0.30 |
| 846 | TMEM93 | -2.02 | 0.63 | 0.30 |
| 847 | *PUS1* | -2.02 | 0.63 | 0.30 |
| 848 | VPS33B | -2.02 | 0.63 | 0.30 |
| 849 | *UCK2* | -2.02 | 0.63 | 0.30 |
| 850 | PLAUR | -2.02 | 0.63 | 0.30 |
| 851 | *CEBPB* | -2.01 | 0.63 | 0.30 |
| 852 | CCDC59 | -2.01 | 0.63 | 0.30 |
| 853 | *UBE2N* | -2.01 | 0.63 | 0.30 |

| | | | | |
|---|---|---|---|---|
| 854 | SGCB | -2.01 | 0.63 | 0.30 |
| 855 | *PPP6C* | -2.00 | 0.63 | 0.30 |
| 856 | VAPA | -2.00 | 0.63 | 0.30 |
| 857 | *RTF1* | -2.00 | 0.63 | 0.30 |
| 858 | FAM60A | -2.00 | 0.63 | 0.30 |
| 859 | *C10orf22* | -2.00 | 0.63 | 0.30 |
| 860 | RPS7 | -2.00 | 0.63 | 0.30 |
| 861 | *RPS3* | -2.00 | 0.63 | 0.30 |
| 862 | PLAA | -2.00 | 0.63 | 0.30 |
| 863 | *ATF4* | -1.99 | 0.63 | 0.30 |
| 864 | PPP1R12A | -1.99 | 0.63 | 0.30 |
| 865 | *RY1* | -1.99 | 0.63 | 0.30 |
| 866 | MTMR9 | -1.99 | 0.63 | 0.30 |
| 867 | *YME1L1* | -1.99 | 0.63 | 0.30 |
| 868 | GTF3A | -1.99 | 0.63 | 0.30 |
| 869 | *C5orf30* | -1.98 | 0.63 | 0.30 |
| 870 | PMPCA | -1.98 | 0.63 | 0.30 |
| 871 | *TATDN2* | -1.98 | 0.63 | 0.30 |
| 872 | NRBF2 | -1.98 | 0.63 | 0.40 |
| 873 | *WIPI1* | -1.98 | 0.63 | 0.40 |
| 874 | CLDND1 | -1.98 | 0.63 | 0.40 |
| 875 | *SLC25A37* | -1.98 | 0.63 | 0.40 |
| 876 | HN1 | -1.97 | 0.63 | 0.40 |
| 877 | *FECH* | -1.97 | 0.64 | 0.40 |
| 878 | CTSL1 | -1.96 | 0.64 | 0.40 |
| 879 | *C12orf5* | -1.96 | 0.64 | 0.40 |
| 880 | FZD3 | -1.96 | 0.64 | 0.40 |
| 881 | *KCTD3* | -1.96 | 0.64 | 0.40 |
| 882 | TNFRSF1A | -1.96 | 0.64 | 0.40 |
| 883 | *MRPL42* | -1.96 | 0.64 | 0.40 |
| 884 | MBD2 | -1.96 | 0.64 | 0.40 |
| 885 | *SPCS3* | -1.96 | 0.64 | 0.40 |
| 886 | RPS19 | -1.96 | 0.64 | 0.40 |
| 887 | *RNF34* | -1.95 | 0.64 | 0.40 |
| 888 | S100P | -1.95 | 0.64 | 0.40 |
| 889 | *CYP24A1* | -1.95 | 0.64 | 0.40 |
| 890 | GZMB | -1.95 | 0.64 | 0.40 |
| 891 | *PPP4R2* | -1.95 | 0.64 | 0.40 |
| 892 | C12orf29 | -1.94 | 0.64 | 0.40 |
| 893 | *CLPX* | -1.94 | 0.64 | 0.40 |
| 894 | ACTR6 | -1.94 | 0.64 | 0.40 |
| 895 | *PFKP* | -1.94 | 0.64 | 0.40 |
| 896 | ASH2L | -1.94 | 0.64 | 0.40 |
| 897 | *EMR2* | -1.94 | 0.64 | 0.40 |
| 898 | CCT7 | -1.94 | 0.64 | 0.40 |

| | | | | |
|---|---|---|---|---|
| 899 | *CUL2* | -1.94 | 0.64 | 0.40 |
| 900 | GLCE | -1.94 | 0.64 | 0.40 |
| 901 | *AVEN* | -1.93 | 0.64 | 0.40 |
| 902 | SART3 | -1.93 | 0.64 | 0.40 |
| 903 | *WASF1* | -1.93 | 0.64 | 0.40 |
| 904 | RPL13A | -1.93 | 0.64 | 0.40 |
| 905 | *KIAA0701* | -1.93 | 0.64 | 0.40 |
| 906 | C15orf15 | -1.93 | 0.64 | 0.40 |
| 907 | *ROD1* | -1.93 | 0.64 | 0.40 |
| 908 | LTF | -1.92 | 0.64 | 0.40 |
| 909 | *ETNK1* | -1.92 | 0.64 | 0.40 |
| 910 | GCN1L1 | -1.92 | 0.64 | 0.40 |
| 911 | *PYROXD1* | -1.92 | 0.64 | 0.40 |
| 912 | WSB2 | -1.92 | 0.64 | 0.40 |
| 913 | *GRB2* | -1.92 | 0.64 | 0.40 |
| 914 | RPS6KB1 | -1.92 | 0.64 | 0.40 |
| 915 | *USP15* | -1.91 | 0.64 | 0.40 |
| 916 | STOML2 | -1.91 | 0.64 | 0.40 |
| 917 | *KIAA1012* | -1.91 | 0.64 | 0.40 |
| 918 | PTS | -1.91 | 0.64 | 0.40 |
| 919 | *RIOK3* | -1.91 | 0.64 | 0.40 |
| 920 | PDCD6IP | -1.91 | 0.64 | 0.40 |
| 921 | *C19orf21* | -1.90 | 0.64 | 0.40 |
| 922 | MAFF | -1.90 | 0.64 | 0.40 |
| 923 | *TRIM32* | -1.90 | 0.64 | 0.40 |
| 924 | ANKRD12 | -1.90 | 0.64 | 0.40 |
| 925 | *FAM20B* | -1.90 | 0.64 | 0.40 |
| 926 | CHST11 | -1.90 | 0.65 | 0.40 |
| 927 | *ATP6V0A2* | -1.90 | 0.65 | 0.40 |
| 928 | HERC2 | -1.89 | 0.65 | 0.40 |
| 929 | *SERPINB8* | -1.89 | 0.65 | 0.40 |
| 930 | TRIP12 | -1.89 | 0.65 | 0.40 |
| 931 | *GTF2A2* | -1.89 | 0.65 | 0.40 |
| 932 | SS18L2 | -1.88 | 0.65 | 0.40 |
| 933 | *PITRM1* | -1.88 | 0.65 | 0.40 |
| 934 | TMEM176A | -1.88 | 0.65 | 0.40 |
| 935 | *CAMSAP1* | -1.88 | 0.65 | 0.40 |
| 936 | MED27 | -1.88 | 0.65 | 0.40 |
| 937 | *PARD3* | -1.88 | 0.65 | 0.40 |
| 938 | PBK | -1.88 | 0.65 | 0.40 |
| 939 | *PLUNC* | -1.87 | 0.65 | 0.40 |
| 940 | NUP88 | -1.87 | 0.65 | 0.40 |
| 941 | *IL8* | -1.87 | 0.65 | 0.40 |
| 942 | ZCCHC6 | -1.86 | 0.65 | 0.40 |
| 943 | *TXNL1* | -1.86 | 0.65 | 0.40 |

| | | | | |
|---|---|---|---|---|
| 944 | POMP | -1.86 | 0.65 | 0.40 |
| 945 | *MAK10* | -1.85 | 0.65 | 0.40 |
| 946 | IL15RA | -1.85 | 0.65 | 0.40 |
| 947 | *MTMR2* | -1.85 | 0.65 | 0.40 |
| 948 | DDX54 | -1.85 | 0.65 | 0.40 |
| 949 | *XPNPEP1* | -1.85 | 0.65 | 0.66 |
| 950 | ECT2 | -1.85 | 0.65 | 0.66 |
| 951 | *CXCL2* | -1.85 | 0.65 | 0.66 |
| 952 | CDV3 | -1.85 | 0.65 | 0.66 |
| 953 | *BASP1* | -1.85 | 0.65 | 0.66 |
| 954 | CHML | -1.85 | 0.65 | 0.66 |
| 955 | *NOL8* | -1.85 | 0.65 | 0.66 |
| 956 | SLC7A1 | -1.85 | 0.65 | 0.66 |
| 957 | *LRFN4* | -1.84 | 0.65 | 0.66 |
| 958 | ISG20L1 | -1.84 | 0.65 | 0.66 |
| 959 | *HIP2* | -1.84 | 0.65 | 0.66 |
| 960 | IPPK | -1.84 | 0.65 | 0.66 |
| 961 | *RSRC2* | -1.84 | 0.65 | 0.66 |
| 962 | PCM1 | -1.84 | 0.65 | 0.66 |
| 963 | *RARRES1* | -1.84 | 0.65 | 0.66 |
| 964 | FBXO21 | -1.84 | 0.65 | 0.66 |
| 965 | *APPL2* | -1.84 | 0.65 | 0.66 |
| 966 | SCD | -1.83 | 0.65 | 0.66 |
| 967 | *COX5A* | -1.83 | 0.65 | 0.66 |
| 968 | TP53BP1 | -1.83 | 0.65 | 0.66 |
| 969 | *INTS7* | -1.83 | 0.65 | 0.66 |
| 970 | EDG2 | -1.83 | 0.65 | 0.66 |
| 971 | *UBE1C* | -1.83 | 0.65 | 0.66 |
| 972 | ZC3H15 | -1.83 | 0.65 | 0.66 |
| 973 | *C14orf161* | -1.83 | 0.66 | 0.66 |
| 974 | CHFR | -1.82 | 0.66 | 0.66 |
| 975 | *BUB1B* | -1.82 | 0.66 | 0.66 |
| 976 | OXSR1 | -1.82 | 0.66 | 0.66 |
| 977 | *NAB1* | -1.82 | 0.66 | 0.66 |
| 978 | AGPS | -1.82 | 0.66 | 0.66 |
| 979 | *TIMM17A* | -1.82 | 0.66 | 0.66 |
| 980 | TNFSF5IP1 | -1.82 | 0.66 | 0.66 |
| 981 | *PRMT1* | -1.82 | 0.66 | 0.66 |
| 982 | TEAD4 | -1.81 | 0.66 | 0.66 |
| 983 | *CDK7* | -1.81 | 0.66 | 0.66 |
| 984 | C19orf10 | -1.81 | 0.66 | 0.66 |
| 985 | *USP34* | -1.81 | 0.66 | 0.66 |
| 986 | PRPF4 | -1.81 | 0.66 | 0.66 |
| 987 | *IARS* | -1.81 | 0.66 | 0.66 |
| 988 | PSMD9 | -1.81 | 0.66 | 0.66 |

| | | | | |
|---|---|---|---|---|
| 989 | *GNLY* | -1.81 | 0.66 | 0.66 |
| 990 | FBXW2 | -1.80 | 0.66 | 0.66 |
| 991 | *UGDH* | -1.80 | 0.66 | 0.66 |
| 992 | VCP | -1.80 | 0.66 | 0.66 |
| 993 | *PRPF40A* | -1.80 | 0.66 | 0.66 |
| 994 | ADAMDEC1 | -1.80 | 0.66 | 0.66 |
| 995 | *P15RS* | -1.80 | 0.66 | 0.66 |
| 996 | ARL4C | -1.80 | 0.66 | 0.66 |
| 997 | *SEC16A* | -1.79 | 0.66 | 0.66 |
| 998 | CSDA | -1.79 | 0.66 | 0.66 |
| 999 | *PAK2* | -1.79 | 0.66 | 0.66 |
| 1000 | RIC8B | -1.79 | 0.66 | 0.66 |
| 1001 | *TMEM176B* | -1.78 | 0.66 | 0.66 |
| 1002 | STK4 | -1.78 | 0.66 | 0.66 |
| 1003 | *AGPAT7* | -1.78 | 0.66 | 0.66 |
| 1004 | IVNS1ABP | -1.78 | 0.66 | 0.66 |
| 1005 | *MED13L* | -1.78 | 0.66 | 0.66 |
| 1006 | BNIP2 | -1.77 | 0.66 | 0.66 |
| 1007 | *ENDOG* | -1.77 | 0.66 | 0.66 |
| 1008 | PDCL | -1.77 | 0.66 | 0.66 |
| 1009 | *MBD1* | -1.77 | 0.66 | 0.66 |
| 1010 | YARS2 | -1.77 | 0.66 | 0.66 |
| 1011 | *P2RX5* | -1.77 | 0.66 | 0.66 |
| 1012 | RABEPK | -1.77 | 0.66 | 0.66 |
| 1013 | *CLN8* | -1.77 | 0.66 | 0.66 |
| 1014 | ENTPD7 | -1.76 | 0.66 | 0.66 |
| 1015 | *RPS27A* | -1.76 | 0.66 | 0.66 |
| 1016 | SPATS2 | -1.76 | 0.67 | 0.66 |
| 1017 | *INTS9* | -1.76 | 0.67 | 0.66 |
| 1018 | KLHL18 | -1.76 | 0.67 | 0.66 |
| 1019 | *F3* | -1.76 | 0.67 | 0.66 |
| 1020 | JMJD6 | -1.76 | 0.67 | 0.66 |
