## Supplementary Table 2 for "Lung Adenocarcinoma Just Desserts: An Expanding Pie of Activating Oncogenes or a Layer Cake of Integrated Alterations"

**Supplementary Table 2. Performance test results between EGFR-like mutant group and WT group based on EGFR-related gene signature.** mt, mutant, WT, wild-type, PPV, positive predictive value, NPV, negative predictive value.

| Dataset | Cohort | EGFR mSig | EGFR Status | | Total | Performance | | | | |
|---|---|---|---|---|---|---|---|---|---|---|
| | | | mt | WT | | Sensitivity | Specificity | Accuracy | PPV | NPV |
| Training | MSKCC | + | 35 | 25 | 60 | 0.90 | 0.84 | 0.85 | 0.58 | 0.97 |
| | | - | 4 | 128 | 132 | | | | | |
| | | Total | 39 | 153 | 192 | | | | | |
| Validation | UNC+TSP | + | 13 | 10 | 23 | 0.72 | 0.90 | 0.87 | 0.57 | 0.95 |
| | | - | 5 | 86 | 91 | | | | | |
| | | Total | 18 | 96 | 114 | | | | | |
| Validation | TCGA | + | 52 | 54 | 106 | 0.81 | 0.87 | 0.86 | 0.49 | 0.97 |
| | | - | 12 | 368 | 380 | | | | | |
| | | Total | 64 | 422 | 486 | | | | | |
| Average | | | | | | 0.81 | 0.87 | 0.86 | 0.55 | 0.96 |
