## Supplementary Table 3 for "Lung Adenocarcinoma Just Desserts: An Expanding Pie of Activating Oncogenes or a Layer Cake of Integrated Alterations"

**Supplementary Table S3**. Odds Ratio and p-value; Subtype vs. Non-Subtypes.

| ***Odds Ratio*** | **EGFR mt** | **EGFR mSig** | **EGFR cn** | **TP53 mt** | **KEAP1/STK11 mt** | **RAS/RAF/RTK mt** | **EGFR ge** | **p16/CDKN2A ge** | **NKX2-1 ge** | **TP63 ge** | **DUSP4 ge** |
|---|---|---|---|---|---|---|---|---|---|---|---|
| *Bronchioid* | 4.48 | 9.18 | 0.50 | 0.26 | 0.35 | 1.44 | 1.46 | 0.71 | 4.08 | 2.73 | 0.43 |
| *Magnoid* | 0.17 | 0.09 | 0.91 | 0.89 | 8.09 | 0.90 | 0.31 | 0.66 | 0.87 | 0.20 | 5.52 |
| *Squamoid* | 0.59 | 0.39 | 2.23 | 4.65 | 0.32 | 0.77 | 1.94 | 2.09 | 0.28 | 1.44 | 0.56 |
| *EGFR mt* | NA | 29.18 | 6.89 | 0.97 | 0.07 | 0.01 | 8.76 | 0.93 | 1.55 | 1.80 | 0.05 |
| *EGFR mSig* | NA | NA | 2.45 | 0.83 | 0.11 | 0.43 | 4.08 | 1.16 | 3.84 | 1.99 | 0.08 |
| *EGFR cn* | NA | NA | NA | 2.67 | 0.60 | 0.45 | 3.10 | 1.37 | 0.62 | 1.28 | 0.49 |
| *TP53 mt* | NA | NA | NA | NA | 0.66 | 0.36 | 1.62 | 2.27 | 0.58 | 0.89 | 0.60 |
| *KEAP1/STK11 mt* | NA | NA | NA | NA | NA | 1.11 | 0.21 | 0.76 | 0.92 | 0.29 | 5.73 |
| *RAS/RAF/RTK mt* | NA | NA | NA | NA | NA | NA | 0.72 | 0.67 | 1.81 | 1.03 | 1.75 |
| *EGFR ge* | NA | NA | NA | NA | NA | NA | NA | 1.05 | 1.37 | 2.11 | 0.22 |
| *p16/CDKN2A ge* | NA | NA | NA | NA | NA | NA | NA | NA | 0.60 | 1.12 | 0.68 |
| *NKX2-1 ge* | NA | NA | NA | NA | NA | NA | NA | NA | NA | 0.76 | 0.40 |
| *TP63 ge* | NA | NA | NA | NA | NA | NA | NA | NA | NA | NA | 0.58 |
| *DUSP4 ge* | NA | NA | NA | NA | NA | NA | NA | NA | NA | NA | NA |

| ***P-values*** | **EGFR mt** | **EGFR mSig** | **EGFR cn** | **TP53 mt** | **KEAP1/STK11 mt** | **RAS/RAF/RTK mt** | **EGFR ge** | **p16/CDKN2A ge** | **NKX2-1 ge** | **TP63 ge** | **DUSP4 ge** |
|---|---|---|---|---|---|---|---|---|---|---|---|
| *Bronchioid* | 8.97E-08 | 4.29E-21 | 3.27E-04 | 1.06E-11 | 2.73E-06 | 5.92E-02 | 5.86E-02 | 8.88E-02 | 1.50E-12 | 2.85E-07 | 1.27E-05 |
| *Magnoid* | 1.32E-05 | 2.96E-12 | 6.90E-01 | 6.18E-01 | 5.33E-22 | 6.19E-01 | 1.81E-08 | 4.63E-02 | 5.50E-01 | 6.31E-14 | 2.28E-15 |
| *Squamoid* | 0.092 | 0.000 | 0.000 | 0.000 | 0.000 | 0.182 | 0.001 | 0.000 | 0.000 | 0.070 | 0.003 |
| *EGFR mt* | NA | 0.000 | 0.000 | 1.000 | 0.000 | 0.000 | 0.000 | 0.893 | 0.140 | 0.044 | 0.000 |
| *EGFR mSig* | NA | NA | 0.000 | 0.441 | 0.000 | 0.000 | 0.000 | 0.583 | 0.000 | 0.003 | 0.000 |
| *EGFR cn* | NA | NA | NA | 0.000 | 0.012 | 0.000 | 0.000 | 0.102 | 0.011 | 0.204 | 0.000 |
| *TP53 mt* | NA | NA | NA | NA | 0.046 | 0.000 | 0.011 | 0.000 | 0.004 | 0.585 | 0.006 |
| *KEAP1/STK11 mt* | NA | NA | NA | NA | NA | 0.619 | 0.000 | 0.195 | 0.765 | 0.000 | 0.000 |
| *RAS/RAF/RTK mt* | NA | NA | NA | NA | NA | NA | 0.085 | 0.037 | 0.001 | 0.928 | 0.003 |
| *EGFR ge* | NA | NA | NA | NA | NA | NA | NA | 0.856 | 0.102 | 0.000 | 0.000 |
| *p16/CDKN2A ge* | NA | NA | NA | NA | NA | NA | NA | NA | 0.006 | 0.586 | 0.046 |
| *NKX2-1 ge* | NA | NA | NA | NA | NA | NA | NA | NA | NA | 0.147 | 0.000 |
| *TP63 ge* | NA | NA | NA | NA | NA | NA | NA | NA | NA | NA | 0.004 |
| *DUSP4 ge* | NA | NA | NA | NA | NA | NA | NA | NA | NA | NA | NA |
