## Supplementary Table 4 for "Lung Adenocarcinoma Just Desserts: An Expanding Pie of Activating Oncogenes or a Layer Cake of Integrated Alterations"

**Supplementary Table S4**. Odds Ratio and p-value; Within Subtype. mt, mutant, cn, copy number, ge, gene expression. Inf, infinite value due to zero count.

***Bronchioid***

| *Odds Ratio* | EGFR mt | EGFR mSig | EGFR cn | TP53 mt | KEAP1/STK11 mt | RAS/RAF/RTK mt | EGFR ge | p16/CDKN2A ge | NKX2-1 ge | TP63 ge | DUSP4 ge |
|---|---|---|---|---|---|---|---|---|---|---|---|
| *EGFR mt* | NA | 47.81 | Inf | 1.65 | 0.09 | 0.01 | 9.67 | 0.72 | 0.93 | 1.34 | 0.15 |
| *EGFR mSig* | NA | NA | 7.20 | 1.96 | 0.20 | 0.24 | 5.65 | 1.70 | 2.05 | 1.18 | 0.15 |
| *EGFR cn* | NA | NA | NA | 4.16 | 0.22 | 0.15 | 3.74 | 1.28 | 0.49 | 3.74 | 0.49 |
| *TP53 mt* | NA | NA | NA | NA | 0.73 | 0.51 | 2.04 | 1.34 | 0.59 | 1.34 | 0.89 |
| *KEAP1/STK11 mt* | NA | NA | NA | NA | NA | 1.61 | 0.08 | 1.26 | 2.54 | 0.21 | 4.74 |
| *RAS/RAF/RTK mt* | NA | NA | NA | NA | NA | NA | 0.31 | 1.55 | 1.55 | 0.97 | 2.26 |
| *EGFR ge* | NA | NA | NA | NA | NA | NA | NA | 1.41 | 1.41 | 1.54 | 0.19 |
| *p16/CDKN2A ge* | NA | NA | NA | NA | NA | NA | NA | NA | 2.04 | 1.07 | 0.68 |
| *NKX2-1 ge* | NA | NA | NA | NA | NA | NA | NA | NA | NA | 0.68 | 0.51 |
| *TP63 ge* | NA | NA | NA | NA | NA | NA | NA | NA | NA | NA | 0.74 |
| *DUSP4 ge* | NA | NA | NA | NA | NA | NA | NA | NA | NA | NA | NA |

| *P-values* | EGFR mt | EGFR mSig | EGFR cn | TP53 mt | KEAP1/STK11 mt | RAS/RAF/RTK mt | EGFR ge | p16/CDKN2A ge | NKX2-1 ge | TP63 ge | DUSP4 ge |
|---|---|---|---|---|---|---|---|---|---|---|---|
| *EGFR mt* | NA | 2.87E-15 | 1.15E-01 | 1.93E-01 | 2.01E-03 | 5.94E-18 | 4.72E-08 | 3.84E-01 | 8.62E-01 | 4.83E-01 | 1.38E-06 |
| *EGFR mSig* | NA | NA | 4.03E-02 | 5.26E-02 | 8.70E-04 | 7.97E-06 | 7.73E-08 | 9.56E-02 | 2.29E-02 | 6.50E-01 | 5.12E-09 |
| *EGFR cn* | NA | NA | NA | 2.75E-01 | 4.26E-02 | 7.95E-02 | 9.93E-02 | 7.47E-01 | 4.96E-01 | 9.93E-02 | 4.96E-01 |
| *TP53 mt* | NA | NA | NA | NA | 5.26E-01 | 5.18E-02 | 3.66E-02 | 4.23E-01 | 1.10E-01 | 4.23E-01 | 7.50E-01 |
| *KEAP1/STK11 mt* | NA | NA | NA | NA | NA | 3.09E-01 | 1.25E-06 | 6.85E-01 | 4.08E-02 | 4.89E-04 | 8.94E-04 |
| *RAS/RAF/RTK mt* | NA | NA | NA | NA | NA | NA | 2.43E-04 | 1.72E-01 | 1.72E-01 | 1.00E+00 | 9.68E-03 |
| *EGFR ge* | NA | NA | NA | NA | NA | NA | NA | 2.91E-01 | 2.91E-01 | 1.75E-01 | 2.09E-07 |
| *p16/CDKN2A ge* | NA | NA | NA | NA | NA | NA | NA | NA | 2.35E-02 | 8.80E-01 | 2.28E-01 |
| *NKX2-1 ge* | NA | NA | NA | NA | NA | NA | NA | NA | NA | 2.28E-01 | 3.45E-02 |
| *TP63 ge* | NA | NA | NA | NA | NA | NA | NA | NA | NA | NA | 3.66E-01 |
| *DUSP4 ge* | NA | NA | NA | NA | NA | NA | NA | NA | NA | NA | NA |

***Magnoid***

| *Odds Ratio* | EGFR mt | EGFR mSig | EGFR cn | TP53 mt | KEAP1/STK11 mt | RAS/RAF/RTK mt | EGFR ge | p16/CDKN2A ge | NKX2-1 ge | TP63 ge | DUSP4 ge |
|---|---|---|---|---|---|---|---|---|---|---|---|
| *EGFR mt* | NA | 8.07 | Inf | 3.85 | 0.00 | 0.00 | 4.08 | 4.08 | 1.50 | 4.08 | 0.00 |
| *EGFR mSig* | NA | NA | 4.08 | Inf | 0.00 | 0.26 | 4.08 | 4.08 | 1.50 | Inf | 0.00 |
| *EGFR cn* | NA | NA | NA | 2.70 | 0.72 | 0.53 | 3.04 | 0.97 | 0.50 | 1.36 | 0.62 |
| *TP53 mt* | NA | NA | NA | NA | 0.34 | 0.15 | 2.40 | 5.00 | 0.49 | 1.09 | 0.24 |
| *KEAP1/STK11 mt* | NA | NA | NA | NA | NA | 2.61 | 0.50 | 0.64 | 0.72 | 0.81 | 2.66 |
| *RAS/RAF/RTK mt* | NA | NA | NA | NA | NA | NA | 1.03 | 0.33 | 1.44 | 0.92 | 6.13 |
| *EGFR ge* | NA | NA | NA | NA | NA | NA | NA | 0.97 | 0.78 | 1.52 | 0.97 |
| *p16/CDKN2A ge* | NA | NA | NA | NA | NA | NA | NA | NA | 0.56 | 1.36 | 0.39 |
| *NKX2-1 ge* | NA | NA | NA | NA | NA | NA | NA | NA | NA | 0.56 | 0.56 |
| *TP63 ge* | NA | NA | NA | NA | NA | NA | NA | NA | NA | NA | 0.62 |
| *DUSP4 ge* | NA | NA | NA | NA | NA | NA | NA | NA | NA | NA | NA |

| *P-values* | EGFR mt | EGFR mSig | EGFR cn | TP53 mt | KEAP1/STK11 mt | RAS/RAF/RTK mt | EGFR ge | p16/CDKN2A ge | NKX2-1 ge | TP63 ge | DUSP4 ge |
|---|---|---|---|---|---|---|---|---|---|---|---|
| *EGFR mt* | NA | 0.17 | 0.06 | 0.37 | 0.01 | 0.06 | 0.37 | 0.37 | 1.00 | 0.37 | 0.03 |
| *EGFR mSig* | NA | NA | 0.37 | 0.06 | 0.01 | 0.37 | 0.37 | 0.37 | 1.00 | 0.06 | 0.03 |
| *EGFR cn* | NA | NA | NA | 0.00 | 0.39 | 0.07 | 0.00 | 1.00 | 0.05 | 0.40 | 0.18 |
| *TP53 mt* | NA | NA | NA | NA | 0.00 | 0.00 | 0.01 | 0.00 | 0.04 | 0.87 | 0.00 |
| *KEAP1/STK11 mt* | NA | NA | NA | NA | NA | 0.01 | 0.06 | 0.23 | 0.39 | 0.61 | 0.01 |
| *RAS/RAF/RTK mt* | NA | NA | NA | NA | NA | NA | 1.00 | 0.00 | 0.32 | 0.87 | 0.00 |
| *EGFR ge* | NA | NA | NA | NA | NA | NA | NA | 1.00 | 0.51 | 0.24 | 1.00 |
| *p16/CDKN2A ge* | NA | NA | NA | NA | NA | NA | NA | NA | 0.10 | 0.40 | 0.01 |
| *NKX2-1 ge* | NA | NA | NA | NA | NA | NA | NA | NA | NA | 0.10 | 0.10 |
| *TP63 ge* | NA | NA | NA | NA | NA | NA | NA | NA | NA | NA | 0.18 |
| *DUSP4 ge* | NA | NA | NA | NA | NA | NA | NA | NA | NA | NA | NA |

***Squamoid***

| *Odds Ratio* | EGFR mt | EGFR mSig | EGFR cn | TP53 mt | KEAP1/STK11 mt | RAS/RAF/RTK mt | EGFR ge | p16/CDKN2A ge | NKX2-1 ge | TP63 ge | DUSP4 ge |
|---|---|---|---|---|---|---|---|---|---|---|---|
| *EGFR mt* | NA | 20.53 | 2.52 | 0.90 | 0.34 | 0.00 | 17.82 | 1.32 | 2.37 | 1.32 | 0.57 |
| *EGFR mSig* | NA | NA | 1.85 | 0.73 | 0.25 | 0.33 | 7.29 | 1.11 | 2.81 | 1.11 | 0.27 |
| *EGFR cn* | NA | NA | NA | 1.21 | 0.70 | 1.00 | 4.42 | 1.30 | 1.17 | 1.78 | 0.95 |
| *TP53 mt* | NA | NA | NA | NA | 1.79 | 0.58 | 2.11 | 2.82 | 1.60 | 0.55 | 0.35 |
| *KEAP1/STK11 mt* | NA | NA | NA | NA | NA | 0.23 | 0.69 | 1.74 | 0.31 | 0.83 | 1.74 |
| *RAS/RAF/RTK mt* | NA | NA | NA | NA | NA | NA | 1.05 | 0.65 | 1.87 | 0.87 | 1.40 |
| *EGFR ge* | NA | NA | NA | NA | NA | NA | NA | 0.75 | 1.95 | 1.46 | 0.62 |
| *p16/CDKN2A ge* | NA | NA | NA | NA | NA | NA | NA | NA | 0.51 | 1.21 | 1.46 |
| *NKX2-1 ge* | NA | NA | NA | NA | NA | NA | NA | NA | NA | 0.68 | 0.34 |
| *TP63 ge* | NA | NA | NA | NA | NA | NA | NA | NA | NA | NA | 1.46 |
| *DUSP4 ge* | NA | NA | NA | NA | NA | NA | NA | NA | NA | NA | NA |

| *P-values* | EGFR mt | EGFR mSig | EGFR cn | TP53 mt | KEAP1/STK11 mt | RAS/RAF/RTK mt | EGFR ge | p16/CDKN2A ge | NKX2-1 ge | TP63 ge | DUSP4 ge |
|---|---|---|---|---|---|---|---|---|---|---|---|
| *EGFR mt* | NA | 5.05E-07 | 1.78E-01 | 1.00E+00 | 4.71E-01 | 2.98E-05 | 2.84E-04 | 7.94E-01 | 1.88E-01 | 7.94E-01 | 4.31E-01 |
| *EGFR mSig* | NA | NA | 3.30E-01 | 5.82E-01 | 2.04E-01 | 3.58E-02 | 7.20E-04 | 1.00E+00 | 6.00E-02 | 1.00E+00 | 1.79E-02 |
| *EGFR cn* | NA | NA | NA | 7.02E-01 | 5.03E-01 | 1.00E+00 | 2.10E-05 | 5.18E-01 | 7.47E-01 | 1.06E-01 | 1.00E+00 |
| *TP53 mt* | NA | NA | NA | NA | 4.49E-01 | 1.45E-01 | 6.68E-02 | 9.95E-03 | 2.73E-01 | 1.43E-01 | 9.95E-03 |
| *KEAP1/STK11 mt* | NA | NA | NA | NA | NA | 4.65E-03 | 5.23E-01 | 2.86E-01 | 1.77E-02 | 8.31E-01 | 2.86E-01 |
| *RAS/RAF/RTK mt* | NA | NA | NA | NA | NA | NA | 1.00E+00 | 2.15E-01 | 6.28E-02 | 7.57E-01 | 3.53E-01 |
| *EGFR ge* | NA | NA | NA | NA | NA | NA | NA | 4.41E-01 | 4.45E-02 | 2.80E-01 | 1.65E-01 |
| *p16/CDKN2A ge* | NA | NA | NA | NA | NA | NA | NA | NA | 4.45E-02 | 6.44E-01 | 2.80E-01 |
| *NKX2-1 ge* | NA | NA | NA | NA | NA | NA | NA | NA | NA | 2.80E-01 | 1.13E-03 |
| *TP63 ge* | NA | NA | NA | NA | NA | NA | NA | NA | NA | NA | 2.80E-01 |
| *DUSP4 ge* | NA | NA | NA | NA | NA | NA | NA | NA | NA | NA | NA |
